# Evolution Inspired Engineering of Megasynthetases

**DOI:** 10.1101/2022.12.02.518901

**Authors:** Kenan A. J. Bozhüyük, Leonard Präve, Carsten Kegler, Sebastian Kaiser, Yan-Ni Shi, Wolfgang Kuttenlochner, Leonie Schenk, T. M. Mohiuddin, Michael Groll, Georg K. A. Hochberg, Helge B. Bode

## Abstract

Many clinically used drugs are derived from or inspired by bacterial natural products that often are biosynthesised via non-ribosomal peptide synthetases (NRPS), giant megasynthases that activate and join individual amino acids in an assembly line fashion. Since NRPS are not limited to the incorporation of the 20 proteinogenic amino acids, their efficient manipulation would allow the biotechnological generation of complex peptides including linear, cyclic and further modified natural product analogues, e.g. to optimise natural product leads. Here we describe a detailed phylogenetic analysis of several bacterial NRPS that led to the identification of a new recombination breakpoint within the thiolation (T) domain that is important for natural NRPS evolution. From this, an evolution-inspired eXchange Unit between T domains (XUT) approach was developed which allows the assembly of NRPS fragments over a broad range of GC contents, protein similarities, and extender unit specificities, as demonstrated for the specific production of a proteasome inhibitor designed and assembled from five different NRPS fragments.

## Introduction

Natural products (NPs) have been extensively studied for their therapeutic potential given their remarkable chemical and structural diversity in nature. Not only are they considered a rich reservoir of pharmacologically active lead compounds with therapeutic potential, but with ~48% of all new medicines approved between 1981 and 2019 originating in nature, NPs play an important role in the drug discovery and development process^1^. In recent decades, the collective efforts of the scientific community have led to tremendous progress in the identification of novel NPs to evaluate their pharmacological properties and mode of action, that could not be easily transferred into the development of new clinical drugs^2^. One of many reasons why the pharmaceutical industry stepped back from NP-based drug discovery.

Genetic engineering of natural products holds the potential for faster and more cost-effective discovery of (tailor-made) biological drugs than conventional methods^3^. Many bioactive bacterial NPs are derived from biosynthetic gene clusters (BGCs), genomic bacterial islands encoding non-ribosomal peptide synthetases (NRPS)^4^. NRPS are genetically encoded molecular assembly lines with many moving parts and reaction centres that all work together, to produce a broad variety of valuable non-ribosomal peptides (NRP) or even clinical drugs – such as penicillins^5–7^, bleomycin^8^, and ciclosporin^9^. Given these outstanding biological activities that benefit global public health, NRPS assembly lines would be an ideal target for synthetic biology, e.g. to improve pharmacological properties of natural product leads for (pre-) clinical development.

NRPS assembly lines consist of sequentially repeating modules of enzymatic domains, each of which catalyses the incorporation and chemical modification of a specific extender unit into the growing chain before the extended chain is passed on to the next module^4^. Hundreds of different extender units, typically derived from amino acids, have been described so far^10,11^. Selection and activation of an extender unit within an NRPS is catalysed by an adenylation (A) domain. The activated substrate is then covalently attached to the post-translationally attached prosthetic thiol (phosphopantetheine) group of a small thiolation (T) domain. Condensation (C) domains then link the covalently bound substrates to the growing NRP-chain in a co-linear fashion. In addition to these “core” domains that define the functional unit of an assembly line module, tailoring domains may be present to modify NRP chain, that are heterocyclization (Cy), epimerisation (E), *N*-methylation (MT), oxidation (Ox) or reduction (R) domains. Finally, the full-length NRP is released from the enzymatic machinery by hydrolysis or macrocyclization catalysed by a thioesterase (TE) domain.

The very logic of this assembly line mechanism inspired numerous rational efforts to engineer megasynthases to produce natural product analogues or even artificial NP-like compounds^12^. Although early engineering attempts yielded biosynthesis clusters that were either greatly impaired in their activity or non-functional, recent technical advances and the growing body of structural data accelerated the development of innovative synthetic biology strategies to engineer megasynthetases. Examples are the identification of interchangeable catalytically functional domain units^13,14^, CRISPR-Cas9 based gene editing to engineer complex antibiotic assembly lines^15^, yeast cell surface display assay to engineer the specificity of individual A domains,^16^ and splitting megasynthases into individually expressible subunits to reduce their complexity and size (up to several MDa) either *via* adding zinc-finger tags^17^ (DNA-templated NRPS) or SYNZIPs^18–20^ (heterospecific coiled-coil peptides). Furthermore, with the continuous increase in publicly available genomic data and the extensive efforts of the community to develop processing tools for BGC and NP identification^21–23^, there is a new trend towards assembly-line engineering using evolution-driven strategies. A number of insightful studies have led to the conclusion that understanding the mechanisms by which nature has evolved these often huge multifunctional enzyme machines will further improve our ability to redesign assembly line proteins to achieve even greater structural diversity while maintaining good production titres and could help us to expand our therapeutic arsenal^24–26^. However, the evolutionary mechanisms to achieve the exchange of individual extender units in NRP scaffolds are still poorly understood.

The genetic and architectural modularity of NRPS, but also of the biochemically distinct yet mechanistically analogous polyketide synthases (PKS), is central for current evolution models of these BGCs. Historically, the functional unit able to perform one round of chain elongation of a PKS (KS-AT-(DH-KR-ER)-T) and NRPS (C-A-T) is called a module. It is yet unclear whether this architectural and genetic unit also corresponds to an evolutionary unit that has been preserved in megasynthases^24,26^. Phylogenetic and computational analyses of the PKS family have led to a proposed redefinition of module boundaries from the “historical” KS-AT-(DH-KR-ER)-ACP to the “alternative” AT-(DH-KR-ER)-ACP-KS^27,28^, and highlighted the presence of genetic repeats^29^ (GRINS = genetic repeats of intense nucleotide skews) in a large number of PKS. The latter play a putative role in accelerating diversification of closely related BGCs by promoting gene conversion. For NRPS, studies on the underlying evolutionary processes have only just begun.

In a recent *in silico* study, the evolution of bacterial NRPS across various phyla was analysed^24^. The authors not only showed that intragenomic recombination along with speciation and horizontal gene transfer together with recombination are important factors in NRPS evolution, but also enabled the authors to introduce a unifying model for the evolution of the present-day variety of NRPs. Within the framework of this model, it was suggested that single recombination events at multiple breakpoints within the A domains of NRPS, referred to as subdomain swapping, are not only a widespread phenomenon, as reported previously^25,30–32^, but are also a major contributing factor to the diversification and functionalisation of NRP families. Further key findings are that stereochemical changes from the L- to the D-configuration in the final NRP seem to be achieved by the combined exchange of T-C di-for T-E-C tri-domains; and that there is a trend to keep intact both the native C-A linker region, physically connecting both domains, and the A-T domain interface. However, the practical evidence of these findings for successful NRPS engineering on a broad basis have not been shown yet. Up to date, there are only a very limited number of examples where evolutionary insights have been successfully used to engineer megasynthases – but only within a very narrow range of genetic and chemical changes introduced into the underlying BGCs and produced NPs, respectively^15,32,33^.

Herein, we particularly focused on deciphering the evolutionary history of NRPS to identify an evolution-inspired moiety that is best suited to enable NRPS engineering in a unified and more efficient manner. In order to approach the problem from different angles, a broad dataset of NRPS sequences from different phyla was analysed *in silico* to identify recombination events, a fusion point screening was performed to identify ideal engineering sites, the identified sites were broadly evaluated by reprogramming NRPS enzymes, and finally this knowledge was used to design a pharmaceutically active peptide *de novo*.

## Results

### Deciphering the Evolutionary History of NRPS

Homologous recombination is a pervasive biological process that affects sequences in all living organisms and undoubtedly is the main driver for megasynthase diversification^34^. Sequences having undergone recombination, such as NRPS, will display two different histories: one history for one part of their sequence, affected by the recombination event, and one history for the other part. Consequently, the evolutionary history of an alignment of homologous NRPS sequences cannot be properly depicted by classical phylogenetic methods because only one bifurcating tree is reconstructed. Therefore, we applied a previously established maximum likelihood method that was explicitly designed to detect multiple phylogenetic histories caused by recombination events. It uses a phylogenetic Hidden Markov Model (HMM) to search for a specified number of independent evolutionary histories that together best explain the alignment^34^. The algorithm returns the site likelihoods for each tree for every single position in an alignment, which can then be used to detect recombination breakpoints. In our analysis, we searched for two different histories, expecting one to broadly fit known A domain trees^35^, and the other to broadly fit known C domain phylogenies^36,37^. To identify which sites, belong to which history, we then subtracted the site-wise log likelihoods of the second tree from those of the first tree (Figs. 1a and Supplementary Fig. 1) – positive values indicate sites that are better described by the first phylogenetic history, sites with negative values are better described by the second phylogenetic history.

**Fig. 1.**
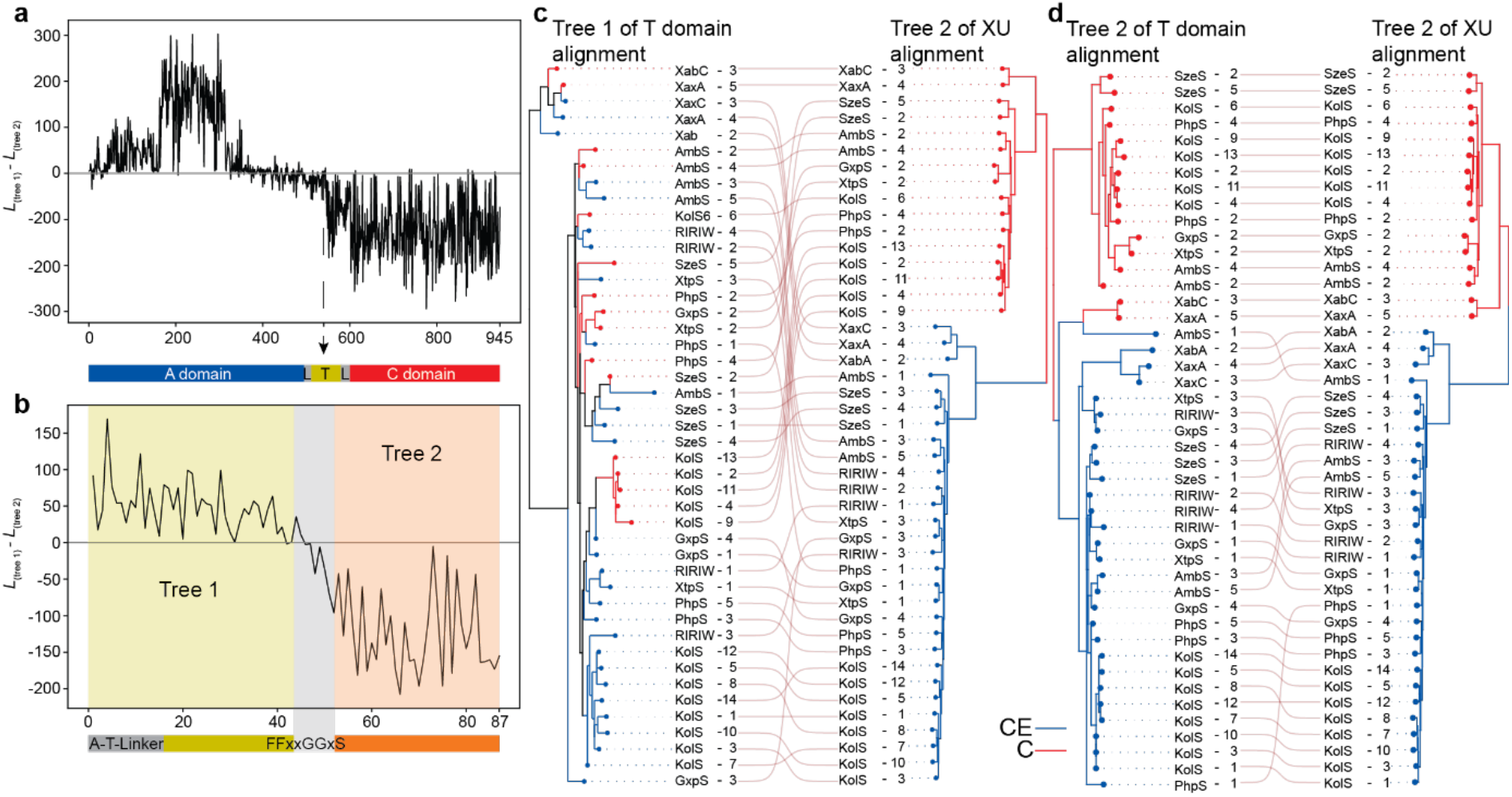
Evolutionary analysis of ATC tri-domains and T domains of representative NRPS. (**a**) Likelihood difference plot of two phylogenetic trees of ATC tri-domains (also called XUs) that together best describe the alignment using a phylogenetic hidden Markov model. Positive numbers indicate that sites are better describe by tree 1, negative numbers indicate sites that are better described by tree two. (**b**) Likelihood difference plot as in **a**, but for an alignment of T domain plus A-T linker. Partitions detected by the hidden Markov model are indicated in different colours according to tree number. Recombination breakpoint is annotated in grey and lies around two conserved glycine residues. (**c**, **d**) Comparison of Tree 1 from the T domain alignment with Tree 2 from the XU alignment (left) and Tree 2 from the T domain alignment with Tree 2 from the XU alignment (right). Names indicate abbreviation of NRPS and numbers the XU within that NRPS. Lines connect the same NRPS and XUs between the two trees. Red branches label XUs that contain ^L^CL domains, blue branches label XUs with C/E (dual C) domains.

We applied this method to a dataset comprising of >200 aligned amino acid sequences of NRPS A-T-C tri-domains from *Photorhabdus* and *Xenorhabdus* species, as well as representative NRPS from firmicutes, actinomycetes, cyanobacteria and other proteobacteria (Supplementary Dataset 1). This analysis revealed two major insights: First, sites in the A domain mostly preferred the first history, and sites in the C domain strongly prefer the second history (Supplementary Fig. 2). This confirms our method can detect that A and C domains have different evolutionary histories. And second, the breakpoint between these two histories appears to lie somewhere inside the T domain (Fig. 1a), though were exactly was not clear from this analysis: The difference in site likelihoods between histories becomes significantly more negative (indicating a preference for the second history) roughly in the middle of the T-domain, from around zero (not preferring either tree) to values well below −50 log units (strongly preferring the second tree).

To gain a better understanding of this potential recombination breakpoint, we repeated our phylogenetic HMM analysis with just the T domain together with the A-T-linker, again searching for two histories (Fig. 1b – d, and Supplementary Fig. 1; Supplementary Dataset 2). We did this, because the first half of the T domain preferred neither tree in our first analysis (Figs. 1c and Supplementary Fig. 3), potentially because it doesn’t exactly share the A or C domain’s history. In this analysis, we see a sharp boundary between the two trees within the conserved FFxxGGxS motif in the T domain (Figs. 1c and Supplementary Fig. 4). Interestingly, the second history has a topology similar to the C domain tree (Figs. 1d and Supplementary Fig. 5). It also contains a clear split that separates T domains according the condensation reaction catalysed by the downstream C domains (Fig. 1d). The first tree, however, is not similar to either the C or A domain trees (Fig. 1c). Taken together, these observations suggest that the T domain may be a frequent recombination site, with a particularly important boundary in the conserved FFxxGGxS motif in the T domain.

To further confirm these *in silico* predictions and to avoid the result being a computational artefact, we have analysed in detail examples of homologous NRPS such as the PAX^38,39^, endopyrrole A^40^, rhizomide A^41^, and syringopeptin SP-25a^42^ producing synthetases to obtain evidence of recombination events within the T domains. In brief, this detailed analysis indeed supported the notion that recombination events within T domains frequently occur either to introduce a stereochemistry change (T-C vs T-C/E), and/or to exchange T-TE domains, and/or to increase/decrease the size of the BGC and the respective NP scaffold. A detailed description of this analysis can be found in the supporting information (Supplementary Figs. 6 – 9).

In summary, the results gained from the phylogenetic HMM (Fig. 1 and Supplementary Figs. 1 – 5) and the detailed analysis of various BGCs results point towards a yet undescribed recombination breakpoint.

### Fusion Point Screening

The conserved core motif (FFxxGGxS) of the ~100 amino acid T domains is located at the *N*-terminus (loop1) of the second helix (α2) holding the invariant serine residue that becomes post-translationally modified by a phosphopantetheinyl (Ppant) transferase^43–45^. Although the T domain is the only NRPS domain without an autonomous catalytic activity, the attachment of Ppant is a functional prerequisite, not only to covalently bind activated extender units and the growing peptide chain, but also to pervade the active sites of A and C domains. In addition, it is known from structural data that the first part of the T-domain (T_p1_), which is *N*-terminal to the core motif, mainly interacts with the A domain via α1 and loop1 and the second half (T_p2_), which is *C*-terminal to the core motif, interacts with the C domain via α4^46^. However, as computational recombination analysis (Fig. 1) naturally does not come up with one specific splicing position but with a sequence region that is likely to promote homologous recombination, initially a fusion point screening was performed (Fig. 2) to verify fusion sites resulting in the best peptide production.

**Fig. 2.**
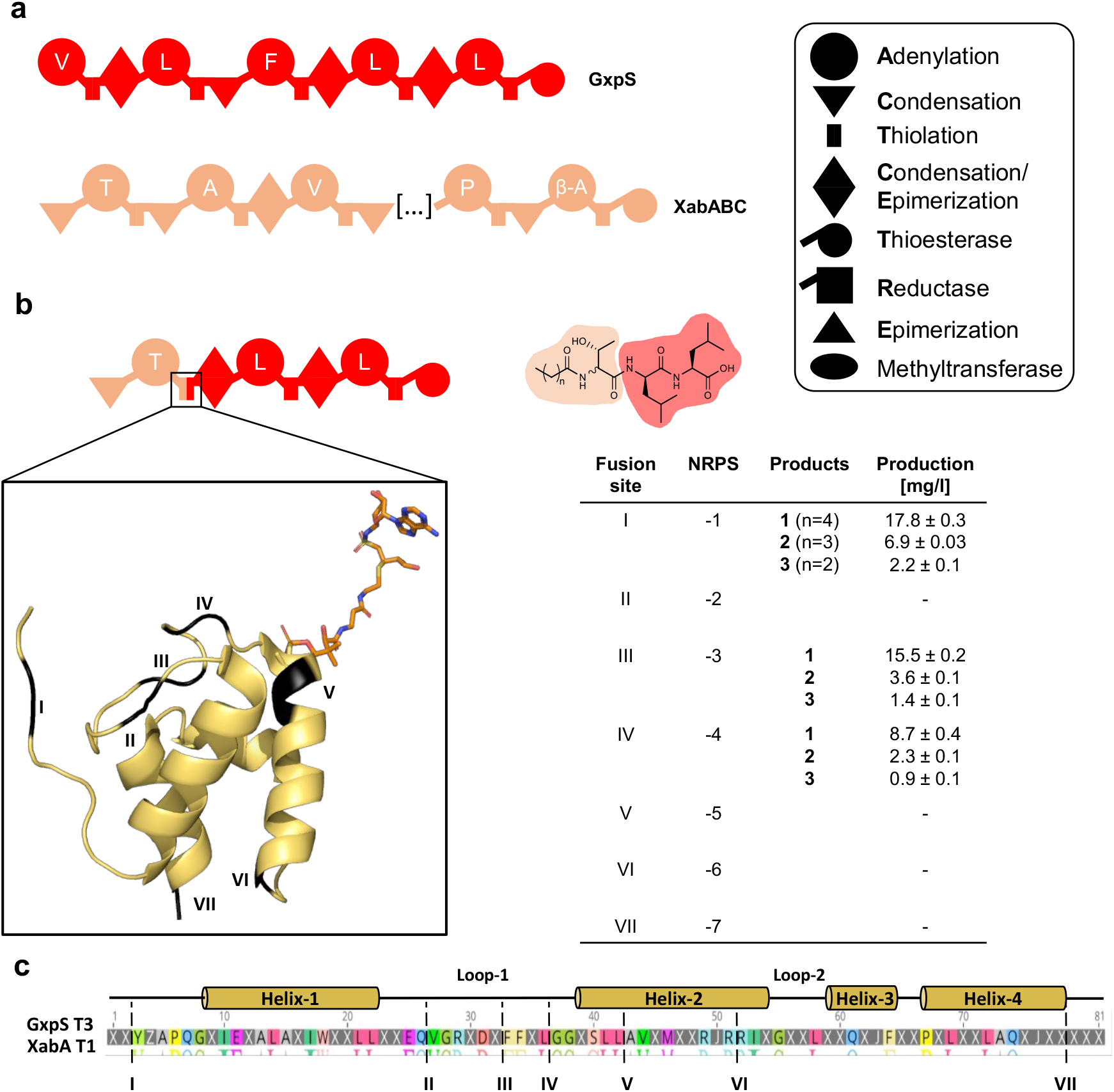
Fusion point screening of a NRPS hybrid assembled within the T domain. (**a**) Schematic representation of precursor NRPS GxpS and XabABC, producing GameXPeptides and xenoamicins, respectively. All domain functions are explained in the box. (**b**) Schematic representation of the XabA-GxpS hybrid NRPS, produced lipopeptide and compounds titres. The colour code of the peptide structure follows the NRPS colour code. The different fusion sites within the T domain are highlighted in black at the respective positions in the crystal structure of the T domain EntF (PDB 4ZXJ)^62^. (**c**) Sequence alignment of GxpS T3 and XabA T1 with secondary structures of the T domain and fusion sites indicated.

As a starting point, we chose the GameXPeptide^47^ (GxpS) and the xenoamicin^48^ producing synthetases (XabABC) from *P. luminescens* TT01 and *X. stockiae* (Fig. 2a), respectively, to produce seven recombinant NRPS (Fig. 2b, NRPS-1 to -7), each with a different fusion site (I to VII, Fig. 2c), with the *in silico* predicted breakpoint represented by fusion site IV. Briefly, this screening led to the identification of three functional fusion sites (I, III, and IV) in NRPS-1, -3 and -4, that all produce the expected lipopeptides **1**-**3**, differing only in the acyl starter originating from the fatty acid pool of *E. coli*, with titres between 12 and 27 mgL^-1^ (Fig. 2b, Supplementary Figs. 10 – 17, and Supplementary Table S6). Of note, throughout the present work, all NRPS were heterologously produced in *E. coli* DH10B∷mtaA^49^. The resulting peptides (Supplementary Table S5) and yields were confirmed by HPLC-MS/MS and comparison of retention times with synthetic standards (see Supplementary Information).

Taken together, the *in silico* observations (Fig. 1) along with the results from the *in vivo* conducted fusion site screening (Fig. 2) led us to the hypothesis that both, T-C-A units (fusion point I), Tp_1_-C-A-Tp_2_ units (fusion points III & IV), and combinations thereof may serve as ideal starting points to do rational evolution-inspired megasynthetase engineering. However, after reviewing crystal structure data of A-T and T-C didomains we decided to proceed with fusion site I and IV, because fusion sites III and IV are both located directly adjacent (III) and within (IV) the conserved T domain motif, respectively, and the two variable positions in between the conserved motif (FF**xx**GGxS) are potentially contributing to a functional A-T interface^46^.

### Evolution Inspired eXchange Units for NRPS Engineering

To further verify the *in silico* identified (IV) and *in vivo* verified (I & IV) fusion sites on a broad scale we targeted the NRPS FitAB (Fig. 3, NRPS-8) and FtrAB (Supplementary Fig. 18, NRPS-17; Supplementary Dataset 3) producing the NRPs fitayylide and faTTTVIR from *X. innexii and X. mauleonii*, respectively, as well as GxpS (Fig. 4). In sum we created 16 recombinant FitAB derivatives (NRPS-9 to -18, Fig. 3), one ftrAB derivative (NRPS-19 and -20, Fig. S18), and eight GxpS derivatives (Fig. 4) applying fusion site I, IV, or both. The building blocks to engineer NRPS-8, NRPS-17 and GxpS were selected to cover a broad range of bacterial genera (*Xenorhabdus, Photorhabdus, Serratia, Myxococcus, Pseudomonas* and *Bacillus*) with GC contents between 50 to 72 % GC to reveal if the identified fusion sites have the potential to mimic horizontal gene transfer along with recombination on a rational scale suitable to re-engineer NRPS.

**Figure 3.**
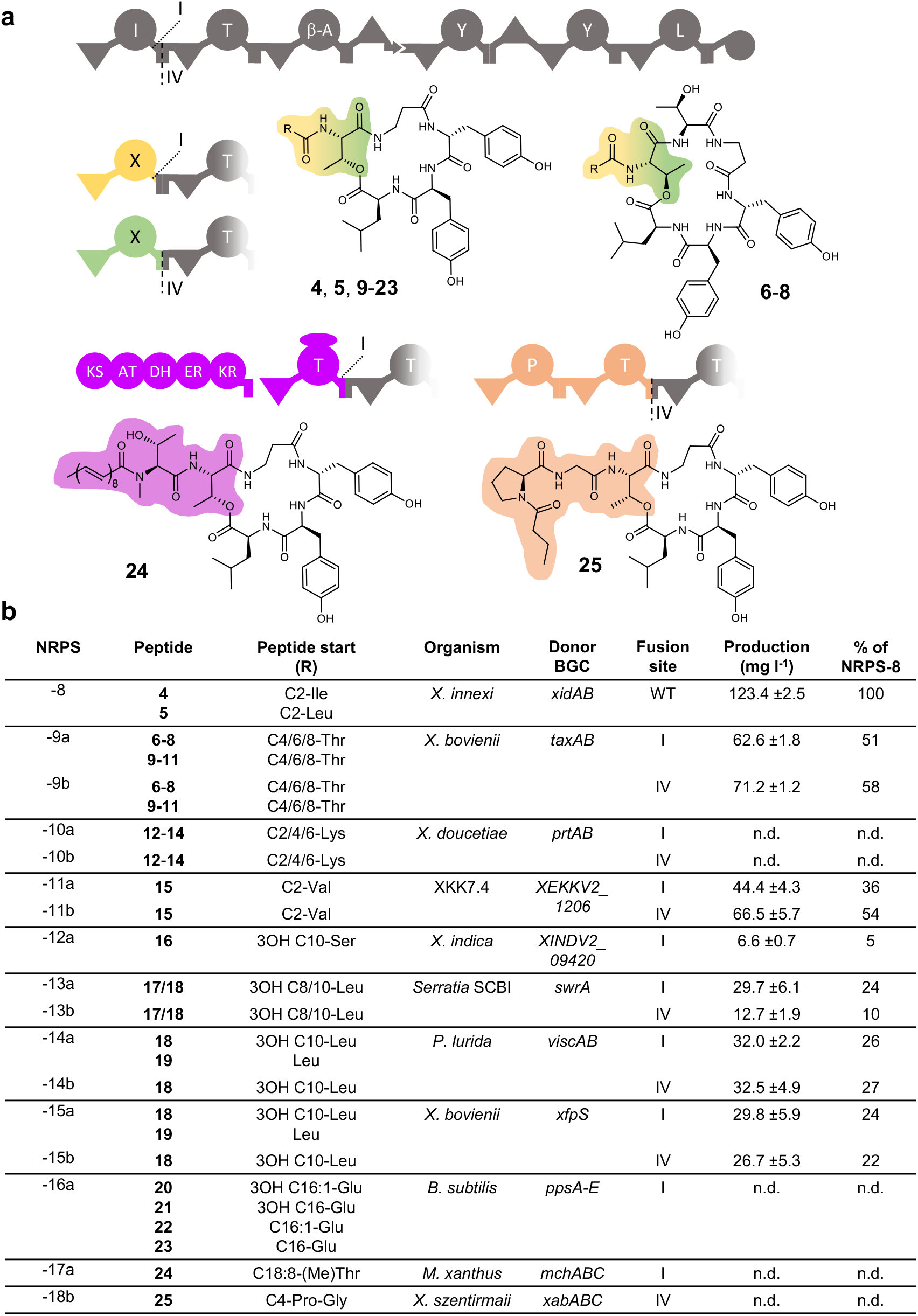
Evolution inspired exchange units replacing NRPS starting modules. (**a**) Schematic representation of the FitAB NRPS producing fitayylide A (**4**) and B (**5**) and selected alternative starting modules from other NRPS with indicated fusion sites I and IV. Amino acid specificities are assigned for all A domains. KS (ketosynthase), AT (acyltransferase), DH (dehydratase), ER (enoylreductase). Selected structures of the produced peptides are shown so that in conjunction with the table in (**b**) all peptide structures can be deduced. Production data relative to the WT NRPS-8 and the absolute peptide yields are based on triplicate production cultures. The origin of the alternative starting module, their cognate gene cluster and the fusion point for each starter module is shown. Production was observed for all NRPS derivatives, but production titres were not determined for all of them (n.d.).

**Fig. 4.**
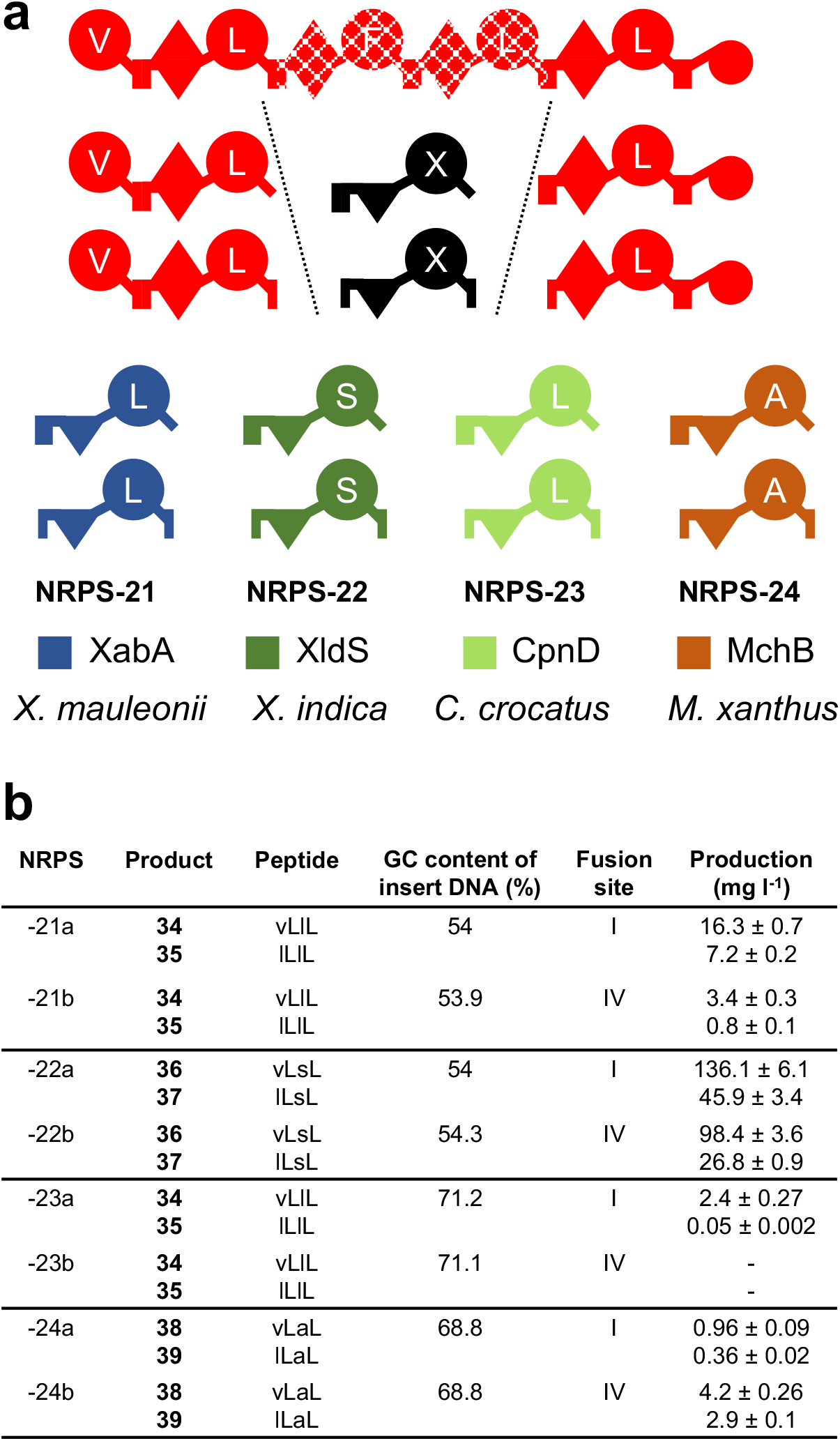
Evolution inspired exchange units replacing internal modules. (**a**) Schematic representation of the precursor NRPS GxpS producing GameXPeptides. A T_2_-T_4_ fragment was exchanged with different XUTs from the xenoamicine, xenolindicine, crocapeptin and myxochromide producing NRPS XabA, XldS, CpnD and MchC, respectively. (**b**) Peptides, GC content of the inserted XUTs, corresponding fusion sites and production titres of the respective peptides.

Interestingly all recombinant NRPS (including a NRPS-PKS assembly line where the PKS is responsible for the polyunsaturated starter acyl moiety [NRPS-19]), showed catalytic activity producing a broad range of cyclic and linear peptides (**4**-**39**) at titres ranging from 2.5 (NRPS-23a) to 136 mgL^−1^ (NRPS-22a) and from 2.5 (NRPS-18b) to 98 mgL^−1^ (NRPS-22b) for fusion site I and IV, respectively (Figs. 3, 4, Supplementary Figs. 18 – 55, and Supplementary Tables 7 – 9). In addition, and as already indicated from our initial fusion point screening (Fig. 2), no trend concerning a preferred fusion site could be observed.

Noteworthy, both fusion sites of the evolution-inspired exchange units between T domains (XUT) enabled us to create chimeric NRPS from completely unrelated BGCs for the first time with respect to taxonomy and GC content. Other methods, such as A subdomain swaps^15,31^ or the previously introduced eXchange unit concepts^13,14^, enabled efficient reprogramming of NRPS only within a narrow range of related BGCs.

Nevertheless, a correlation between GC content of the introduced NRPS building blocks and peptide production can be observed (Fig. 3 and 4). Whereas building blocks of genera with a similar or slightly higher (50 to 65 %) GC content (i.e., NRPS-13 and NRPS-14; Fig. 3) are generally well tolerated, building blocks originating from the high-GC branch (~70 %, i.e. NRPS-17, -23, and -24; Fig. 3 and 4) are resulting in impaired assembly lines when recombined with NRPS originating from *Xenorhabdus* and *Photorhabdus*. The initial reduction of catalytic activity when building blocks of different GC-content are recombined with each other might also occur naturally during homologous recombination after a horizontal gene transfer event.

### Evolution Inspired eXchange Units allow targeted peptide production

In order to validate the strength of these evolution-inspired exchange units (XUT), an artificial biosynthetic assembly line producing a novel pharmacological active peptide against a well characterised target was designed *de novo*. We chose the eukaryotic proteasome as target which plays pivotal roles in protein homeostasis affecting cell cycle, signal transduction and general cell physiology. Proteasomes are a family of *N*-terminal nucleophilic hydrolases consisting of two sets of seven copies of α and β subunits that assemble into a barrel-shaped complex (Fig. 5)^50^. Peptides inhibiting the proteasome, such as the clinically used bortezomib^51^, can lead to apoptosis, making the human proteasome a target for anti-cancer chemotherapy. Similar to well-known strategies from the pharmaceutical industry, we used the lipopeptide aldehyde fellutamide B^52^ as inspiration from nature that is not only active against the eukaryotic proteasome of humans and yeast, but is also the most potent inhibitor of the *Mycobacterium tuberculosis* proteasome tested to date. Fellutamide B consist of a C8-3OH acyl chain, L-Asn, L-Gln, and a L-Leu-aldehyde. The aldehyde moiety is responsible for the reversible binding to the active site threonine (Thr1) of the proteasome. From an NRPS engineering perspective, in particular the introduction of reactive groups, denoted as warheads, is a major challenge. As an alternative to TE domains, nature applies thioester reductase (R) domains^53–55^, not only to release the synthesised peptide, but also to introduce the aldehyde function by catalysing an NAD(P)H dependent two-electron reduction of the thioester.

**Fig. 5.**
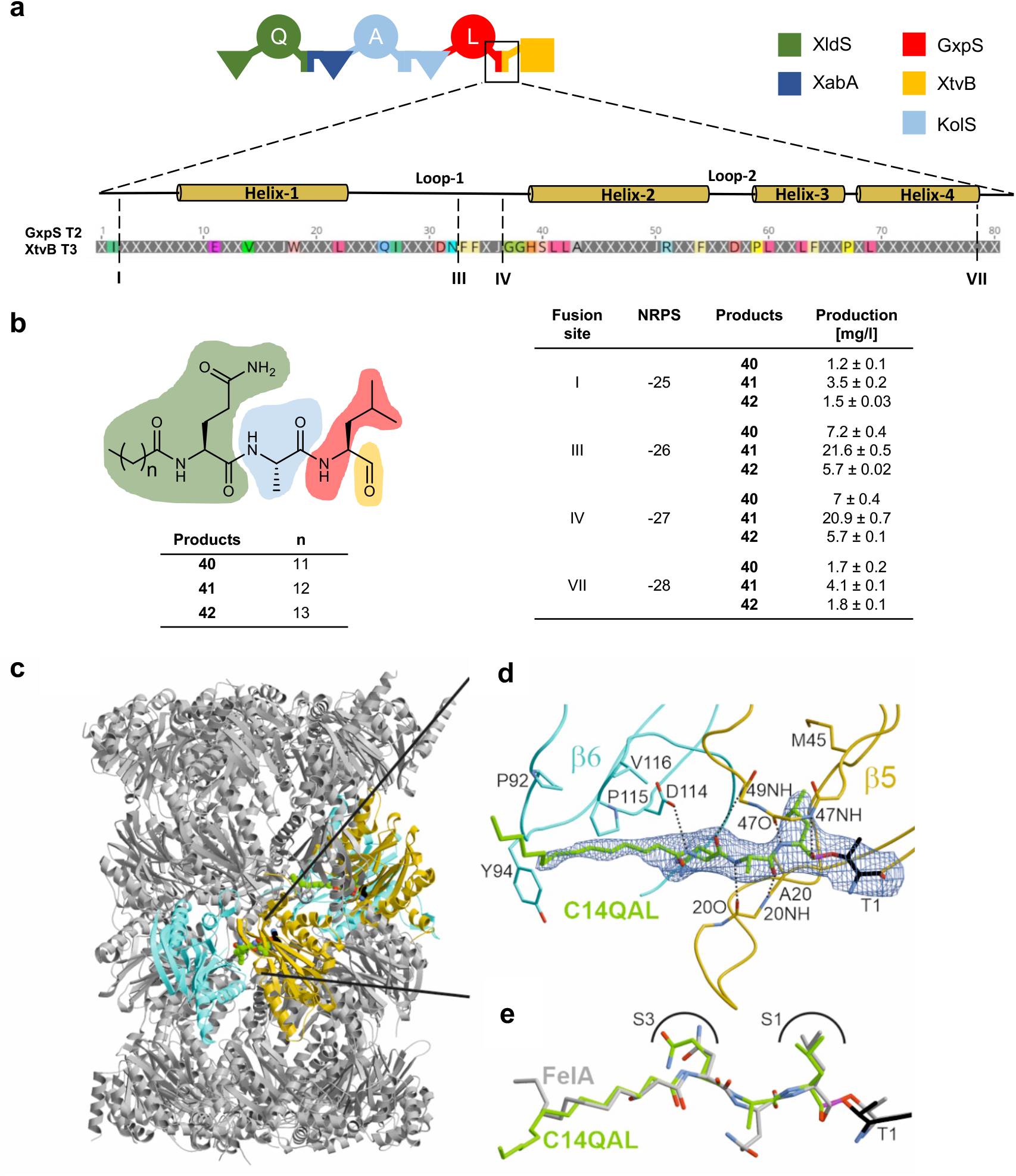
XUT approach for the design of a proteasome inhibitor. (**a**) Schematic representation of reassembled NRPS-25 to -28 composed of NRPS fragments from XldS, XabA, KolS (kolossin), GxpS and XtvB. The terminal T domain is shown as a sequence alignment of GxpS T2 and XtvB T3 indicating secondary structures and fusion sites. (**b**) Production titres corresponding to the fusion sites within the terminal T domain. The colour code in the peptides follows that of the NRPS fragments used. (**c**) Crystal structure of the yeast 20S proteasome in complex with **41** (spherical model, green carbon atoms) bound to the chymotrypsin-like active sites (β5 subunits, gold, PDB ID 8BW1). (**d**) Illustration of the 2*F*_O_−*F*_C_ electron density map (blue mesh, contoured to 1σ) of **41** (depicted as C14-QAL) covalently linked through a hemiacetal bond (magenta) to Thr1O^γ^. Protein residues interacting with **41** are highlighted in black. Dots illustrate hydrogen bonds between **41** and protein residues. (**e**) Superposition of **41** (depicted as C14-QAL) (green) and fellutamide A (grey, PDB ID s3D29)^63^ complex structures highlighting similar conformations at the chymotrypsin-like active site.

For the final XUT proof-of-concept experiment we *in silico* designed an artificial three-modular assembly line composed from NRPS building blocks derived from five different origins (Fig. 5a): a C_start_ domain to introduce the acyl chain and A domains with specificities (*N-* to *C*-terminus) for L-Gln (A1), L-Ala (A2), and L-Leu (A3). To achieve the reduction of leucine into an aldehyde, the R-domain of the tilivalline producing NRPS (XtvB)^56^ from *X. indica* was used as termination domain using the fusions sites I, III, IV, and VII. The resulting assembly lines NRPS-25 to -28 all showed catalytic activity producing the desired lipopeptide aldehydes **40**–**42**, differing only in the acyl group used as a starter, with titres between ~1 and ~22 mgL^−1^ (Fig. 5b, Supplementary Figs. 56 – 62, and Supplementary Table 10). Compared to the NRPS generated with fusion sites I (NRPS-25) and VII (NRPS-28), the NRPS generated with fusion sites III (NRPS-26) and IV (NRPS-27) produced about tenfold more peptides. Whereas low titres of NRPS-28 are in good agreement with our initial fusion point screening (Fig. 2), the peptide amount biosynthesised by NRPS-25 was unexpectedly low. The impaired formation of a functional A-T domain-domain interface^46^, in the case of NRPS-25, and a functional T-R^53^ domain-domain interface in the case of NRPS-28 could serve as an explanation for this result, as shown previously^56,57^. Furthermore, these results highlight the advantages of the evolution-inspired fusion sites III and IV compared to fusion sites I and VII, which are located within the A-T and T-C linker regions, respectively.

In order to test whether the new-to-nature lipopeptide aldehyde **41** is indeed able to inhibit the yeast 20S proteasome core particle (yCP) by binding the active site Thr1, the half maximum inhibitory concentration (IC_50_) and co-crystallization of yCP together with **41** (yCP:41 complex) was performed (Supplementary Fig. 63 and Supplementary Table 11). Both experiments confirmed the expected activity of **41** against the yCP β5 subunit at 3.6 ± 0.8 μM and a binding mode to Thr1 equivalent to that of fellutamide B (Fig. 5c-e). In summary, this proof-of-concept experiment not only revealed that reactive groups efficiently can be introduced by applying the novel XUT approach but also that tailormade bioactive peptides can be created *de novo* in a retro-biosynthetic manner.

## Conclusion

Despite all the technical advances and our knowledge of the fundamental biochemical and structural properties of assembly line enzymes^4^, their engineering has remained a major challenge^58^. Nature, however, appears to have been successful at engineering biosynthetic pathways through the process of BGC evolution using a broad range of mechanisms. Previous studies either comprehensively analysed a diverse range of NRPS families or focused on deciphering the evolution of one specific NRP family. Both approaches have dramatically improved our understanding of the underlying mechanisms of megasynthetase evolution. Pioneering studies for example proposed the *N*-terminal expansion of modules in BGC evolution^59^, highlighted the role of the A domains in NRPS diversification^24,30,32^, and introduced models explaining the mechanisms resulting in present day NRPS families^24,33,60^. However, most of these studies have not succeeded in developing these findings into an overall rational engineering approach. When the available datasets describing the evolution of NRPS synthesising syringopeptin^59^, jessipeptin, virginafactin, chicofactin, and syringafactin^60^ were reanalysed, we clearly could identify the T domain as an additional recombination hot spot.

Compared to these previous studies the major aim of this work was not the identification of the exact mechanisms that led to present day NRPS families, but to understand the major driving forces in NRPS evolution and how these insights can be leveraged to improve rational engineering of assembly line enzymes. Based on our findings we propose a yet undescribed recombination breakpoint within the conserved core motif of T domains (fusion sites III and IV), resulting in the XUT Tp_1_-C-A-Tp_2_. Interestingly, the XUT approach is completely in line with recent structural findings on the catalytic cycle of NRPS^46,61^ and, with exceptions, mostly consistent with the recently introduce unifying model for the evolution of the present-day variety of NRPs^24^. Although we are convinced that A subdomain exchanges are another important driver for NRPS evolution, our data does not suggest such a recombination, probably because the two data sets and the method of analysis are fundamentally different. From an applied engineering perspective, XUT appears to be much more versatile compared to A subdomain swaps^15,31,32^, allowing the rational recombination of completely unrelated NRPS building blocks over a broad range of GC contents (from 50% to 70 %), protein similarities (< 39 %), and extender unit specificities (Figs. 3 and 4).

To conclude, the XUT approach enables the mimicking of horizontal gene transfer followed by a recombination event, opening up avenues for the expansion of structural diversity that we can address through rational engineering – even beyond natural diversity. This is clearly illustrated by the example of the artificial proteasome inhibitor (Fig. 5) leading to the first time rational *de novo* design of a new-to-nature pharmacologically active peptide.

## Acknowledgements

This work was supported by an ERC Advanced Grant (835108), the LOEWE Research center TBG funded by the State of Hesse, the Max-Planck Society (all to H.B.B.), and the German Research Foundation (SFB1035, project number: 201302640, A02 to W.K. and M.G.). The authors thank the staff of beamline X06SA at the Paul Scherrer Institute, SLS, Villigen (Switzerland), for their assistance during data collection and are grateful to all Bode lab members for continuous discussions about further developments of NRPS engineering methods.

## Author contributions

K.A.J.B., L.P., C.K., L.S. and T.M.M. planned and performed all NRPS engineering experiments and peptide productions. Y.-N.S. isolated all peptides and elucidated their structure. S.K., C.K. and G.K.A.H. performed all phylogenetic analyses. W.K. and M.G. performed proteasome assays and crystallization. K.A.J.B. and H.B.B. conceived all experiments and wrote the paper with input from all authors.

## References

1 Newman, D. J. & Cragg, G. M. Natural Products as Sources of New Drugs over the Nearly Four Decades from 01/1981 to 09/2019. J Nat Prod 83, 770–803, doi:10.1021/acs.jnatprod.9b01285 (2020).

2 Huang, M., Lu, J. J. & Ding, J. Natural Products in Cancer Therapy: Past, Present and Future. Nat Prod Bioprospect 11, 5–13, doi:10.1007/s13659-020-00293-7 (2021).

3 David, F. et al. A Perspective on Synthetic Biology in Drug Discovery and Development-Current Impact and Future Opportunities. SLAS Discov 26, 581–603, doi:10.1177/24725552211000669 (2021).

4 Sussmuth, R. D. & Mainz, A. Nonribosomal Peptide Synthesis-Principles and Prospects. Angew Chem Int Ed Engl 56, 3770–3821, doi:10.1002/anie.201609079 (2017).

5 Tahlan, K., Moore, M. A. & Jensen, S. E. delta-(L-alpha-aminoadipyl)-L-cysteinyl-D-valine synthetase (ACVS): discovery and perspectives. J Ind Microbiol Biotechnol 44, 517–524, doi:10.1007/s10295-016-1850-7 (2017).

6 Byford, M. F., Baldwin, J. E., Shiau, C. Y. & Schofield, C. J. The Mechanism of ACV Synthetase. Chem Rev 97, 2631–2650, doi:10.1021/cr960018l (1997).

7 Zhang, J. & Demain, A. L. ACV synthetase. Crit Rev Biotechnol 12, 245–260, doi:10.3109/07388559209069194 (1992).

8 Umezawa, H. Structure and action of bleomycin. Prog Biochem Pharmacol 11, 18–27 (1976).

9 Fahr, A. Cyclosporin clinical pharmacokinetics. Clin Pharmacokinet 24, 472–495, doi:10.2165/00003088-199324060-00004 (1993).

10 Caboche, S., Leclere, V., Pupin, M., Kucherov, G. & Jacques, P. Diversity of monomers in nonribosomal peptides: towards the prediction of origin and biological activity. J Bacteriol 192, 5143–5150, doi:10.1128/JB.00315-10 (2010).

11 Walsh, C. T., O’Brien, R. V. & Khosla, C. Nonproteinogenic amino acid building blocks for nonribosomal peptide and hybrid polyketide scaffolds. Angew Chem Int Ed Engl 52, 7098–7124, doi:10.1002/anie.201208344 (2013).

12 Bozhuyuk, K. A., Micklefield, J. & Wilkinson, B. Engineering enzymatic assembly lines to produce new antibiotics. Curr Opin Microbiol 51, 88–96, doi:10.1016/j.mib.2019.10.007 (2019).

13 Bozhuyuk, K. A. J. et al. Modification and de novo design of non-ribosomal peptide synthetases using specific assembly points within condensation domains. Nat Chem 11, 653–661, doi:10.1038/s41557-019-0276-z (2019).

14 Bozhuyuk, K. A. J. et al. De novo design and engineering of non-ribosomal peptide synthetases. Nat Chem 10, 275–281, doi:10.1038/nchem.2890 (2018).

15 Thong, W. L. et al. Gene editing enables rapid engineering of complex antibiotic assembly lines. Nat Commun 12, 6872, doi:10.1038/s41467-021-27139-1 (2021).

16 Niquille, D. L. et al. Nonribosomal biosynthesis of backbone-modified peptides. Nat Chem 10, 282–287, doi:10.1038/nchem.2891 (2018).

17 Huang, H. M., Stephan, P. & Kries, H. Engineering DNA-Templated Nonribosomal Peptide Synthesis. Cell Chem Biol 28, 221–227 e227, doi:10.1016/j.chembiol.2020.11.004 (2021).

18 Abbood, N., Duy Vo, T., Watzel, J., Bozhueyuek, K. A. J. & Bode, H. B. Type S Non-Ribosomal Peptide Synthetases for the Rapid Generation of Tailormade Peptide Libraries. Chemistry, e202103963, doi:10.1002/chem.202103963 (2022).

19 Bozhueyuek, K. A. J., Watzel, J., Abbood, N. & Bode, H. B. Synthetic Zippers as an Enabling Tool for Engineering of Non-Ribosomal Peptide Synthetases*. Angew Chem Int Ed Engl 60, 17531–17538, doi:10.1002/anie.202102859 (2021).

20 Klaus, M., D’Souza, A. D., Nivina, A., Khosla, C. & Grininger, M. Engineering of Chimeric Polyketide Synthases Using SYNZIP Docking Domains. ACS Chem Biol 14, 426–433, doi:10.1021/acschembio.8b01060 (2019).

21 Kautsar, S. A., Blin, K., Shaw, S., Weber, T. & Medema, M. H. BiG-FAM: the biosynthetic gene cluster families database. Nucleic Acids Res 49, D490–D497, doi:10.1093/nar/gkaa812 (2021).

22 Blin, K. et al. antiSMASH 6.0: improving cluster detection and comparison capabilities. Nucleic Acids Res 49, W29–W35, doi:10.1093/nar/gkab335 (2021).

23 Blin, K., Shaw, S., Kautsar, S. A., Medema, M. H. & Weber, T. The antiSMASH database version 3: increased taxonomic coverage and new query features for modular enzymes. Nucleic Acids Res 49, D639–D643, doi:10.1093/nar/gkaa978 (2021).

24 Baunach, M., Chowdhury, S., Stallforth, P. & Dittmann, E. The Landscape of Recombination Events That Create Nonribosomal Peptide Diversity. Mol Biol Evol 38, 2116–2130, doi:10.1093/molbev/msab015 (2021).

25 Booth, T. J., Bozhüyük, K. A. J., Liston, J. D., Lacey, E. & Wilkinson, B. Bifurcation drives the evolution of assembly-line biosynthesis. bioRxiv, 2021.2006.2023.449585, doi:10.1101/2021.06.23.449585 (2021).

26 Nivina, A., Yuet, K. P., Hsu, J. & Khosla, C. Evolution and Diversity of Assembly-Line Polyketide Synthases. Chem Rev 119, 12524–12547, doi:10.1021/acs.chemrev.9b00525 (2019).

27 Vander Wood, D. A. & Keatinge-Clay, A. T. The modules of trans-acyltransferase assembly lines redefined with a central acyl carrier protein. Proteins 86, 664–675, doi:10.1002/prot.25493 (2018).

28 Keatinge-Clay, A. T. Polyketide Synthase Modules Redefined. Angew Chem Int Ed Engl 56, 4658–4660, doi:10.1002/anie.201701281 (2017).

29 Nivina, A., Herrera Paredes, S., Fraser, H. B. & Khosla, C. GRINS: Genetic elements that recode assembly-line polyketide synthases and accelerate their diversification. Proc Natl Acad Sci U S A 118, doi:10.1073/pnas.2100751118 (2021).

30 Calcott, M. J., Owen, J. G. & Ackerley, D. F. Efficient rational modification of non-ribosomal peptides by adenylation domain substitution. Nat Commun 11, 4554, doi:10.1038/s41467-020-18365-0 (2020).

31 Kries, H., Niquille, D. L. & Hilvert, D. A subdomain swap strategy for reengineering nonribosomal peptides. Chem Biol 22, 640–648, doi:10.1016/j.chembiol.2015.04.015 (2015).

32 Crüsemann, M., Kohlhaas, C. & Piel, J. Evolution-guided engineering of nonribosomal peptide synthetase adenylation domains. Chem. Sci., 1041–1045, doi:10.1039/C2SC21722H (2013).

33 Meyer, S. et al. Biochemical Dissection of the Natural Diversification of Microcystin Provides Lessons for Synthetic Biology of NRPS. Cell Chem Biol 23, 462–471, doi:10.1016/j.chembiol.2016.03.011 (2016).

34 Boussau, B., Gueguen, L. & Gouy, M. A mixture model and a hidden markov model to simultaneously detect recombination breakpoints and reconstruct phylogenies. Evol Bioinform Online 5, 67–79, doi:10.4137/ebo.s2242 (2009).

35 Stachelhaus, T., Mootz, H. D. & Marahiel, M. A. The specificity-conferring code of adenylation domains in nonribosomal peptide synthetases. Chem Biol 6, 493–505, doi:10.1016/S1074-5521(99)80082-9 (1999).

36 Rausch, C., Hoof, I., Weber, T., Wohlleben, W. & Huson, D. H. Phylogenetic analysis of condensation domains in NRPS sheds light on their functional evolution. BMC Evol Biol 7, 78, doi:10.1186/1471-2148-7-78 (2007).

37 Wheadon, M. J. & Townsend, C. A. Evolutionary and functional analysis of an NRPS condensation domain integrates beta-lactam, ᴅ-amino acid, and dehydroamino acid synthesis. Proc Natl Acad Sci U S A 118, doi:10.1073/pnas.2026017118 (2021).

38 Fuchs, S. W., Proschak, A., Jaskolla, T. W., Karas, M. & Bode, H. B. Structure elucidation and biosynthesis of lysine-rich cyclic peptides in Xenorhabdus nematophila. Org Biomol Chem 9, 3130–3132, doi:10.1039/c1ob05097d (2011).

39 Vo, T. D., Spahn, C., Heilemann, M. & Bode, H. B. Microbial Cationic Peptides as a Natural Defense Mechanism against Insect Antimicrobial Peptides. ACS Chem Biol 16, 447–451, doi:10.1021/acschembio.0c00794 (2021).

40 Niehs, S. P. et al. Genome Mining Reveals Endopyrroles from a Nonribosomal Peptide Assembly Line Triggered in Fungal-Bacterial Symbiosis. ACS Chem Biol 14, 1811–1818, doi:10.1021/acschembio.9b00406 (2019).

41 Wang, X. et al. Discovery of recombinases enables genome mining of cryptic biosynthetic gene clusters in Burkholderiales species. Proc Natl Acad Sci U S A 115, E4255–E4263, doi:10.1073/pnas.1720941115 (2018).

42 Isogai, A. et al. Structural analysis of new syringopeptins by tandem mass spectrometry. Biosci Biotechnol Biochem 59, 1374–1376, doi:10.1271/bbb.59.1374 (1995).

43 Sieber, S. A. & Marahiel, M. A. Molecular mechanisms underlying nonribosomal peptide synthesis: approaches to new antibiotics. Chem Rev 105, 715–738, doi:10.1021/cr0301191 (2005).

44 Zhou, Z., Lai, J. R. & Walsh, C. T. Directed evolution of aryl carrier proteins in the enterobactin synthetase. Proc Natl Acad Sci U S A 104, 11621–11626, doi:10.1073/pnas.0705122104 (2007).

45 Lai, J. R., Koglin, A. & Walsh, C. T. Carrier protein structure and recognition in polyketide and nonribosomal peptide biosynthesis. Biochemistry 45, 14869–14879, doi:10.1021/bi061979p (2006).

46 Reimer, J. M. et al. Structures of a dimodular nonribosomal peptide synthetase reveal conformational flexibility. Science 366, doi:10.1126/science.aaw4388 (2019).

47 Nollmann, F. I. et al. Insect-specific production of new GameXPeptides in photorhabdus luminescens TTO1, widespread natural products in entomopathogenic bacteria. Chembiochem 16, 205–208, doi:10.1002/cbic.201402603 (2015).

48 Zhou, Q. et al. Structure and biosynthesis of xenoamicins from entomopathogenic Xenorhabdus. Chemistry 19, 16772–16779, doi:10.1002/chem.201302481 (2013).

49 Schimming, O., Fleischhacker, F., Nollmann, F. I. & Bode, H. B. Yeast homologous recombination cloning leading to the novel peptides ambactin and xenolindicin. Chembiochem 15, 1290–1294, doi:10.1002/cbic.201402065 (2014).

50 Baumeister, W., Walz, J., Zuhl, F. & Seemuller, E. The proteasome: paradigm of a self-compartmentalizing protease. Cell 92, 367–380, doi:10.1016/s0092-8674(00)80929-0 (1998).

51 Scott, K., Hayden, P. J., Will, A., Wheatley, K. & Coyne, I. Bortezomib for the treatment of multiple myeloma. Cochrane Database Syst Rev 4, CD010816, doi:10.1002/14651858.CD010816.pub2 (2016).

52 Lin, G., Li, D., Chidawanyika, T., Nathan, C. & Li, H. Fellutamide B is a potent inhibitor of the Mycobacterium tuberculosis proteasome. Arch Biochem Biophys 501, 214–220, doi:10.1016/j.abb.2010.06.009 (2010).

53 Deshpande, S., Altermann, E., Sarojini, V., Lott, J. S. & Lee, T. V. Structural characterization of a PCP-R didomain from an archaeal nonribosomal peptide synthetase reveals novel interdomain interactions. J Biol Chem 296, 100432, doi:10.1016/j.jbc.2021.100432 (2021).

54 Kavanagh, K. L., Jornvall, H., Persson, B. & Oppermann, U. Medium- and short-chain dehydrogenase/reductase gene and protein families : the SDR superfamily: functional and structural diversity within a family of metabolic and regulatory enzymes. Cell Mol Life Sci 65, 3895–3906, doi:10.1007/s00018-008-8588-y (2008).

55 Persson, B. & Kallberg, Y. Classification and nomenclature of the superfamily of short-chain dehydrogenases/reductases (SDRs). Chem Biol Interact 202, 111–115, doi:10.1016/j.cbi.2012.11.009 (2013).

56 Tietze, A., Shi, Y. N., Kronenwerth, M. & Bode, H. B. Nonribosomal Peptides Produced by Minimal and Engineered Synthetases with Terminal Reductase Domains. Chembiochem 21, 2750–2754, doi:10.1002/cbic.202000176 (2020).

57 Calcott, M. J. & Ackerley, D. F. Portability of the thiolation domain in recombinant pyoverdine non-ribosomal peptide synthetases. BMC Microbiol 15, 162, doi:10.1186/s12866-015-0496-3 (2015).

58 Brown, A. S., Calcott, M. J., Owen, J. G. & Ackerley, D. F. Structural, functional and evolutionary perspectives on effective re-engineering of non-ribosomal peptide synthetase assembly lines. Nat Prod Rep 35, 1210–1228, doi:10.1039/c8np00036k (2018).

59 Medema, M. H., Cimermancic, P., Sali, A., Takano, E. & Fischbach, M. A. A systematic computational analysis of biosynthetic gene cluster evolution: lessons for engineering biosynthesis. PLoS Comput Biol 10, e1004016, doi:10.1371/journal.pcbi.1004016 (2014).

60 Gotze, S. et al. Structure elucidation of the syringafactin lipopeptides provides insight in the evolution of nonribosomal peptide synthetases. Chem Sci 10, 10979–10990, doi:10.1039/c9sc03633d (2019).

61 Reimer, J. M., Haque, A. S., Tarry, M. J. & Schmeing, T. M. Piecing together nonribosomal peptide synthesis. Curr Opin Struct Biol 49, 104–113, doi:10.1016/j.sbi.2018.01.011 (2018).

62 Drake, E. J. et al. Structures of two distinct conformations of holo-non-ribosomal peptide synthetases. Nature 529, 235–238, doi:10.1038/nature16163 (2016).

63 Hines, J., Groll, M., Fahnestock, M. & Crews, C. M. Proteasome inhibition by fellutamide B induces nerve growth factor synthesis. Chem Biol 15, 501–512, doi:10.1016/j.chembiol.2008.03.020 (2008).

