## Supplementary material for "Evolution Inspired Engineering of Megasynthetases": Material & Methods, supplementary Tables and Figures.

### 1. Material and Methods

#### 1.1. Cultivation of strains

All *E. coli* DH10B::*mtaA* cells were cultured either on liquid or solid low salt LB medium ((pH 7.5, 10 g/L tryptone, 5 g/L yeast extract and 5 g/L NaCl). Either kanamycin (50 µg/ml), chloramphenicol (34 µg/ml) or spectinomycin (50 µg/ml) were added as selection markers. Solid media contained 1% (w/v) agar. Cells were cultivated at 37 °C and at 22 °C for peptide production cultures.

#### 1.2. Cloning of biosynthetic gene clusters and NRPS modules

For use as template, genomic DNA (gDNA) was extracted from bacteria indicated in Table S1 by use of the Gentra Puregene Yeast/Bact. Kit (Qiagen) and the Monarch® Genomic DNA Purification Kit (NEB) which in turn was taken as template for the PCR amplification. The proof-reading PCR polymerase Q5® High-Fidelity DNA Polymerase (NEB) and *Phusion* DNA Polymerase (NEB/Thermo Fisher Scientific) in their standard and hot start variations were employed. Oligonucleotides for the PCR and the correct product size are documented in Table S4. In specified cases (Table S4) already cloned NRPS parts were used as template for the PCR. PCR products were agarose gel purified taking the Monarch® DNA Gel Extraction Kit (NEB) to be used as substrate for the Gibson cloning procedure using the Gibson Assembly® Master Mix or the NEBuilder® HiFi DNA Assembly Cloning Kit (NEB). In cases indicated in Table S4 restriction enzyme digests with enzymes indicated were used as one part of the substrate for the Gibson cloning step.

The vector pCK_0407 was cloned in a classic fashion. To this end the plasmid pCK_0407 was linearised using the restriction enzymes AvrII/XbaI and the 1.750 bp fragment ligated to the 1.933 bp fragment of the AvrII/XbaI digest of pCDFDuet (Merck-Novagen).

#### 1.3. Heterologous expression of NRPS constructs and HPLC-MS analysis

After plasmid transformation into *E. coli* DH10B::*mtaA*, cells were grown overnight in LB medium containing all necessary antibiotics (50 µg/ml kanamycin, 34 µg/ml chloramphenicol, 100 µg/ml spectinomycin). 10 ml LB medium containing antibiotics, 0.002 mg/ml L-arabinose and 2 % (v/v) XAD-16 were inoculated with 1 % overnight grown culture. After incubation for 72 h at 22 °C, XAD-16 beads were harvested and one culture volume methanol was added. Methanol extraction was conducted for 60 min at 22 °C. The organic phase was filtrated and diluted 1:10 in methanol. Cleared HPLC-UV-MS analysis was conducted on an UltiMate 3000 system (Thermo Fisher) coupled to an AmaZonX mass spectrometer (Bruker) with an ACQUITY UPLC BEH C18 column

(130 Å, 2.1 mm × 100 mm, 1.7-µm particle size, Waters) at a flow rate of 0.4 ml min−1 (5–95% acetonitrile/water with 0.1% formic acid, vol/vol, 16 min, UV detection wavelength 190–800 nm). HPLC-UV-HRMS analysis was conducted on an UltiMate 3000 system (Thermo Fisher) coupled to an Impact II qTof mass spectrometer (Bruker) with an ACQUITY UPLC BEH C18 column (130 Å, 2.1 mm × 100 mm, 1.7-µm particle size, Waters) at a flow rate of 0.4 ml min−1 16 min, UV detection wavelength 190–800 nm). Evaluation was performed using DataAnalysis 5.3 software (Bruker).

For peptide quantification of NRPS-8- to -20 the production medium was, deviating from above, XPP medium[1] without phenylalanine with 1 mM ß-alanine added.

##### 1.4. Peptide Purification

Compounds **4**, **5**, **7**, **10**, **26**, **41** and **61** were produced in *E. coli* DH10B::*mtaA* expressing the respective NRPS variants. 4L XPP medium containing 34 µg/ml chloramphenicol, 0.002 % L-arabinose and 2 % XAD 16N beads was inoculated with 1 % overnight grown culture as described in section S1.3. The culture was incubated at 180 rpm for 72 h at 22 °C. Subsequently, the XAD 16N beads were extracted 3 times with 500 ml methanol for 30 minutes, stirring. Solvent was fully removed at reduced pressure and the crude extract was completely solved in DMSO in order to purify it by preparative HPLC–MS (LC-MS-System 1260 Infinity II Preparative LC/MSD from Agilent). A C3 column (Agilent ZORBAX 300XB-C3) utilizing a gradient of 40-55 % ACN/H2O (+0.1 % formic acid) was used. The compound was freeze-dried and the purity of the compound was determined by NMR and HPLC-HR-MS.

##### 1.5. Peptide quantification

The absolute production titres were calculated as previously described[2]. Therefore, calibration curves based on pure **1** (for quantification of **1**, **2** and **3**), **4** (**4**, **5**, **15**, **17**, **18**, **32** and **33**), **10** (**6**, **7**, **8**, **9**, **10**, **11** and **16**), **26** (**26**, **27**, **28**, **29**), **34** (**34** and **35**), **36** (**36** and **37**), **38** (**38** and **39**), and **41** (**40**, **41** and **42**), were prepared. The pure compounds were prepared at different concentrations: **1** utilizing a standard curve with concentrations of 5000, 500, 50, 5 and 0.5 µg $L^{-1}$; **4** utilizing a standard curve with concentrations of 10, 4, 1, 0.4, 0.1, 0.04 and 0.01 mg $l^{-1}$, **10** utilizing a standard curve with concentrations of 10, 4, 1, 0.4, 0.1 and 0.04 mg $l^{-1}$, **26** utilizing a standard curve with concentrations of 40, 4, 0.4, 0.04 and 0.004 mg $l^{-1}$, **34**, **36** and **38** utilizing a standard curve with concentrations of 100, 20, 4, 0.8 and 0.16 mg $l^{-1}$, **41** utilizing a standard curve with concentrations 10, 5, 2.5, 1.25, 0.625, 0.3125 and 0.1562 mg $l^{-1}$ and measured by LC-MS using HPLC/MS measurements as described above. To ensure sample signals being within the range of the standard curve they were diluted when necessary. The peak area for each compound at different concentrations was calculated using Compass Data Analysis and used for the calculation of a standard curve passing through the zero point. Triplicates of all *in vivo* experiments were measured. The pure peptide standards **1**, **34**, **36**, **38** were synthesized in-house, **4, 10**, **26** and **41** were purified from production cultures.

##### 1.6. Chemical Synthesis

The linear peptide **1** was synthesized on preloaded resin (0.25 mmol H-Leu-2ClTrt PS resin, Sigma Aldrich, Germany) by solid phase peptide synthesis using standard Fmoc/*t*-Bu chemistry. Fmoc protected amino acids or fatty acids were activated by mixture of 5 eq. Fmoc-AA-OH (or fatty acid), 12 eq. N,N-diisopropylethylamine (DIPEA, Iris Biotech, c = 2.4 M), 5 eq. O-(7-azabenzotriazol-1-yl)-N,N,N′,N′-tetramethyluronium hexafluorophosphate (HATU, Carbolution Chemicals) in 15 ml dimethylformamide (DMF, Carl Roth, Germany). The resin was incubated with the activated amino acid/fatty acid mixture for 2 h at room temperature. After each coupling, the resin was washed with NMP (5 ×), DMF (5 ×) and DCM (5 ×). Finally, the peptide was cleaved by addition of 20 ml of a mixture of Hexafluoroisopropanol (HFIP) and DCM (1:4 v/v). Subsequently, the peptide was deprotected upon addition of 2 ml Trifluoroacetic acid (TFA) incubating for 2 h at room temperature. The linear peptide was dissolved in MeOH in order to purify it by semi-preparative HPLC–MS (Agilent LC-MS-System 1260 Infinity II Analytical-Scale LC/MSD) utilizing a C18 column (Eclipse XDB-C18 (9.4 x 250 mm, 5 µm). The purity was determined by NMR and HPLC-HR-MS analysis.

Chemical synthesis of peptides **34**, **36**, **38** was performed as described previously[2]. The linear sequences were synthesized on preloaded resins (H-AA-2ClTrt PS resin, Sigma Aldrich, Germany) on a 25 µM scale with a Syro Wave peptide synthesizer (Biotage, Sweden) by using standard Fmoc/*t*-Bu chemistry. Fmoc-amino acids were purchased from Carbolution Chemicals (Germany), Iris Biotech (Germany) or Bachem (Switzerland). Therefore, the resin was placed in a plastic reactor vessel with a Teflon frit and an amount of 6 eq. of amino acid derivative (c = 0.2 M) was activated *in situ* at room temperature with 6 eq. of *O*-(6-chlorobenzotriazol-1-yl)-*N*,*N*,*N′*,*N′*-tetramethyluronium hexafluorophosphate (HCTU, Carl Roth, Germany, c = 0.6 M) in dimethylformamide (DMF, Carl Roth, Germany) in the presence of 12 eq. *N,N*-diisopropylethylamine (DIPEA, Iris Biotech, c = 2.4 M) in *N*-methylpyrrolidone (NMP, Iris Biotech) for 50 min. Fmoc-protecting groups were removed with a solution of 40 % piperidine (Iris Biotech) in NMP (v/v %) for 5 min and followed by a second deprotection step with 20 % piperidine in NMP (v/v %) for 10 min. After each coupling and deprotection step, the resin was washed with NMP (4

×). After addition of the final amino acid and deprotection step, the resin was washed with NMP (5 ×), DMF (5 ×) and DCM (5 ×).

For total deprotection or cleavage 0.5 mL 95 % trifluoroacetic acid (TFA, Iris Biotech) and 2.5 % triisopropylsilane (TIS, Sigma Aldrich) in water were added to peptidyl resin and the mixture was agitated for at least 1 h at room temperature. The resin was removed by filtration and washed twice with TFA. Then the cleavage cocktail was evaporated. Linear peptide was dissolved in MeOH in order to purify it by semi-preparative HPLC–MS (Agilent LC-MS-System 1260 Infinity II Analytical-Scale LC/MSD) utilizing a C18 column (Eclipse XDB-C18 (9.4 x 250 mm, 5 μm). The purity was determined by HPLC-HR-MS and NMR.

#### 1.7. Expression and purification of yeast 20 S proteasome

The yeast 20S proteasome was prepared as previously described[3,4].

#### 1.8. $IC_{50}$ value determination with purified yCP

Concentration of purified yeast 20 S proteasome (yCP) was determined spectrophotometrically at 280 nm. yCP (final concentration: 0.05 mg/mL in 100 mM Tris-HCl, pH 7.5) was mixed with DMSO as a control or serial dilutions of fellutamide derivatives in DMSO, thereby not surpassing a final concentration of 10% (v/v) DMSO. After an incubation time of 45 min at RT, fluorogenic substrates Boc-Leu-Arg-Arg-AMC, Z-Leu-Leu-Glu-AMC and Suc-Leu-Leu-Val-Tyr-AMC (final concentration of 200 μM) were added to measure the residual activity of caspase-like (C-L, β1 subunit), trypsin-like (T-L, β2 subunit) and chymotrypsin-like (ChT-L, β5 subunit), respectively. The assay mixture was incubated for another 60 min at RT and afterwards diluted 1:10 in 20 mM Tris-HCl, pH 7.5. The AMC-molecules released by hydrolysis were measured in triplicate with a Varian Cary Eclipse Fluorescence Spectrophotometer (Agilent Technologies) at $\lambda_{exc}$=360 nm and $\lambda_{em}$=460 nm. Relative fluorescence units were normalized to the DMSO treated control. The calculated residual activities were plotted against the logarithm of the applied inhibitor concentration and fitted with GraphPad Prism 5. Half maximum inhibitory concentration ($IC_{50}$) values were deduced from the fitted data. They depend on enzyme concentration and are comparable within the same experimental settings.

#### 1.9. Crystallisation and structure determination of the yeast 20S proteasome core particle (yCP) in complex with 41.

Crystals of yCP were grown in hanging drops at 20°C as previously described[3,4]. The protein concentration used for crystallization was 40 mg/mL in Tris / HCl (20 mM, pH 7.5) and EDTA (1 mM). The drops contained 1 µL of protein and 1 µL of the reservoir solution [30 mM magnesium acetate, 100 mM 2-(N-morpholino)ethanesulfonic acid (pH 6.8) and 10% (wt/vol) 2-methyl-2,4-pentanediol]. Crystals appeared after two days and were incubated with a fellutamide derivative at final concentrations of 10 mM for at least 24 h. Droplets were then complemented with a cryoprotecting buffer [30% (wt/vol) 2-methyl-2,4-pentanediol, 15 mM magnesium acetate, 100 mM 2-(N-morpholino)ethanesulfonic acid, pH 6.9] and vitrified in liquid nitrogen. The dataset from the yCP: **41** complex was collected using synchrotron radiation (λ =1.0 Å) at the X06SA-beamline (Swiss Light Source, Villingen, Switzerland). X-ray intensities and data reduction were evaluated using the XDS program package (Table Sx)[5]. Conventional crystallographic rigid body, positional, and temperature factor refinements were carried out with REFMAC5[6] using coordinates of the yCP structure as starting model (PDB ID 5CZ4)[7]. For model building, the programs SYBYL and COOT[8] were used. The final coordinates yielded excellent R factors, as well as geometric bond and angle values. Coordinates were confirmed to fulfill the Ramachandran plot and have been deposited in the RCSB (PDB ID 8BW1)

##### 1.10 Evolutionary analysis of ATC tridomains (XUs) from NRPS using PhyML_Multi

The amino acid sequence of NRPS were collected from our *Photorhabdus* and *Xenorhabdus* genome collection. We also included a few NRPS representatives from actinomycetes, cyanobacteria and other proteobacteria in our analysis (sup. x). XUs from NRPS protein sequences were extracted from our NRPS dataset using local BLAST with the second XU from GxpS of Photorhabdus laumondii TT01 as query. XUs were aligned using MUSCLE v3.8.31[9] and trimed with trimAl v1.2[10]. This alignment was used for the evolutionary analysis using the software PhyML_Multi. We specified that PhyML_multi search for two trees under a hidden markov model that together best fit the alignment. Since PhyML_Multi does not have a model finder, the model finder of IQ-tree[10] with the selection of '-msub nuclear' was used. IQ-tree chose JTT[11] as the best fit model which was also used for the analysis with PhyML_Multi with a 4-category gamma distribution of among site rate-variation. Afterwards, the log likelihood of tree 1 was deducted from the log likelihood of tree 2 and plotted.

##### 1.11 Evolutionary analysis of the T domain from NRPS using PhyML_Multi

The T domain dataset covered the amino acid sequence of the A-T-Linker and the T domain. This area was extracted from our NRPS dataset using local BLAST with the third T domain from GxpS of Photorhabdus laumondii TT01 as query. The T domains were aligned using MUSCLE and carefully trimmed manually to reduce gaps. Afterwards, the software PhyML_Multi was used to detect recombination breakpoints and phylogenetic histories within the T domain.

##### 1.12 Topological comparison of different phylogenetic trees

The four different trees generated by PhyML_Multi were pruned using the software mesquite[12] to reduce the number branches on the trees for visual clarity. Trees were compared using the R package phytools[13].

**Table S1.** Strains used in this work.

| Strain | Genotype/NRPS | Reference |
|---|---|---|
| *E. coli* DH10B | F_mcrA (*mrr*-*hsd*RMS-*mcr*BC), 80*lac*ZΔ, M15, Δ*lac*X74 *rec*A1 *end*A1 *ara*D 139Δ(*ara*, *leu*)7697 *gal*U *gal*K λ *rps*L (*Strr*) *nup*G / - | [14] |
| *E. coli* DH10B::*mta*A | DH10B with *mtaA* from pCK_*mtaA*Δ*entD* / - | [15] |
| *Bacillus subtilis* subsp. subtilis str. 168 DSM 402 | WT (*srfAB, ppsA*) | DSMZ |
| *M. xanthus* DK1622 | WT (*MchABC*) | [16, 17] |
| *Pseudomonas lurida* sp. MYb11 | WT (v*iscA*) | [18] |
| *Serratia* sp. SCBI | WT (*swrA*) | [19] |
| *S. marcescens* DSM 12481 | WT (*swrW*) | DSMZ |
| *P. luminescens* subsp. laumondii TT01 | WT (*gxpS*, *kolS*) | DSMZ |
| *P. temperata KT122* | WT (4325) | [20] |
| *X. bovienii* SS-2004 | WT (*txlA*) | [21] |
| *X. doucetiae* DSM 17909 | WT (*xabA, prtA*) | DMSZ |
| *X. indica* DSM17382 | WT (*xldS, xtvB, xeyS, XINDV2_09420*) | DSMZ |
| *X. innexi* DSM 16336 | WT (*fitAB*[*1]) | DSMZ |
| *X. mauleonii* DSM 17908 | WT (*ftrAB*[*2]) | DSMZ |
| *X. miraniensis* DSM 17902 | WT (*ambS*) | DMSZ |
| *X. nematophila* ATCC19061 | WT (*xtpS*, *PAX*) | ATCC |
| *X. stockiae* DSM 17904 | WT (*xabA*) | DMSZ |
| *X. szentirmaii* DSM 16338 | WT (*xabA*) | DMSZ |
| *Xenorhabdus* sp. KK7.4 | WT (*XEKKV2_12060*) | [21,22] |
| *Chondromyces crocatus Cm c5* DSM 14714 | WT (*cpnD*) | DMSZ |

**Table S2.** Protein and nucleic acid references to data bank used for NRPS-constructs.

| **NRPS-construct** | **GenPept locus/protein ID** | **GenBank** | **locus tag** | **gene** |
|---|---|---|---|---|
| NRPS-8 | PHM30481.1<br>PHM29999 | NIBU01000054.1<br>NIBU01000077.1 | Xinn_03284<br>Xinn_03635 | *fitAB* |
| NRPS-9 | YP_003466710.1 | FN667741 | XBJ1_0775 | *txlA* |
| NRPS-10 | WP_148886166.1 | NZ_VNHN01000062.1 | LY16_RS14705 | *prtA* |
| NRPS-11 | WP_099121989.1 | NZ_NJAH01000014.1 | Xekk_RS12280 | XEKKV2_12060 |
| NRPS-12 | MBC8943736.1 | NKHP01000001.1 | Xind_00118 | XINDV2_09420 |
| NRPS-13 | AIM23801.1 | CP003424.1 | SERRSCBI_21215 | *swrA* |
| NRPS-14 | WP_187681863 | NZ_JACSZU010000009.1 | IAI52_RS13305 | *viscA* |
| NRPS-15 | WP_012987679 | NC_013892.1 | XBJ1_1126 | *xfpS* |
| NRPS-16 | CAB13717.2 | AL009126.3 | BSU18340 | *ppsA* |
| NRPS-19 | PHM39367.1<br>PHM39368 | NITY01000011.1 | Xmau_02974<br>Xmau_02975 | *ftrAB* |
| NRPS-20 | BAD60917.1 | AB193098.2 | AB193098.2 | *swrW* |
| NRPS-17 | ABF89060.1<br>ABF89457.1 | CP000113.1 | MXAN_4077<br>MXAN_4078 | *mchAB* |
| NRPS-18 | PHM40846.1 | NIUA01000001.1 | Xszus_00521 | *xabA* |

**Table S3.** Plasmids and corresponding NRPSs used in this work.

| NRPS | Plasmids | Genotype | Reference |
|---|---|---|---|
| | pCOLA_ara/tacI | ori ColA, kanR, *araC*- $P_{BAD}$, and tacI | [23] |
| | pCK_0401 | ori p15A, cmR, *araC*- $P_{BAD}$, and tacI | [24] |
| | pCK_0407 | ori ColDF13, specR, *araC*- $P_{BAD}$, and tacI; mtaA | This work |
| -1 | pLP23 | ori ColA, $kan^R$, *araC*- $P_{BAD}$, *xabABC*_C1A1-*gxpS*_T3C/E4A4T4C/E5A5T5TE and *tacI* | This work |
| -2 | pLP24 | ori ColA, $kan^R$, *araC*- $P_{BAD}$, *xabABC*_C1A1T$1_{1/2}$-*gxpS*_T$3_{1/2}$C/E4A4T4C/E5A5T5TE and *tacI* | This work |
| -3 | pFP7 | ori ColA, $kan^R$, *araC*- $P_{BAD}$, *xabABC*_C1A1T$1_{1/2}$-*gxpS*_T$3_{1/2}$C/E4A4T4C/E5A5T5TE and *tacI* | This work |
| -4 | pFP8 | ori ColA, $kan^R$, *araC*- $P_{BAD}$, *xabABC*_C1A1T$1_{1/2}$-*gxpS*_T$3_{1/2}$C/E4A4T4C/E5A5T5TE and *tacI* | This work |
| -5 | pFP9 | ori ColA, $kan^R$, *araC*- $P_{BAD}$, *xabABC*_C1A1T$1_{1/2}$-*gxpS*_T$3_{1/2}$C/E4A4T4C/E5A5T5TE and *tacI* | This work |
| -6 | pFP11 | ori ColA, $kan^R$, *araC*- $P_{BAD}$, *xabABC*_C1A1T$1_{1/2}$-*gxpS*_T$3_{1/2}$C/E4A4T4C/E5A5T5TE and *tacI* | This work |
| -7 | pLP31 | ori ColA, $kan^R$, *araC*- $P_{BAD}$, *xabABC*_C1A1T1-*gxpS*_C/E4A4T4C/E5A5T5TE and *tacI* | This work |
| -8 | pCK_0683 | ori p15A, cmR, *araC*- $P_{BAD}$, *fitAB* 6 modular NRPS *X. mauleonii* | This work |
| -9a | pCK_0760 | ori p15A, cmR, *araC*- $P_{BAD}$, *txlA* C1A1 – T1 modules 2-6 *fitAB* | This work |
| -9b | pCK_0761 | ori p15A, cmR, *araC*- $P_{BAD}$, *txlA* C1A1 T$1_{1/2}$ – modules 2-6 *fitAB* | This work |
| -10a | pCK_0762 | ori p15A, cmR, *araC*- $P_{BAD}$, *prtA* C1A1 – T1 modules 2-6 *fitAB* | This work |
| -10b | pCK_0762 | ori p15A, cmR, *araC*- $P_{BAD}$, *prtA* C1A1 T$1_{1/2}$ – modules 2-6 *fitAB* | This work |
| -11a | pCK_0768 | ori p15A, cmR, *araC*- $P_{BAD}$, *xucA** C1A$1^{Val}$ – T1 modules 2-6 *fitAB* | This work |
| -11b | pCK_0768 | ori p15A, cmR, *araC*- $P_{BAD}$, *xucA** C1A$1^{Val}$ T$1_{1/2}$ – modules 2-6 *fitAB* | This work |
| -12a | pCK_0820 | ori p15A, cmR, *araC*- $P_{BAD}$, *xucA** C1A$1^{Ser}$ – T1 modules 2-6 *fitAB* | This work |
| -13a | pCK_0822 | ori p15A, cmR, *araC*- $P_{BAD}$, *xucA** C1A$1^{Leu}$ – T1 modules 2-6 *fitAB* | This work |
| -13b | pCK_0823 | ori p15A, cmR, *araC*- $P_{BAD}$, *viscA* C1A$1^{Leu}$ T$1_{1/2}$ – modules 2-6 *fitAB* | This work |
| -14a | pCK_0824 | ori p15A, cmR, *araC*- $P_{BAD}$, *xucA** C1A$1^{Leu}$ – T1 modules 2-6 *fitAB* | This work |
| -14b | pCK_0825 | ori p15A, cmR, *araC*- $P_{BAD}$, *xucA** C1A$1^{Leu}$ T$1_{1/2}$ – modules 2-6 *fitAB* | This work |
| -15a | pCK_0826 | ori p15A, cmR, *araC*- $P_{BAD}$, *xtpS* C1A$1^{Leu}$ – T1 modules 2-6 *fitAB* | This work |
| -15b | pCK_0827 | ori p15A, cmR, *araC*- $P_{BAD}$, *xtpS* C1A$1^{Leu}$ T$1_{1/2}$ – modules 2-6 *fitAB* | This work |
| -16a | pCK_0828 | ori p15A, cmR, *araC*- $P_{BAD}$, *xtpS* C1A$1^{Leu}$T$1_{1/2}$ – modules 2-6 *fitAB* | This work |
| -19 | pCK_0680 | ori p15A, cmR, *araC*- $P_{BAD}$, *ftrAB* 6 modular WT NRPS | This work |
| -17a | pCK_0868 | ori CloDF13, specR, *araC*-$P_{BAD}$ *mchA*-PKS and tacI | This work |

| | | | |
|---|---|---|---|
| -20b | pCK_0870 | ori p15A, cmR, *araC*- $P_{BAD}$, *swrW* C1A1$^{Ser}$ modules 2-6 *ftrAB* | This work |
| -19a | pCK_0873 | ori p15A, cmR, *araC*- $P_{BAD}$, (*mchA*-PKS *mchB* C1A1MT$^{Thr}$ - modules 2-6 *fitAB* | This work |
| -20b | pSB002 | ori p15A, cmR, *araC*- $P_{BAD}$, *xabA* C1A1$^{Pro}$T1_C2A2$^{Gly}$T2$_{1/2}$ – modules 2-6 *fitAB* | This work |
| -21a | pLS_019 | ori p15A, cm$^R$, araC-$P_{BAD}$ I-*Ceu*I, I-*Sce*I gxps_$A_1T_1C/E_2A_2$_*xabA*_$T_3C_4A_4$_*gxps*_$T_4C/E_5A_5T_5$TE and *tacI-araE* | This work |
| -21b | pLS_191 | ori p15A, cm$^R$, araC-$P_{BAD}$ I-*Ceu*I, I-*Sce*I gxps_$A_1T_1C/E_2A_2T_2^{1/2}$_*xabA*_$T3^{1/2}C_4A_4T_4^{1/2}$_*gxps*_$T4^{1/2}$ $C/E_5A_5T_5$TE and *tacI-araE* | This work |
| -22a | pLS_018 | ori p15A, cm$^R$, araC-$P_{BAD}$ I-*Ceu*I, I-*Sce*I gxps_$A_1T_1C/E_2A_2$_*xlds*_$T_2C_3A_3$_*gxps*_$T_4C/E_5A_5T_5$TE and *tacI-araE* | This work |
| -22b | pLS_017 | ori p15A, cm$^R$, araC-$P_{BAD}$ I-*Ceu*I, I-*Sce*I gxps_$A_1T_1C/E_2A_2T_2^{1/2}$_*xlds*_$T_2^{1/2}C_3A_3T_3^{1/2}$_*gxps*_$T_4^{1/2}C/E_5A_5T_5$TE and *tacI-araE* | This work |
| -23a | pLS_009 | ori p15A, cm$^R$, araC-$P_{BAD}$ I-*Ceu*I, I-*Sce*I gxps_$A_1T_1C/E_2A_2$_*cpnd*_$T_2C_3A_3$_*gxps*_$T_4C/E_5A_5T_5$TE and *tacI-araE* | This work |
| -23b | pLS_008 | ori p15A, cm$^R$, araC-$P_{BAD}$ I-*Ceu*I, I-*Sce*I gxps_$A_1T_1C/E_2A_2T2^{1/2}$_*cpnd*_$T_2^{1/2}C_3A_3T_3^{1/2}$_*gxps*_$T_4^{1/2}$ $C/E_5A_5T_5$TE and *tacI-araE* | This work |
| -24a | pLS_003 | ori p15A, cm$^R$, araC-$P_{BAD}$ I-*Ceu*I, I-*Sce*I gxps_$A_1T_1C/E_2A_2$_*mchC$_A$*_$T_2C_3A_3$_*gxps*_$T_4C/E_5A_5T_5$TE and *tacI-araE* | This work |
| -24b | pLS_002 | ori p15A, cm$^R$, araC-$P_{BAD}$ I-*Ceu*I, I-*Sce*I gxps_$A_1T_1C/E_2A_2T2^{1/2}$_*mchC$_A$*_$T_2^{1/2}C_3A_3T_3^{1/2}$_*gxps*_$T_4^{1/2}C/E_5A_5T_5$TE and *tacI-araE* | This work |
| -25 | pPI16_XUT | ori p15A, *cm$^R$*, *araC-$P_{BAD}$* *xldS*_C1A1T1$_{1/2}$-*xabA*_T1$_{1/2}$C1-*kolS*_A2T2C3-*gxpS*_A2T2$_{1/2}$-*xtvAB*_T2$_{1/2}$Red *tacI* and *araE* | This work |
| -26 | pPI16 | ori p15A, *cm$^R$*, *araC-$P_{BAD}$* *xldS*_C1A1T1$_{1/2}$-*xabA*_T1$_{1/2}$C1-*kolS*_A2T2C3-*gxpS*_A2T2$_{1/2}$-*xtvAB*_T2$_{1/2}$Red *tacI* and *araE* | This work |
| -27 | pPI16_typeII | ori p15A, *cm$^R$*, *araC-$P_{BAD}$* *xldS*_C1A1T1$_{1/2}$-*xabA*_T1$_{1/2}$C1-*kolS*_A2T2C3-*gxpS*_A2T2$_{1/2}$-*xtvAB*_T2$_{1/2}$Red *tacI* and *araE* | This work |
| -28 | pPI16_end | ori p15A, *cm$^R$*, *araC-$P_{BAD}$* *xldS*_C1A1T1$_{1/2}$-*xabA*_T1$_{1/2}$C1-*kolS*_A2T2C3-*gxpS*_A2T2$_{1/2}$-*xtvAB*_T2$_{1/2}$Red *tacI* and *araE* | This work |

**Table S4.** Primer and templates used in this work to generate indicated plasmids. Sizes of the PCR products are depicted below the template.

| Plasmids | Oligo-nucleotides | Sequence (5' to 3'), alternatively restriction enzymes | Template Product size in bp |
|---|---|---|---|
| pLP23 | LP134 | TGGGCTAACAGGAGGAATTCCATGCCTATGTCG TGCAATCG | *X. stockiae* gDNA 3.062 |
| | LP135 | GCTTGGTACTCATGCGTGACTACCGC | |
| | LP132 | CAATCTGCGGTAGTCACGCATGAGTACCAAGC GCCACAAGGGGAAATTG | pJW76 5.347 |
| | LP133 | GAACATTCGGATCAAGTACCGTTAACGCGG | |
| | LP136 | AACGGTACTTGATCCGAATGTTC | pJW76 5.545 |
| | LP137 | GGAATTCCTCCTGTTAGCCC | |
| pLP24 | LP134 | TGGGCTAACAGGAGGAATTCCATGCCTATGTCG TGCAATCG | *X. stockiae* gDNA 3.148 |
| | LP139 | AGAAACTGTCATGTCGGCCAACCTGTTCTAATC CTAATAAACTTTGC | |
| | LP138 | GTTGGCCGACATGACAGTTTCTTTGCC | pJW76 5.251 |
| | LP133 | GAACATTCGGATCAAGTACCGTTAACGCGG | |
| | LP136 | AACGGTACTTGATCCGAATGTTC | pJW76 5.545 |
| | LP137 | GGAATTCCTCCTGTTAGCCC | |
| pFP7 | LP55 | GAGGAATTCCATGCCTATGTCGTGCAATCG | *X. stockiae* gDNA 3.149 |
| | LP60 | CGCCCAAGGCAAAGAAATGGTCACGGCGACCA ACCTG | |
| | LP59 | CCATTTCTTTGCCTTGGGCGGTCAC | pJW76 5.308 |
| | LP44 | GTAAATCACATACGCCAGATGTCGTGAGGTC | |
| | LP43 | CGACATCTGGCGTATGTGATTTACACTTCTG | pJW76 5.487 |
| | LP56 | CGACATAGGCATGGAATTCCTCCTGTTAGC | |
| pFP8 | LP55 | GAGGAATTCCATGCCTATGTCGTGCAATCG | *X. stockiae* gDNA 3.157 |
| | LP62 | CGAGTGACCGCCCAATTCAAAGAAATGGTCAC | |
| | LP61 | TTGAATTGGGCGGTCACTCGCTGTTGGC | pJW76 5.300 |
| | LP44 | GTAAATCACATACGCCAGATGTCGTGAGGTC | |
| | LP43 | CGACATCTGGCGTATGTGATTTACACTTCTG | pJW76 5.487 |
| | LP56 | CGACATAGGCATGGAATTCCTCCTGTTAGC | |
| pFP9 | LP55 | GAGGAATTCCATGCCTATGTCGTGCAATCG | *X. stockiae* gDNA 3.171 |
| | LP64 | CTGACTGCCAGAAGAGAGTCACCACCC | |
| | LP63 | GACTCTCTTCTGGCAGTCAGGATGATCGAACG | pJW76 5.286 |
| | LP44 | GTAAATCACATACGCCAGATGTCGTGAGGTC | |
| | LP43 | CGACATCTGGCGTATGTGATTTACACTTCTG | pJW76 5.487 |
| | LP56 | CGACATAGGCATGGAATTCCTCCTGTTAGC | |
| pFP11 | LP55 | GAGGAATTCCATGCCTATGTCGTGCAATCG | *X. stockiae* gDNA 3.202 |
| | LP66 | CAATCCTATACGACGTATACGGGCAGTCATCTG | |
| | LP65 | CCGTATACGTCGTATAGGATTGGGCCTGTC | pJW76 5.257 |
| | LP44 | GTAAATCACATACGCCAGATGTCGTGAGGTC | |
| | LP43 | CGACATCTGGCGTATGTGATTTACACTTCTG | pJW76 5.487 |
| | LP56 | CGACATAGGCATGGAATTCCTCCTGTTAGC | |
| pLP31 | LP134 | TGGGCTAACAGGAGGAATTCCATGCCTATGTCG TGCAATCG | *X. stockiae* gDNA 3.297 |
| | LP160 | GCTAATTTCACGATGTTCAGTAATAACCTGAGC CAACTC | |
| | LP161 | GTTATTACTGAACATCGTGAAATTAGCGTGCCT G | pJW76 5.110 |
| | LP133 | GAACATTCGGATCAAGTACCGTTAACGCGG | |

| | | | |
|---|---|---|---|
| | LP136 | AACGGTACTTGATCCGAATGTTC | pJW76 5.545 |
| | LP137 | GGAATTCCTCCTGTTAGCCC | |
| pPI16 | 26 | TTTTTGGGCTAACAGGAGGAATTCCATGAATAT GACACGTAACCATACATCC | *X. indica* gDNA 3.098 |
| | 29 | GTGAGTGCCCGCCAAGCTCAAAGAAATGATCG TGGCGACCGACAC | |
| | 12 | TTCTTTGAGCTTGGCGGGC | *X. doucetiae* gDNA 1.536 |
| | AL13-2 | ATCCACCAGCAGTTGTTGTCG | |
| | 40 | GGAGCGACAACAACTGCTGGTGGATTGGAATG CAACCGCAACC | *P. luminescens* subsp. laumondii TT01 gDNA 3.203 |
| | AT_492 | GATAGGGGGTTTCTGTCGCGTTCCAAGTTTCCA ATAACAACTTGCGCTC | |
| | AT_226 | TGGAACGCGACAGAAACC | *P. luminescens* subsp. laumondii TT01 gDNA 1.668 |
| | 9 | ATTATCGTGTCGGCCGATTTGCTC | |
| | 14 | AAATCGGCCGACACGATAATTTTTTCAATATCG GAGGACATTCGC | *X. indica* gDNA 1.383 |
| | 6 | TATACGAGCCGATGATTAATTGTCATTACTTATA TTCCGGTTCATATTTTTTGTCC | |
| | pACYC-2 | TGACAATTAATCATCGGCTCG | pJW75 5.220 |
| | pACYC-1 | GGAATTCCTCCTGTTAGCC | |
| pPI16_XUT | 26 | TTTTTGGGCTAACAGGAGGAATTCCATGAATAT GACACGTAACCATACATCC | *X. indica* gDNA 3.098 |
| | 29 | GTGAGTGCCCGCCAAGCTCAAAGAAATGATCG TGGCGACCGACAC | |
| | 12 | TTCTTTGAGCTTGGCGGGC | *X. doucetiae* gDNA 1.536 |
| | AL13-2 | ATCCACCAGCAGTTGTTGTCG | |
| | 40 | GGAGCGACAACAACTGCTGGTGGATTGGAATG CAACCGCAACC | *P. luminescens* subsp. laumondii TT01 gDNA 3.203 |
| | AT_492 | GATAGGGGGTTTCTGTCGCGTTCCAAGTTTCCA ATAACAACTTGCGCTC | |
| | AT_226 | TGGAACGCGACAGAAACC | *P. luminescens* subsp. laumondii TT01 gDNA 1.578 |
| | LP356 | AATTTGGCGAGCAAAAGCATCC | |
| | LP357 | AGAGGATGCTTTTGCTCGCCAAATTTCTGAGGA ACGTCTGACTTC | *X. indica* gDNA 1.478 |
| | 6 | TATACGAGCCGATGATTAATTGTCATTACTTATA TTCCGGTTCATATTTTTTGTCC | |
| | pACYC-2 | TGACAATTAATCATCGGCTCG | pJW75 5.220 |
| | pACYC-1 | GGAATTCCTCCTGTTAGCC | |
| pPI16_type II | 26 | TTTTTGGGCTAACAGGAGGAATTCCATGAATAT GACACGTAACCATACATCC | *X. indica* gDNA 3.098 |
| | 29 | GTGAGTGCCCGCCAAGCTCAAAGAAATGATCG TGGCGACCGACAC | |
| | 12 | TTCTTTGAGCTTGGCGGGC | *X. doucetiae* gDNA 1.536 |
| | AL13-2 | ATCCACCAGCAGTTGTTGTCG | |

| | | | |
|---|---|---|---|
| | 40 | GGAGCGACAACAACTGCTGGTGGATTGGAATG CAACCGCAACC | *P. luminescens* subsp. laumondii TT01 gDNA 3.203 |
| | AT_492 | GATAGGGGGTTTCTGTCGCGTTCCAAGTTTCCA ATAACAACTTGCGCTC | |
| | AT_226 | TGGAACGCGACAGAAACC | *P. luminescens* subsp. laumondii TT01 gDNA 1.680 |
| | LP358 | CAAGGCAAAAAAATTATCGTGTCGGC | |
| | LP359 | CCGACACGATAATTTTTTTGCCTTGGGAGGACA TTCGCTATTAGC | *X. indica* gDNA 1.376 |
| | 6 | TATACGAGCCGATGATTAATTGTCATTACTTATA TTCCGGTTCATATTTTTTGTCC | |
| | pACYC-2 | TGACAATTAATCATCGGCTCG | pJW75 5.220 |
| | pACYC-1 | GGAATTCCTCCTGTTAGCC | |
| pPI16_end | 26 | TTTTTGGGCTAACAGGAGGAATTCCATGAATAT GACACGTAACCATACATCC | *X. indica* gDNA 3.098 |
| | 29 | GTGAGTGCCCGCCAAGCTCAAAGAAATGATCG TGGCGACCGACAC | |
| | 12 | TTCTTTGAGCTTGGCGGGC | *X. doucetiae* gDNA 1.536 |
| | AL13-2 | ATCCACCAGCAGTTGTTGTCG | |
| | 40 | GGAGCGACAACAACTGCTGGTGGATTGGAATG CAACCGCAACC | *P. luminescens* subsp. laumondii TT01 gDNA 3.203 |
| | AT_492 | GATAGGGGGTTTCTGTCGCGTTCCAAGTTTCCA ATAACAACTTGCGCTC | |
| | AT_226 | TGGAACGCGACAGAAACC | *P. luminescens* subsp. laumondii TT01 gDNA 1.803 |
| | LP360 | TGCGCAGATTTTCTCGGTAAATGTCGCC | |
| | LP361 | GACATTTACCGAGAAAATCTGCGCATATCTGAA TAATAATCAAAAAAACAATAACGAAATG | *X. indica* gDNA 1.250 |
| | 6 | TATACGAGCCGATGATTAATTGTCATTACTTATA TTCCGGTTCATATTTTTTGTCC | |
| | pACYC-2 | TGACAATTAATCATCGGCTCG | pJW75 5.220 |
| | pACYC-1 | GGAATTCCTCCTGTTAGCC | |
| pCK_0678 | ck002<br>ck0467<br><br>ck0436b<br><br>ck0468<br>ck0465<br><br>ck0468 | CATGGAATTCCTCCTGTTAG<br>CATCAGGATATGTTAATTAACCTAGGCTGCTGC CAC<br>AGGAATTCCATGACAAAATCTGAATATTTAGTAA GTTCA<br>GAATTGTCAGAAACCTACCAAGCTTTGCG<br>CCTAGGTTAATTAACATATCCTGATGGGCTTTG GCTCCTG<br>TTCCCGCAAAGCTTGGTAGGTTTCTGAC | pCK_0401 3.672<br><br>*X. mauleonii* gDNA 3.773<br>*X. mauleonii* gDNA 6.992 |
| pCK_0679 | <br>ck0459<br>ck0460<br><br>ck0471<br><br>ck0472 | MluI/SnaBI<br>CAAAGCGGGACCAAAGCCATG<br>TGAGACCTTTTTTTGGTCTCGGAATTCCTCCTGT TAG<br>CCGAGACCAAAAAAAGGTCTCACCCCTTGAATA CAAGGCGTTGC<br>CCCGTTCGCTGGGATATTCTGG | pCK_0678 12.499<br>pCK_0401 230<br><br>pCK_0678 365 |

| | | | |
|---|---|---|---|
| pCK_0680<br>NRPS-19 | <br><br>ck0469b<br><br>ck0470 | NcoI/PacI<br><br>GGTGGCAGCAGCCTAGGTTAATTAACTGGCTTT<br>ATTAAGAT-<br>ACCTCAAGAAAACCCCAGCCCCTGATAGGTATG<br>TTTG | pCK_0678<br>14.247<br>*X. mauleonii*<br>gDNA<br>12.632 |
| pCK_0681 | <br><br>ck0594<br>ck0463b<br><br>ck0459<br>ck0460 | MluI/AscI<br><br>GGCACCACCGATATACAGTTCACC<br>AGGAATTCCATGACAAAATCTGAATATTTAGTAA<br>GTTCA<br>CAAAGCGGGACCAAAGCCATG<br>TGAGACCTTTTTTTGGTCTCGGAATTCCTCCTGT<br>TAG | pCK_0680<br>12.905<br>pCK_0680<br>2.520<br>pCK_0401<br>230 |
| pCK_0682 | ck0455<br><br>ck0456<br><br>ck0451<br><br>ck0452<br>ck0453<br>ck0454 | CTGTGATATCAGCCAATTAATTAACCTAGGCTG<br>CTGCCAC<br>GATCTCATGGAATTCCTCCTGTTAGCCCA<br><br>TTTGGGCTAACAGGAGGAATTCCATGAGATCAT<br>TTGAG-GATTCACTGA<br>GGGTCTTTAGACCACCCGATTGC<br>GCGCAATCGGGTGGTCTAAAGAC<br>CTAGGTTAATTAATTGGCTGATATCACAGTGCT<br>GTAATGG | pCK_0401<br>3.681<br><br><br>*X. innexi* gDNA<br>10.179<br><br>*X. innexi* gDNA<br>3.904 |
| pCK_0683<br>NRPS-8 | <br><br><br>ck0457<br>ck0522 | BglII/AvrII<br><br><br>GAACCAAACAGGGTTATCGTCAGTGC<br>TGCTCAGCGGTGGCAGCAGCCTAGGTTAATTTA<br>CGCCAATACCTTTTCCTGAC | pCK_0682<br>17.584<br><br>*X. innexi* gDNA<br>8.441 |
| pCK_0684 | <br><br>ck0454<br><br>ck0460<br><br><br>ck0461b<br>ck0462 | AvrII/AscI<br><br>CTAGGTTAATTAATTGGCTGATATCACAGTGCT<br>GTAATGG<br>TGAGACCTTTTTTTGGTCTCGGAATTCCTCCTGT<br>TAG<br>CGAGACCAAAAAAAGGTCTCAGCCCCTTATCCG<br>CAGGATAAAC<br>TGTCATCAGATGATGCGCCAGTTGG | pCK_0682<br>12.549<br>pCK_0682<br>3693<br><br><br>pCK_0682<br>174 |
| pCK_0685 | <br><br>ck0457<br>ck0523 | BglII/AvrII<br><br>GAACCAAACAGGGTTATCGTCAGTGC<br>TATTGCTCAGCGGTGGCAGCAGCCTAGGTTAAT<br>TTACGCCAATACCTTTTCCTGAC | pCK_0684<br>16.242<br>*X. innexi* gDNA<br>8.444 |
| pCK_0760<br>NRPS-9a | <br><br>ck0618<br>ck0592<br><br>ck0475<br><br>ck0635 | BsaI/AatII<br><br>TATGTTGCCCCCCGTAACGCA<br>AATATAAGCAGCCATATCGCTGAGCG<br><br>TTTTTTTGGGCTAACAGGAGGAATTCAATGAGA<br>ACATCTGAAAGCTCGTTG<br>TGCGTTACGGGGGGCAACATAACCGTCCCGGT<br>TTCCCCA | pCK_0685<br>20.443<br>pCK_0683<br>2.658<br><br>*X. bovienii* gDNA<br>3.003 |
| pCK_0761<br>NRPS-9b | | BsaI/AatII | pCK_0685<br>20.443 |

| | | | |
|---|---|---|---|
| | ck0617<br>ck0592<br><br>ck0475<br><br>ck0636 | AATTCCCTGAGCGCCATTAAGCTG<br>AATATAAGCAGCCATATCGCTGAGCG<br><br>TTTTTTTGGGCTAACAGGAGGAATTCAATGAGA<br>ACATCTGAAAGCTCGTTG<br>GCTTAATGGCGCTCAGGGAATTTCCCCCGATCC<br>GGAAAAAGTTA | pCK_0683<br>2.550<br><br>*X. bovienii* gDNA<br>3.112 |
| pCK_0762<br>NRPS-10a | <br><br>ck0618<br>ck0592<br>ck0477<br><br>ck0637 | BsaI/AatII<br><br>TATGTTGCCCCCCGTAACGCA<br>AATATAAGCAGCCATATCGCTGAGCG<br>TTTTTTTGGGCTAACAGGAGGAATTCAATGAGA<br>ATACCTGAAGGTTCGT<br>TGCGTTACGGGGGGCAACATAACTGTCCCGGT<br>TTTCCCATACG | pCK_0685<br>20.443<br>pCK_0683<br>2.658<br>*X. doucetiae*<br>gDNA<br>2.997 |
| pCK_0763<br>NRPS-10b | <br><br>ck0617<br>ck0592<br>ck0477<br><br>ck0638 | BsaI/AatII<br><br>AATTCCCTGAGCGCCATTAAGCTG<br>AATATAAGCAGCCATATCGCTGAGCG<br>TTTTTTTGGGCTAACAGGAGGAATTCAATGAGA<br>ATACCTGAAGGTTCGT<br>GCTTAATGGCGCTCAGGGAATTGCCACCGATA<br>CGGAAAAAATTATCC | pCK_0685<br>20.443<br>pCK_0683<br>2.550<br>*X. doucetiae*<br>gDNA<br>3.106 |
| pCK_0768<br>NRPS-11a | <br><br>ck0618<br>ck0592<br>ck0487<br><br>ck0648 | BsaI/AatII<br><br>TATGTTGCCCCCCGTAACGCA<br>AATATAAGCAGCCATATCGCTGAGCG<br>TTTTTTTGGGCTAACAGGAGGAATTCAATGAGA<br>AAAGCTGAGGATCATTTGAA<br>TGCGTTACGGGGGGCAACATAACTGTCTCTGTT<br>GCCGAAAGC | pCK_0685<br>20.443<br>pCK_0683<br>2.658<br>X. sp. KK7.4<br>gDNA<br>2.943 |
| pCK_0769<br>NRPS-11b | <br><br>ck0617<br>ck0592<br><br>ck0487<br><br>ck0649 | BsaI/AatII<br><br>AATTCCCTGAGCGCCATTAAGCTG<br>AATATAAGCAGCCATATCGCTGAGCG<br><br>TTTTTTTGGGCTAACAGGAGGAATTCAATGAGA<br>AAAGCTGAGGATCATTTGAA<br>GCTTAATGGCGCTCAGGGAATTGCCGCCGATA<br>CGGAAGAAATTATC | pCK_0685<br>20.443<br>pCK_0683<br>2.550<br><br>X. sp. KK7.4<br>gDNA<br>3.052 |
| pCK_0820<br>NRPS-12a | <br><br>ck0618<br>ck0592<br>ck0708<br><br>ck0717 | BsaI/AatII<br><br>TATGTTGCCCCCCGTAACGCA<br>AATATAAGCAGCCATATCGCTGAGCG<br>TTTTTTTGGGCTAACAGGAGGAATTCAATGAAT<br>CACCCTGAAAATATGAAAC<br>TGCGTTACGGGGGGCAACATATTCTTGTGTGAT<br>TACTGCTGAATG | pCK_0685<br>20.443<br>pCK_0683<br>2.658<br>*X. indica*<br>gDNA<br>3.030 |
| pCK_0822<br>NRPS-13a | <br><br>ck0618<br>ck0592<br><br>ck0711<br><br>ck0719 | BsaI/AatII<br><br>TATGTTGCCCCCCGTAACGCA<br>AATATAAGCAGCCATATCGCTGAGCG<br><br>TTTTTTTGGGCTAACAGGAGGAATTCAATGAAC<br>AAACAAACTGATGTGAAGAG | pCK_0685<br>20.443<br>pCK_0683<br>2.658<br><br>Serratia sp. SCBI<br>gDNA<br>4011 |

| | | | |
|---|---|---|---|
| | | TGCGTTACGGGGGGCAACATAGTTTTCACGCAT GGCGGC | |
| pCK_0823<br>NRPS-13b | <br><br>ck0617<br>ck0592<br>ck0711<br><br>ck0720 | BsaI/AatII<br><br>AATTCCCTGAGCGCCATTAAGCTG<br>AATATAAGCAGCCATATCGCTGAGCG<br>TTTTTTTGGGCTAACAGGAGGAATTCAATGAAC AAACAAACTGATGTGAAGAG<br>GCTTAATGGCGCTCAGGGAATTACCGCCCAACT CGAAGAAG | pCK_0685<br>20.443<br>pCK_0683<br>2.550<br>Serratia sp. SCBI<br>gDNA<br>4120 |
| pCK_0824<br>NRPS-14a | <br><br>ck0618<br>ck0592<br>ck0714<br><br>ck0721 | BsaI/AatII<br><br>TATGTTGCCCCCCGTAACGCA<br>AATATAAGCAGCCATATCGCTGAGCG<br>TTTTTTTGGGCTAACAGGAGGAATTCAATGAAG CATTCCACCCGCC<br>TGCGTTACGGGGGGCAACATACAGGCGAGTGA CGAAGGC | pCK_0685<br>20.443<br>pCK_0683<br>2.658<br>*P. lurida*<br>gDNA<br>2.877 |
| pCK_0825<br>NRPS-14b | <br><br>ck0617<br>ck0592<br>ck0714<br><br>ck0722 | BsaI/AatII<br><br>AATTCCCTGAGCGCCATTAAGCTG<br>AATATAAGCAGCCATATCGCTGAGCG<br>TTTTTTTGGGCTAACAGGAGGAATTCAATGAAG CATTCCACCCGCC<br>GCTTAATGGCGCTCAGGGAATTCCCGCCGAGT TCAAAGAAG | pCK_0685<br>20.443<br>pCK_0683<br>2.550<br>*P. lurida*<br>gDNA<br>2.986 |
| pCK_0826<br>NRPS-15a | <br><br>ck0618<br>ck0592<br>ck0723<br><br>ck0729 | BsaI/AatII<br><br>TATGTTGCCCCCCGTAACGCA<br>AATATAAGCAGCCATATCGCTGAGCG<br>TTTTTTTGGGCTAACAGGAGGAATTCAATGGAT AACATTCTGGCCTCG<br>TGCGTTACGGGGGGCAACATAAACATAGCGGC TCTGTTTAAAATC | pCK_0685<br>20.443<br>pCK_0683<br>2.658<br>*X. bovienii* gDNA<br>2.877 |
| pCK_0827<br>NRPS-15b | <br><br>ck0617<br>ck0592<br>ck0723<br><br>ck0730 | BsaI/AatII<br><br>AATTCCCTGAGCGCCATTAAGCTG<br>AATATAAGCAGCCATATCGCTGAGCG<br>TTTTTTTGGGCTAACAGGAGGAATTCAATGGAT AACATTCTGGCCTCG<br>GCTTAATGGCGCTCAGGGAATTGCCCCCCAGA TGAAAAAAGT | pCK_0685<br>20.443<br>pCK_0683<br>2.550<br>*X. bovienii* gDNA<br>2.986 |
| pCK_0828<br>NRPS-16 | <br><br>ck0618<br>ck0592<br>ck0726<br><br>ck0731 | BsaI/AatII<br><br>TATGTTGCCCCCCGTAACGCA<br>AATATAAGCAGCCATATCGCTGAGCG<br>TTTTTTTGGGCTAACAGGAGGAATTCAATGAGC GAACATACTTATTCTTTAACC<br>TGCGTTACGGGGGGCAACATAGTAGGTCTCCG CATCTGC | pCK_0685<br>20.443<br>pCK_0683<br>2.658<br>*B. subtilis* 168<br>gDNA<br>2.925 |
| pCK_0868 | ck0828<br>ck0829<br>ck0785b<br><br>ck0867 | TTAATTAACCTAGGCTGCTGCCACC<br>CATTGAATTCCTCCTGTTAGCCCAAAAAAACG<br>TTTTTTTGGGCTAACAGGAGGAATTCAATGAGC GCAGTGTCCAATATTGA<br>ACTTCCGCTTCGGGAAGGACAATCT | pCK_0406<br>3.173<br>*M. xanthus* gDNA<br>3.281 |

| | | | |
|---|---|---|---|
| | ck0866<br>ck0868 | AGATTGTCCTTCCCGAAGCGGAAGT<br>TGGCAGCAGCCTAGGTTAATTAATGGTGTACTC<br>ATGCTGTCTCCCTCT | *M. xanthus* gDNA<br>3.261 |
| pCK_0870<br>NRPS-20b<br><br><br><br><br><br> | <br><br>ck0787<br>ck0788<br>ck0820<br><br><br>ck0822 | BsaI/AatII<br><br>GGCGGCAATTCCCTGATGG<br>GCATTGAAGAATTTTTCTTGTGCAGC<br>TTTTTTTGGGCTAACAGGAGGAATTCAATGTCC<br>GCTTATTCCCTGACGA<br>TAGCCATCAGGGAATTGCCGCCCAGCGCGAAG<br>AA | pCK_0681<br>21.482<br>pCK_0680<br>2135<br>*S. marcescens*<br>*gDNA*<br>3.055 |
| pCK_0873 | <br><br>ck0870<br><br>ck0798<br><br>ck0790<br>ck0592 | BsaI/AatII<br><br>TTGGGCTAACAGGAGGAATTCAATGAGTACACC<br>AGCTGACAACATGAA<br>TTCCTGTGCGTTACGGGGGGCAACGTAGGCCG<br>TCTCCAGG<br>GTTGCCCCCCGTAACGCA<br>AATATAAGCAGCCATATCGCTGAGCG | pCK_0685<br>20.451<br>*M. xanthus* gDNA<br>4.375<br><br>pCK_0683<br>2.665 |
| pSB002<br>NRPS-18b | <br><br>ck0617<br>ck0592<br>SB001<br><br>SB003 | BsaI/AatII<br><br>AATTCCCTGAGCGCCATTAAGCTG<br>AATATAAGCAGCCATATCGCTGAGCG<br>TTTTTTTGGGCTAACAGGAGGAATTCCATGTCT<br>ATGTCATGT-CACCGTATTAACAACG<br>GCTTAATGGCGCTCAGGGAATTCCCGCCCAGC<br>TCAAAGAAATG | pCK_0685<br>20.443<br>pCK_0683<br>2.550<br>*X. szentirmaii*<br>gDNA<br>6.490 |
| pCK_0881 | ck0828<br>ck0921<br>ck0857<br><br>ck0886<br><br>ck0873<br><br>ck0874 | TTAATTAACCTAGGCTGCTGCCACC<br>CATTGAATTCCTCCTGTTAGCCCAAA<br>CGAGACCAAAGAAGAAGGTCTCAGCTGCACCG<br>CAAGGAGAAACCGAAAC<br>TTGCTCAGCGGTGGCAGCAGCCTAGGTTAATTA<br>ATTACAGCGCCTCCGCTTCACAATTCATTG<br>TTTGGGCTAACAGGAGGAATTCAATGAAAGATA<br>GCATGGCTAAAAAGGAA<br>TGAGACCTTCTTCTTTGGTCTCGATAAATTTGGC<br>GAGCAAAAGCATC | pCK_0401<br>3.669<br>*P. luminescens*<br>subsp. laumondii<br>TT01 gDNA<br>4.365<br>*P. luminescens*<br>subsp. laumondii<br>TT01 gDNA<br>4.993 |
| pCK_0882 | ck0828<br>ck0921<br>ck0860<br><br>ck0886<br><br>ck0873<br><br>ck0875 | TTAATTAACCTAGGCTGCTGCCACC<br>CATTGAATTCCTCCTGTTAGCCCAAA<br>CGAGACCAAAGAAGAAGGTCTCAGGTGGCCAT<br>TCGTTGCTTGCGG<br>TTGCTCAGCGGTGGCAGCAGCCTAGGTTAATTA<br>ATTACAGCGCCTCCGCTTCACAATTCATTG<br>TTTGGGCTAACAGGAGGAATTCAATGAAAGATA<br>GCATGGCTAAAAAGGAA<br>TGAGACCTTCTTCTTTGGTCTCGCAAGGCAAAA<br>AAATTATCGTGTCGG | pCK_0401<br>3.669<br>*P. luminescens*<br>subsp. laumondii<br>TT01 gDNA<br>4.266<br>*P. luminescens*<br>subsp. laumondii<br>TT01 gDNA<br>5.092 |
| pLS_002 | <br><br><br>ls06<br><br>ls07 | BsaI<br><br><br>GCCGACACGATAATTTTTTTGCCTTGGGCGGGC<br>ACTCGCTGCTCGCGAT<br>TACCGCAAGCAACGAATGGCCCCCCAAGTCGA<br>AGAAGTTGTCCTCCGCG | pCK_0882<br>12.921<br><br>*M. xanthus* gDNA<br>3.194 |
| pLS_003 | | BsaI | pCK_0881 |

| | | | |
|---|---|---|---|
| | <br>ls08<br><br>ls09 | <br>GCTTTTGCTCGCCAAATTTATGAGCCGCCTCGC ACGCCTA<br>TTCGGTTTCTCCTTGCGGTGCAGCGAAGCGCG TCTCGCTCGCG | 12.921<br>*M. xanthus* gDNA<br>3.189 |
| pLS_008 | <br>ls24<br><br>ls25 | BsaI<br>CGACACGATAATTTTTTTGCCTTGGGCGGCCAC TCCTTGCTGGC<br>ACCGCAAGCAACGAATGGCCCCCCAGCGCGAA GAAGTCGTCCTGC | pCK_0882<br>12.921<br>*C. crocatus* gDNA<br>3.200 |
| pLS_009 | <br>ls26b<br><br>ls27b | BsaI<br>GGATGCTTTTGCTCGCCAAATTTATGTCACGCC CCGCACGCC<br>GGTTTCGGTTTCTCCTTGCGGTGCAGCGAACTC GAAAGCTCCCTCGGCA | pCK_0881<br>12.921<br>*C. crocatus* gDNA<br>3.205 |
| pLS_017 | <br>ls52<br><br>ls53 | BsaI<br>CCGACACGATAATTTTTTTGCCTTGGGTGGCCA TTCATTACTCGCTG<br>TACCGCAAGCAACGAATGGCCACCGAGTTCGA AGAAGTGGTCATAACG | pCK_0882<br>12.921<br>*X. indica* gDNA<br>3.241 |
| pLS_018 | <br>ls68<br><br>ls55 | BsaI<br>GCTTTTGCTCGCCAAATTTATGAAGCGCCCATT GGCAAATTGGAA<br>CGGTTTCTCCTTGCGGTGCAGCATAGCCACGT GTAACAACCGCTG | pCK_0881<br>12.921<br>*X. indica* gDNA<br>3.241 |
| pLS_019 | <br>ls60<br><br>ls61 | BsaI<br>GAGGATGCTTTTGCTCGCCAAATTTATCAAGCG CCGGAAAGCCCAATGGA<br>GGTTTCGGTTTCTCCTTGCGGTGCAGCATATTG ACTCAATACAAACGCGGATGGC | pCK_0882<br>12.921<br>*X. mauleonii* gDNA<br>3.288 |
| pLS_0191 | ls71_1<br>ls74_1<br><br>ls73<br>ls72_1<br><br>ls62<br><br>ls63 | GGTGGCCATTCGTTGCTTGCG<br>CAGGTGCTACATTTGAAGAGATAAATTGC<br><br>CTCTTCAAATGTAGCACCTGAAGTCAGC<br>CAAGGCAAAAAAATTATCGTGTCGGCC<br><br>CGGCCGACACGATAATTTTTTTGCCTTGGGCGG CCATTCATTGCTTG<br>CGTACCGCAAGCAACGAATGGCCACCCAATTC AAAGAAATGATCATGGCGAC | pCK_0882<br>6.395<br><br>pCK_0882<br>6.546<br><br>*X. mauleonii* gDNA<br>3.288 |

**Table S5.** Detected compounds in this work.

| Peptide | MS detected $[M+H]^+$ | MS calculated $[M+H]^+$ | Molecular ion formula | Δppm | Reference |
|---|---|---|---|---|---|
| **1** | 444.3061 | 444.3068 | $C_{22}H_{42}N_3O_6$ | 1.5 | synthetic |
| **1** | 444.3062 | 444.3068 | $C_{22}H_{42}N_3O_6$ | 1.3 | |
| **2** | 430.2911 | 430.2912 | $C_{21}H_{40}N_3O_6$ | 0.1 | |
| **3** | 416.2750 | 416.2755 | $C_{20}H_{38}N_3O_6$ | 1.3 | |
| **4**, **5** | 767.3932 | 767.3974 | $C_{39}H_{55}N_6O_{10}$ | 5.5 | isolated NP |
| **6** | 783.3912 | 783.3923 | $C_{39}H_{54}N_6O_{11}$ | 1.4 | |
| **7** | 811.4217 | 811.4236 | $C_{41}H_{58}N_6O_{11}$ | 2.4 | |
| **8** | 839.4531 | 839.4549 | $C_{43}H_{62}N_6O_{11}$ | 2.2 | |
| **9** | 783.3912 | 783.3923 | $C_{39}H_{54}N_6O_{11}$ | 0.9 | |
| **10** | 811.4219 | 811.4236 | $C_{41}H_{58}N_6O_{11}$ | 2.1 | |
| **11** | 839.4531 | 839.4549 | $C_{43}H_{62}N_6O_{11}$ | 2.2 | |
| **12** | 782.4071 | 782.4083 | $C_{39}H_{55}N_7O_{10}$ | 1.6 | |
| **13** | 810.4375 | 810.4397 | $C_{41}H_{59}N_7O_{10}$ | 2.6 | |
| **14** | 838.4680 | 838.4710 | $C_{43}H_{63}N_7O_{10}$ | 3.5 | |
| **15** | 753.3808 | 753.3818 | $C_{38}H_{52}N_6O_{10}$ | 1.3 | |
| **16** | 869.4631 | 869.4655 | $C_{44}H_{64}N_6O_{12}$ | 2.8 | |
| **17** | 867.4833 | 867.4862 | $C_{45}H_{66}N_6O_{11}$ | 3.4 | |
| **18** | 895.5143 | 895.5175 | $C_{47}H_{70}N_6O_{11}$ | 3.6 | |
| **19** | 725.3855 | 725.3869 | $C_{37}H_{52}N_6O_9$ | 1.9 | |
| **20** | 993.5499 | 993.5543 | $C_{52}H_{76}N_6O_{13}$ | 4.4 | |
| **21** | 995.5662 | 995.5700 | $C_{52}H_{78}N_6O_{13}$ | 3.8 | |
| **22** | 977.5557 | 977.5594 | $C_{52}H_{76}N_6O_{12}$ | 3.8 | |
| **23** | 979.5715 | 979.5751 | $C_{52}H_{78}N_6O_{12}$ | 3.6 | |
| **24** | 955.4986 | 977.5019 | $C_{54}H_{68}N_6O_{11}$ | 3.3 | |
| **25** | 836.4167 | 836.4888 | $C_{42}H_{57}N_7O_{11}$ | 2.6 | |
| **26** | 799.4429 | 799.4461 | $C_{38}H_{58}N_{10}O_9$ | 4.0 | Isolated NP |
| **27** | 813.4579 | 813.4618 | $C_{39}H_{60}N_{10}O_9$ | 4.7 | |
| **28** | 913.5475 | 913.5506 | $C_{45}H_{72}N_{10}O_{10}$ | 3.4 | |
| **29** | 941.5780 | 941.5819 | $C_{47}H_{76}N_{10}O_{10}$ | 4.1 | |
| **30** | 931.5573 | 931.5611 | $C_{45}H_{74}N_{10}O_{11}$ | 4.1 | |
| **31** | 945.5728 | 945.5768 | $C_{46}H_{76}N_{10}O_{11}$ | 4.2 | |
| **32** | 959.5892 | 959.5924 | $C_{47}H_{78}N_{10}O_{11}$ | 3.4 | |
| **33** | 973.6033 | 973.6080 | $C_{48}H_{80}N_{10}O_{11}$ | 4.9 | |
| **34** | 457.3378 | 457.3384 | $C_{23}H_{44}N_4O_5$ | 1.3 | synthetic |
| **35** | 471.3534 | 471.3541 | $C_{24}H_{46}N_4O_5$ | 1.4 | |
| **36** | 431.2857 | 431.2864 | $C_{20}H_{39}N_4O_6$ | 1.6 | synthetic |
| **37** | 445.3010 | 445.3021 | $C_{21}H_{41}N_4O_6$ | 1.7 | |
| **38** | 415.2910 | 415.2915 | $C_{20}H_{39}N_4O_5$ | 1.2 | synthetic |
| **39** | 429.3064 | 429.3071 | $C_{21}H_{41}N_4O_5$ | 1.8 | |
| **40** | 511.3845 | 511.3854 | $C_{27}H_{51}N_4O_5$ | 1.7 | |
| **41** | 525.3998 | 525.4010 | $C_{28}H_{53}N_4O_5$ | 2.3 | |
| **42** | 539.4159 | 539.4167 | $C_{29}H_{55}N_4O_5$ | 1.5 | |
| **43** | 458.3218 | 458.3225 | $C_{23}H_{44}N_3O_6$ | 1.5 | |

**Table S6.** $^1$H (500 MHz) and $^{13}$C (125 MHz) NMR data of compound **1** in DMSO-d$_6$ ($\delta$ in ppm). COSY (bold) and key HMBC (arrows) are shown.

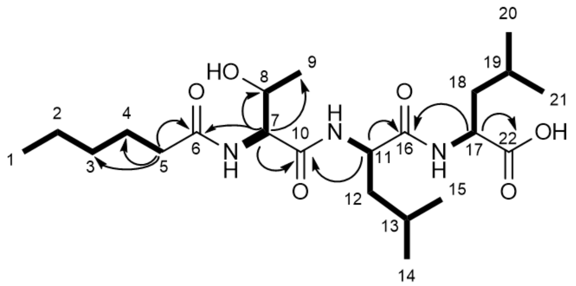


| Position | $\delta_C$, type$^a$ | $\delta_H$, mult. ($J$ in Hz) |
|---|---|---|
| 1 | 13.86, $CH_3$ | 0.88-0.79, ov |
| 2 | 21.90, $CH_2$ | 1.31-1.81, m |
| 3 | 30.85, $CH_2$ | 1.31-1.81, m |
| 4 | 25.00, $CH_2$ | 1.64-1.42, m |
| 5 | 35.10, $CH_2$ | 2.17, m |
| 6 | 172.54, C | - |
| 7 | 57.85, CH | 4.26, dd (12.0, 6.0) |
| 7NH | - | 7.70, d (8.32) |
| 8 | 66.44, CH | 3.95, m |
| 9 | 19.39, $CH_3$ | 1.01, d (6.34) |
| 10 | 169.76, C | - |
| 11 | 50.90, CH | 4.35, dd(15.0, 8.4) |
| 11NH | - | 7.81, d (8.53) |
| 12 | 41.01, $CH_2$ | 1.64-1.42, ov |
| 13 | 24.27 – 24.03, CH | 1.64-1.42, ov |
| 14 | - | 0.88-0.79, ov |
| 15 | - | 0.88-0.79, ov |
| 16 | 171.82, C | - |
| 17 | 50.10, CH | 4.20, ddd (10.0, 8.3, 4.8)f |
| 17NH | - | 8.03, d (8.19) |
| 18 | 40.04, $CH_2$ | 1.64-1.42, ov |
| 19 | 24.27 – 24.03, CH | 1.64-1.42, ov |
| 20 | - | 0.88-0.79, ov |
| 21 | - | 0.88-0.79, ov |
| 22 | 173.92, C | - |

**Table S7**. $^{1}H$ (500 MHz) and $^{13}C$ (125 MHz) NMR spectroscopic data of compounds **4** and **5** in DMSO-$d_6$ ($\delta$ in ppm and $J$ in Hz).

| no. | **4** | | **5** | |
|---|---|---|---|---|
| | $\delta_C$, type | $\delta_H$ (mult., $J$) | $\delta_C$, type | $\delta_H$ (mult., $J$) |
| 1 | 169.4 | | 169.4 | |
| 2 | | 6.69 (d, 5.7) | | 7.06 (d, 5.7) |
| 3 | 37.1 | 3.46 (td, 13.0, 5.6) | 36.9 | 3.46 (td, 12.8, 5.8) |
| | | 3.31 (m) | | 3.34 (m) |
| 4 | 34.3 | 2.52 (m) | 34.4 | 2.54 (m) |
| | | 2.23 (m) | | 2.22 (m) |
| 5 | 173.3 | | 173.6 | |
| 6 | | 8.50 (d, 4.4) | | 8.38 (d, 5.1) |
| 7 | 57.4 | 4.25 (m) | 57.4 | 4.30 (m) |
| 8 | 35.3 | 2.61 (dd, 14.4, 4.8) | 35.3 | 2.64 (m) |
| | | 2.34 (dd, 14.4, 3.7) | | 2.37 (dd, 14.0, 2.2) |
| 9 | 127.8 | | 128.0 | |
| 10 | 130.2 | 7.04 (d, 8.3) | 130.1 | 7.07 (d, 8.3) |
| 11 | 115.6 | 6.71 (d, 8.3) | 115.6 | 6.73 (d, 8.3) |
| 12 | 156.6 | | 156.6 | |
| 13 | 115.6 | 6.71 (d, 8.3) | 115.6 | 6.73 (d, 8.3) |
| 14 | 130.2 | 7.04 (d, 8.3) | 130.1 | 7.07 (d, 8.3) |
| 15 | 174.6 | | 174.6 | |
| 16 | | 8.76 (d, 8.7) | | 8.97 (d, 8.4) |
| 17 | 54.4 | 4.81 (m) | 55.0 | 4.62 (m) |
| 18 | 34.9 | 3.17 (dd, 14.2, 3.2) | 35.0 | 3.24 (dd, 14.0, 3.2) |
| | | 2.65 (m) | | 2.64 (overlap) |
| 19 | 128.4 | | 128.5 | |
| 20 | 130.3 | 7.02 (d, 8.3) | 130.3 | 7.06 (d, 8.4) |
| 21 | 115.1 | 6.58 (d, 8.3) | 115.2 | 6.60 (d, 8.4) |
| 22 | 156.1 | | 156.2 | |
| 23 | 115.1 | 6.58 (d, 8.3) | 115.2 | 6.60 (d, 8.4) |
| 24 | 130.3 | 7.02 (d, 8.3) | 130.3 | 7.06 (d, 8.4) |
| 25 | 171.8 | | 171.7 | |
| 26 | | 7.40 (d, 8.1) | | 7.36 (d, 7.9) |
| 27 | 51.9 | 4.17 (ddd, 12.0, 8.1, 4.2) | 51.9 | 4.17 (ddd, 11.9, 8.0, 4.2) |
| 28 | 39.2 | 1.82 (m) | 39.3 | 1.81 (m) |
| | | 1.40 (m) | | 1.42 (m) |
| 29 | 24.6 | 1.74 (m) | 24.6 | 1.78 (m) |
| 30 | 21.1 | 0.83 (d, 6.5) | 21.1 | 0.84 (d, 6.4) |
| 31 | 23.4 | 0.88 (d, 6.6) | 23.8 | 0.88 (d, 5.2) |
| 32 | 171.7 | | 171.7 | |
| 33 | 72.0 | 5.11 (qd, 6.2, 1.8) | 72.1 | 5.12 (qd, 6.1, 1.7) |
| 34 | 16.0 | 1.02 (d, 6.2) | 16.0 | 1.04 (d, 6.1) |
| 35 | 56.2 | 4.46 (dd, 10.2, 1.8) | 56.4 | 4.44 (m) |
| 36 | | 7.97 (d, 10.2) | | 7.88 (d, 10.1) |
| 37 | 173.1 | | 173.9 | |
| 38 | 56.9 | 4.33 (dd, 10.5, 8.6) | 51.6 | 4.42 (m) |
| 39 | 35.2 | 1.86 (overlap) | 39.8 | 1.54 (m) |
| 40 | 24.7 | 1.44 (overlap) | 24.6 | 1.60 (m) |
| | | 1.20 (m) | | |
| 41 | 10.2 | 0.74 (t, 7.4) | 21.1 | 0.62 (d, 6.4) |
| 42 | 15.8 | 0.91 (d, 6.8) | 23.4 | 0.90 (d, 5.9) |
| 43 | | 8.00 (d, 8.6) | | 8.01 (d, 7.9) |
| 44 | 169.5 | | 169.5 | |
| 45 | 22.8 | 1.83 (s) | 22.8 | 1.81 (s) |

**Table S8**. $^{1}H$ (500 MHz) and $^{13}C$ (125 MHz) NMR spectroscopic data fof compounds **7** and **10** in DMSO-$d_6$ ($\delta$ in ppm and $J$ in Hz).

| no. | **7** | | **10** | |
|---|---|---|---|---|
| | $\delta_C$, type | $\delta_H$ (mult., $J$) | $\delta_C$, type | $\delta_H$ (mult., $J$) |
| 1 | 170.3 | | 169.0 | |
| 2 | | 7.03 (m) | | 6.93 (m) |
| 3 | 35.1 | 3.39 (m) | 36.7 | 3.50 (m) |
| | | 3.21 (m) | | 3.20 (m) |
| 4 | 34.4 | 2.37 (m) | 34.1 | 2.50 (m) |
| | | 2.19 (m) | | 2.26 (m) |
| 5 | 172.2 | | 173.0 | |
| 6 | | 8.20 (d, 6.6) | | 8.38 (d, 4.6) |
| 7 | 56.4 | 4.22 (m) | 57.2 | 4.28 (m) |
| 8 | 36.1 | 2.54 (m) | 35.5 | 2.65 (m) |
| | | | | 2.42 (dd, 14.2, 3.8) |
| 9 | 128.2 | | 127.8 | |
| 10 | 130.4 | 6.94 (d, 8.4) | 130.2 | 7.08 (d, 8.4) |
| 11 | 115.4 | 6.65 (d, 8.4) | 115.6 | 6.72 (d, 8.4) |
| 12 | 156.4 | | 156.6 | |
| 13 | 115.4 | 6.65 (d, 8.4) | 115.6 | 6.72 (d, 8.4) |
| 14 | 130.4 | 6.94 (d, 8.4) | 130.2 | 7.08 (d, 8.4) |
| 15 | 172.2 | | 174.8 | |
| 16 | | 8.29 (d, 8.9) | | 8.89 (d, 8.4) |
| 17 | 55.1 | 4.32 (m) | 55.0 | 4.49 (m) |
| 18 | 36.0 | 3.02 (dd, 13.9, 3.8) | 35.5 | 3.11 (dd, 14.0, 3.0) |
| | | 2.68 (dd, 13.9, 10.4) | | 2.65 (m) |
| 19 | 128.6 | | 128.5 | |
| 20 | 130.4 | 6.97 (d, 8.5) | 130.4 | 7.04 (d, 8.4) |
| 21 | 115.4 | 6.65 (d, 8.5) | 115.3 | 6.62 (d, 8.4) |
| 22 | 156.3 | | 156.3 | |
| 23 | 115.4 | 6.65 (d, 8.5) | 115.3 | 6.62 (d, 8.4) |
| 24 | 130.4 | 6.97 (d, 8.5) | 130.4 | 7.04 (d, 8.4) |
| 25 | 171.8 | | 171.6 | |
| 26 | | 7.57 (d, 7.9) | | 7.40 (d, 7.9) |
| 27 | 51.5 | 4.32 (overlap) | 51.9 | 4.13 (m) |
| 28 | 39.6 | 1.67 (m) | 39.4 | 1.78 (m) |
| | | 1.51 (m) | | 1.41 (m) |
| 29 | 24.5 | 1.67 (overlap) | 24.5 | 1.73 (m) |
| 30 | 22.0 | 0.86 (d, 6.3) | 21.2 | 0.84 (d, 7.0) |
| 31 | 22.3 | 0.90 (d, 6.3) | 23.3 | 0.88 (d, 6.5) |
| 32 | 171.5 | | 171.6 | |
| 33 | 71.4 | 5.21 (qd, 6.3, 3.5) | 71.5 | 5.20 (qd, 6.4, 2.1) |
| 34 | 16.9 | 1.06 (d, 6.3) | 16.1 | 1.04 (d, 6.4) |
| 35 | 54.9 | 4.70 (dd, 9.3, 3.5) | 56.2 | 4.50 (dd, 9.8, 2.1) |
| 36 | | 7.78 (d, 9.3) | | 7.69 (d, 9.8) |
| 37 | 173.2 | | 171.6 | |
| 38 | 35.5 | 2.19 (overlap) | 57.9 | 4.48 (dd, 8.6, 3.0) |
| 39 | 25.5 | 1.51 (overlap) | | 7.63 (d, 8.6) |
| 40 | 31.4 | 1.21 (m) | 172.9 | |
| 41 | 22.3 | 1.25 (m) | 35.1 | 2.08 (m) |
| 42 | 14.3 | 0.84 (t, 7.0) | 25.4 | 1.47 (m) |
| 43 | 169.8 | | 31.3 | 1.20 (m) |
| 44 | | 7.93 (d, 7.6) | 22.4 | 1.26 (m) |
| 45 | 59.6 | 3.93 (dd, 7.6, 3.8) | 14.4 | 0.84 (t, 7.0) |
| 46 | 65.8 | 4.09 (m) | 67.0 | 4.23 (m) |
| 47 | 20.7 | 1.00 (d, 6.4) | 20.1 | 1.07 (d, 6.3) |

**Table S9.** $^{1}H$ (500 MHz) and $^{13}C$ (125 MHz) NMR spectroscopic data for compound **26** in DMSO-$d_6$ ($\delta$ in ppm and $J$ in Hz).

| no. | **26** | |
|---|---|---|
| | $\delta_C$, type | $\delta_H$ (mult., $J$) |
| 1 | 157.7 | |
| 2 | | 8.05 (t, 5.0) |
| 3 | 40.8 | 3.04 (m) |
| 4 | 25.6 | 1.54 (m) |
| 5 | 28.1 | 1.74 (m) |
| 6 | 53.7 | 4.01 (m) |
| 7 | 171.0 | |
| 9 | 69.9 | 5.00 (m) |
| 10 | 15.0 | 1.10 (d, 6.5) |
| 11 | 54.5 | 4.48 (dd, 8.4, 4.4) |
| 12 | | 8.03 (d, 8.4) |
| 13 | 169.7 | |
| 14 | 22.8 | 1.90 (s) |
| 15 | 167.6 | |
| 16 | | 8.75 (d, 9.2) |
| 17 | 55.6 | 4.62 (m) |
| 18 | 29.8 | 3.16 (dd, 14.4, 8.0) |
| | | 2.99 (dd, 14.4, 6.7) |
| 19 | 109.9 | |
| 20 | 124.1 | 7.12 (d, 2.0) |
| 21 | | 10.87 (d, 2.0) |
| 22 | 136.5 | |
| 23 | 111.7 | 7.32 (d, 8.1) |
| 24 | 121.3 | 7.05 (m) |
| 25 | 118.8 | 6.96 (m) |
| 26 | 118.6 | 7.54 (d, 7.9) |
| 27 | 127.8 | |
| 28 | 172.0 | |
| 29 | | 8.56 (d, 4.7) |
| 30 | 61.0 | 4.19 (dd, 7.7, 4.7) |
| 31 | 66.0 | 3.91 (m) |
| 32 | 20.1 | 1.08 (d, 6.3) |
| 33 | 173.6 | |
| 34 | | 8.73 (d, 5.7) |
| 35 | 60.9 | 3.88 (dd, 5.7, 4.7) |
| 36 | 29.1 | 2.18 (m) |
| 37 | 17.8 | 0.90 (d, 6.9) |
| 38 | 19.4 | 0.95 (d, 7.0) |
| 39 | 171.4 | |
| 40 | | 6.83 (d, 8.0) |
| 41 | 56.7 | 4.19 (dd, 8.0, 4.9) |
| 42 | 35.5 | 1.87 (m) |
| 43 | 26.3 | 1.19 (m) |
| | | 1.14 (m) |
| 44 | 11.9 | 0.77 (t, 7.4) |
| 45 | 15.0 | 0.75 (d, 7.0) |
| 46 | 171.9 | |
| 47 | | 7.43 (d, 5.8) |

**Table S10.** $^{1}H$ (500 MHz) and $^{13}C$ (125 MHz) NMR data of compound **41** in DMSO-$d_6$ ($\delta$ in ppm). COSY (bold) and key HMBC (arrows) are shown.

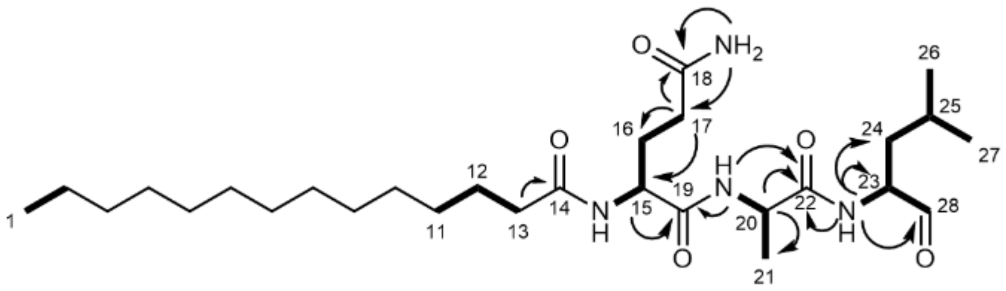


| Position | $\delta_C$, type$^a$ | $\delta_H$, mult. ($J$ in Hz) |
|---|---|---|
| 1 | - | 0.91 – 0.80, ov |
| 11 | - | 1.30 – 1.15, ov |
| 12 | 25.19, $CH_2$ | 1.55 – 1.38, ov |
| 13 | 35.15, $CH_2$ | 2.15 – 2.03, ov |
| 14 | 172.74, C | - |
| 15 | 52.33, CH | 4.21 - 4.14, m |
| 15NH | - | 8.05 – 7.92, m |
| 16 | 28.66, $CH_2$ | 1.91 – 1.66, m |
| 17 | 31.53, $CH_2$ | 2.15 – 2.03, m |
| 17NH | - | 7.24 (s) |
| 17NH | - | 6.74 (s) |
| 18 | 173.80, C | - |
| 19 | 171.21, C | - |
| 20 | 48.05, CH | 4.33 – 4.21, m |
| 20NH | - | 8.05 – 7.92, m |
| 21 | 18.19, $CH_3$ | 1.30 – 1.15, ov |
| 22 | 172.56, C | - |
| 23 | 56.54, CH | 4.14 – 4.05, m |
| 23NH | - | 8.23 – 8.17, m |
| 24 | 36.33, $CH_2$ | 1.55 – 1.38, ov |
| 25 | 24.00, CH | 2.15 – 2.03, ov |
| 26 | - | 0.91 – 0.80, ov |
| 27 | - | 0.91 – 0.80, ov |
| 28 | 201.05, CH | 9.39 – 9.35, m |

**Table S11**. Crystallographic data collection and refinement statistics of yCP:41.

| | ***yCPC14QAL*** |
|---|---|
| **Crystal parameters** | |
| Space group | $P2_1$ |
| Cell constants | a = 135.0 Å |
| | b = 300.9 Å |
| | c = 144.0 Å |
| | β = 112.8 ° |
| CPs / AU[a] | 1 |
| **Data collection** | |
| Beam line | X06SA, SLS |
| Wavelength (Å) | 1.0 |
| Resolution range (Å)[b] | 50–3.25 (3-35–3.25) |
| No. observations | 481076 |
| No. unique reflections[c] | 157953 |
| Completeness (%)[b] | 95.1 (93.7) |
| $R_{merge}$ (%)[b, d] | 10.5 (65.4) |
| I/□ (I)[b] | 11.1 (2.4) |
| **Refinement (REFMAC5)** | |
| Resolution range (Å) | 30-3.25 |
| No. refl. working set | 149904 |
| No. refl. test set | 7890 |
| No. non hydrogen | 49565 |
| No. of ligand atoms | 148 |
| Solvent ($H_2O$, ions, MES) | 95 |
| $R_{work}/R_{free}$ (%)[e] | 17.5 / 21.2 |
| r.m.s.d. bond (Å) / angle (°)[f] | 0.003 / 1.2 |
| Average B-factor ($Å^2$) | 91.3 |
| Ramachandran Plot (%)[g] | 97.6 / 2.2 / 0.2 |
| PDB accession code | 8BW1 |

[a] Asymmetric unit

[b] The values in parentheses for resolution range, completeness, $R_{merge}$ and I/σ (I) correspond to the highest resolution shell

[c] Data reduction was carried out from a single crystal. Friedel pairs were treated as identical reflections

[d] $R_{merge}(I) = \Sigma_{hkl}\Sigma_j \mid I(hkl)_j - \langle I(hkl)\rangle \mid / \Sigma_{hkl} \Sigma_j I(hkl)_j$, where $I(hkl)_j$ is the $j^{th}$ measurement of the intensity of reflection hkl and $\langle I(hkl)\rangle$ is the average intensity

[e] $R = \Sigma_{hkl} \mid |F_{obs}| - |F_{calc}| \mid / \Sigma_{hkl} |F_{obs}|$, where $R_{free}$ is calculated without a sigma cut off for a randomly chosen 5% of reflections, which were not used for structure refinement, and $R_{work}$ is calculated for the remaining reflections

[f] Deviations from ideal bond lengths/angles

[g] Percentage of residues in favored / allowed / outlier region

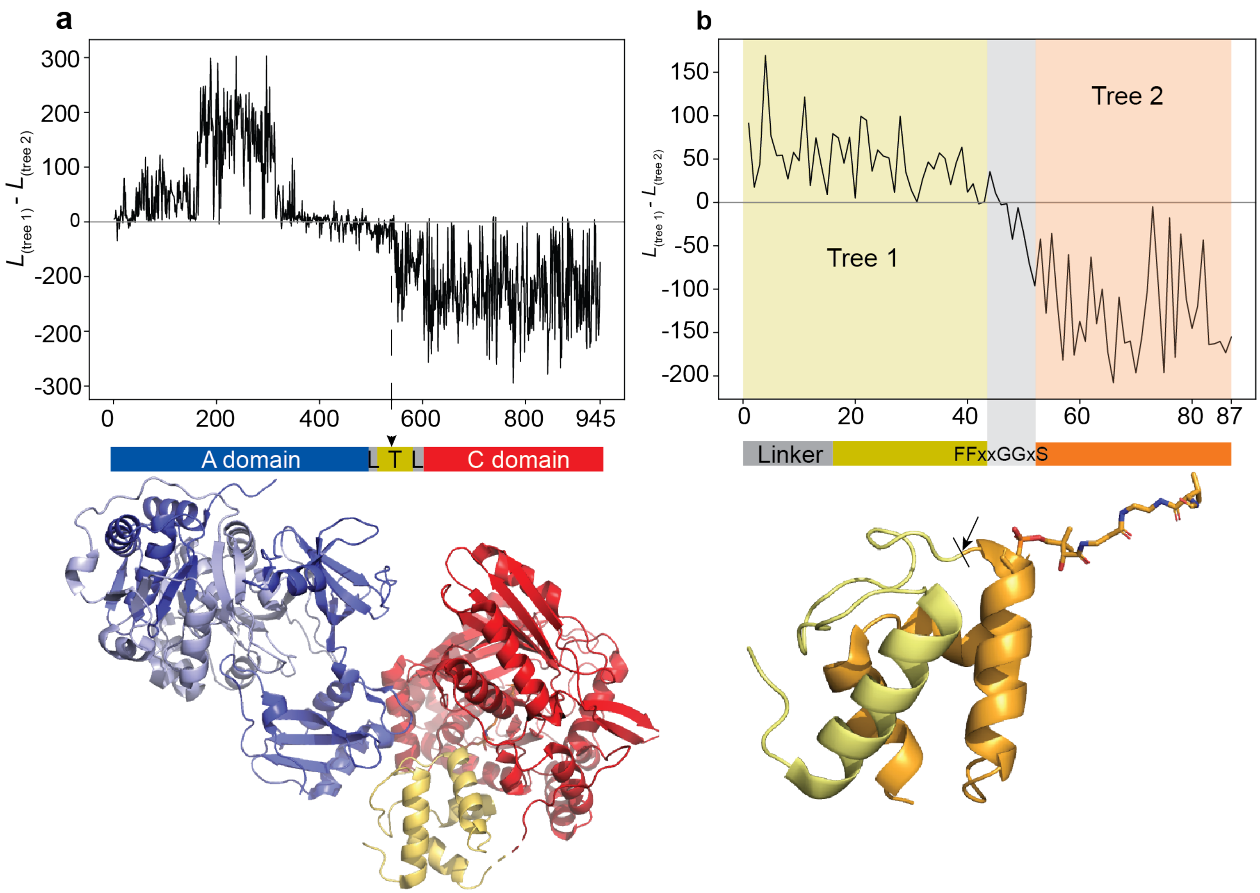


**Fig. S1. Evolutionary analysis of ATC tridomains and T domains of representative NRPS.** (**a**) Likelihood difference plot of two phylogenetic trees of ATC tridomains (also called XUs) that together best describe the alignment using a phylogenetic hidden Markov model. Positive numbers indicate that sites are better describe by tree 1, negative numbers indicate sites that are better described by tree two. Protein structure of XU is shown below. A domain is colored in blue, T-domain in yellow and C domain in red. (**b**) Likelihood difference plot as in **a**, but for an alignment of T domain plus A-T linker. Partitions detected by the hidden Markov model are indicated in different colors according to tree number. Recombination breakpoint is annotated in grey and lies around two conserved glycines. Protein structure of A-T-Linker and T domain is shown below. The first part of the T domain is colored in yellow and the second part in orange. An arrow points to the fusion site used for engineering.

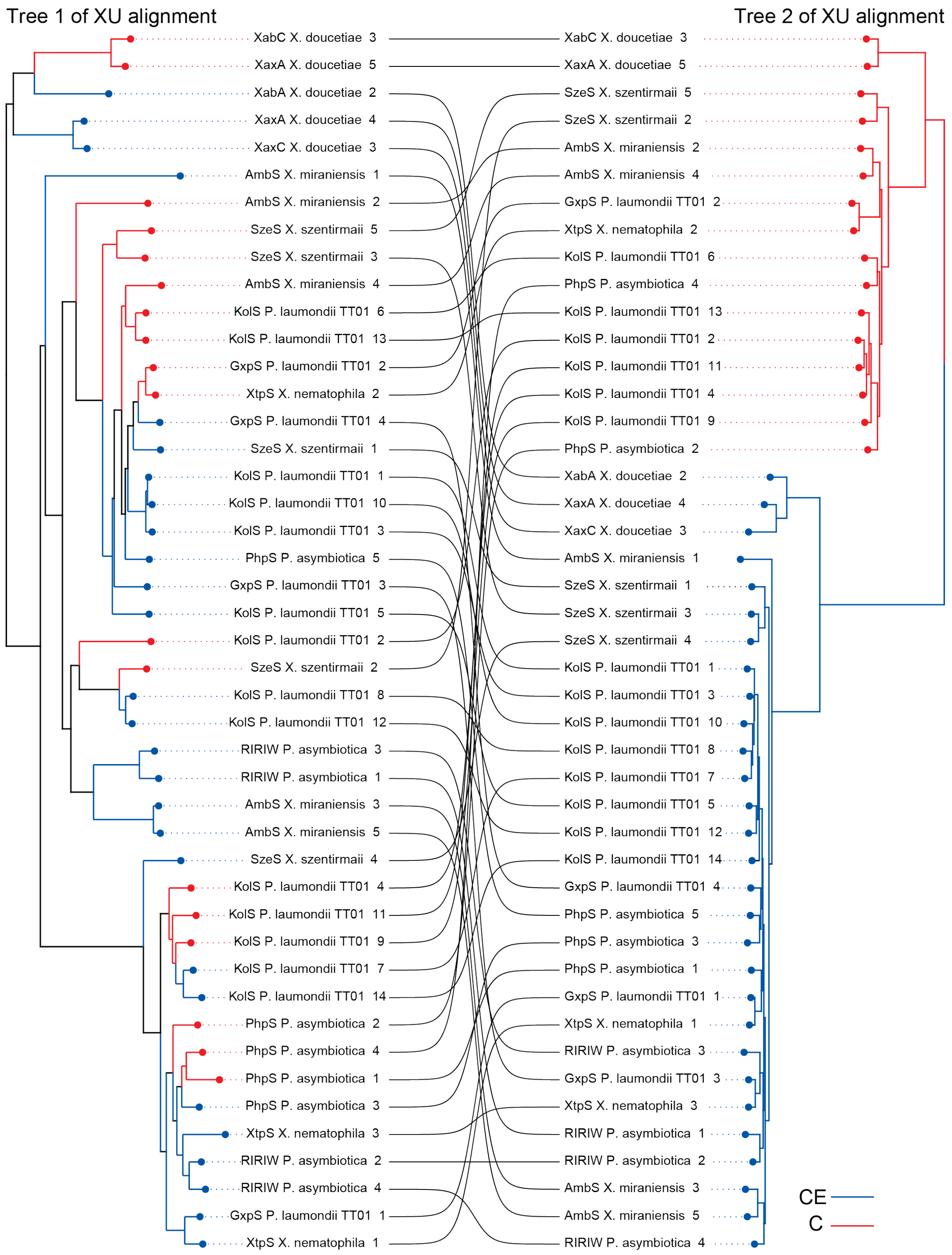


**Fig S2.** Comparison of Tree 1 and Tree 2 from the XU alignment. Taxon names indicate abbreviation of NRPSs, followed by bacterial species and then numbers of the XU within that NRPS. Lines connect the same NRPS and XU between the two trees. Red branches label XUs that contain $^{L}C_{L}$ domains and blue branches label XUs with CE (dual C) domains.

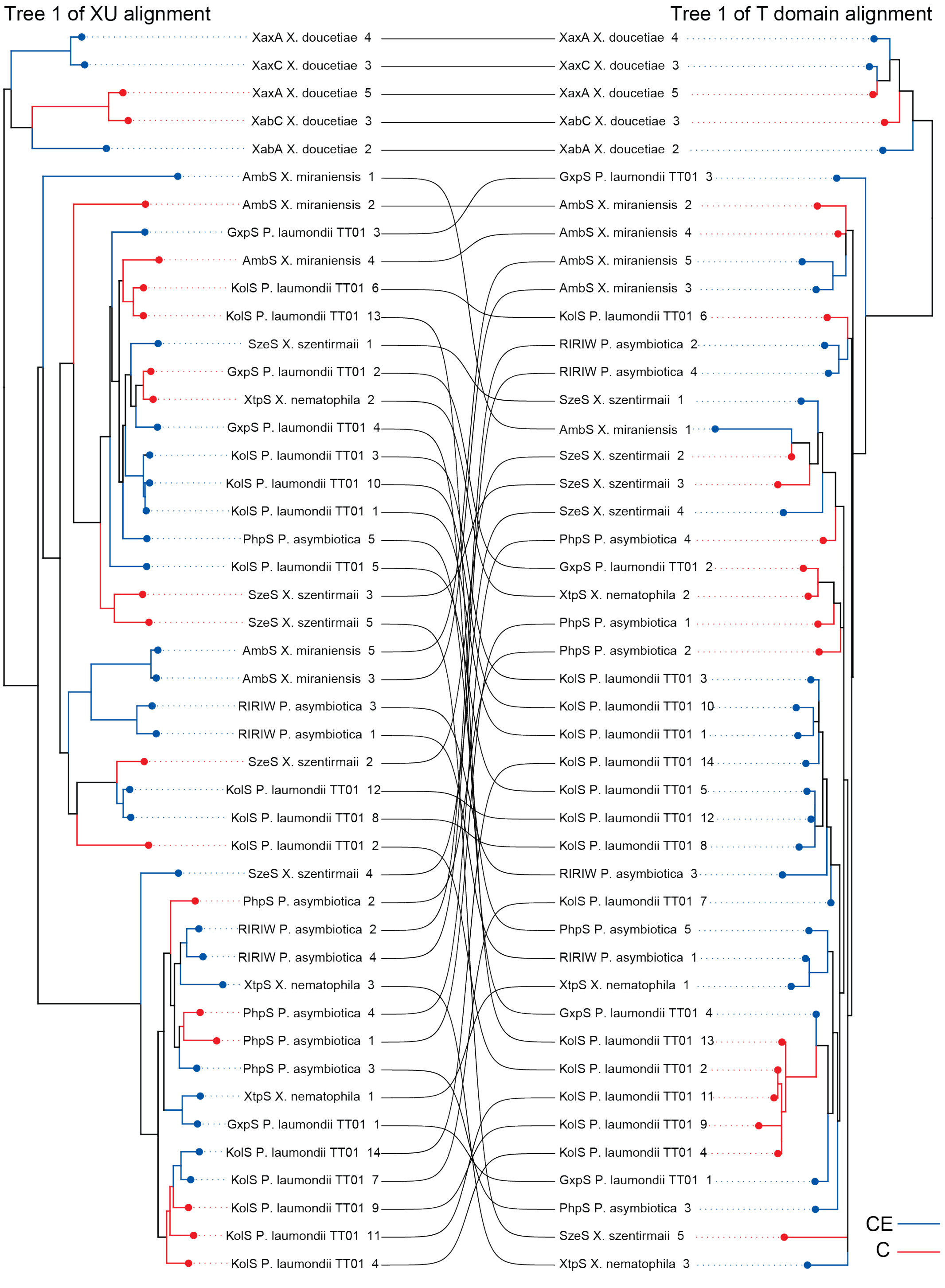


**Fig. S3.** Comparison of Tree 1 from XU domain alignment and Tree 1 from T domain alignment. Taxon names indicate abbreviation of NRPS, followed by bacterial species and then numbers of the XU within that NRPS. Lines connect the same NRPS and XU between the two trees. Red branches label $^{L}C_{L}$ domains and blue branches label CE (dual C) domains.

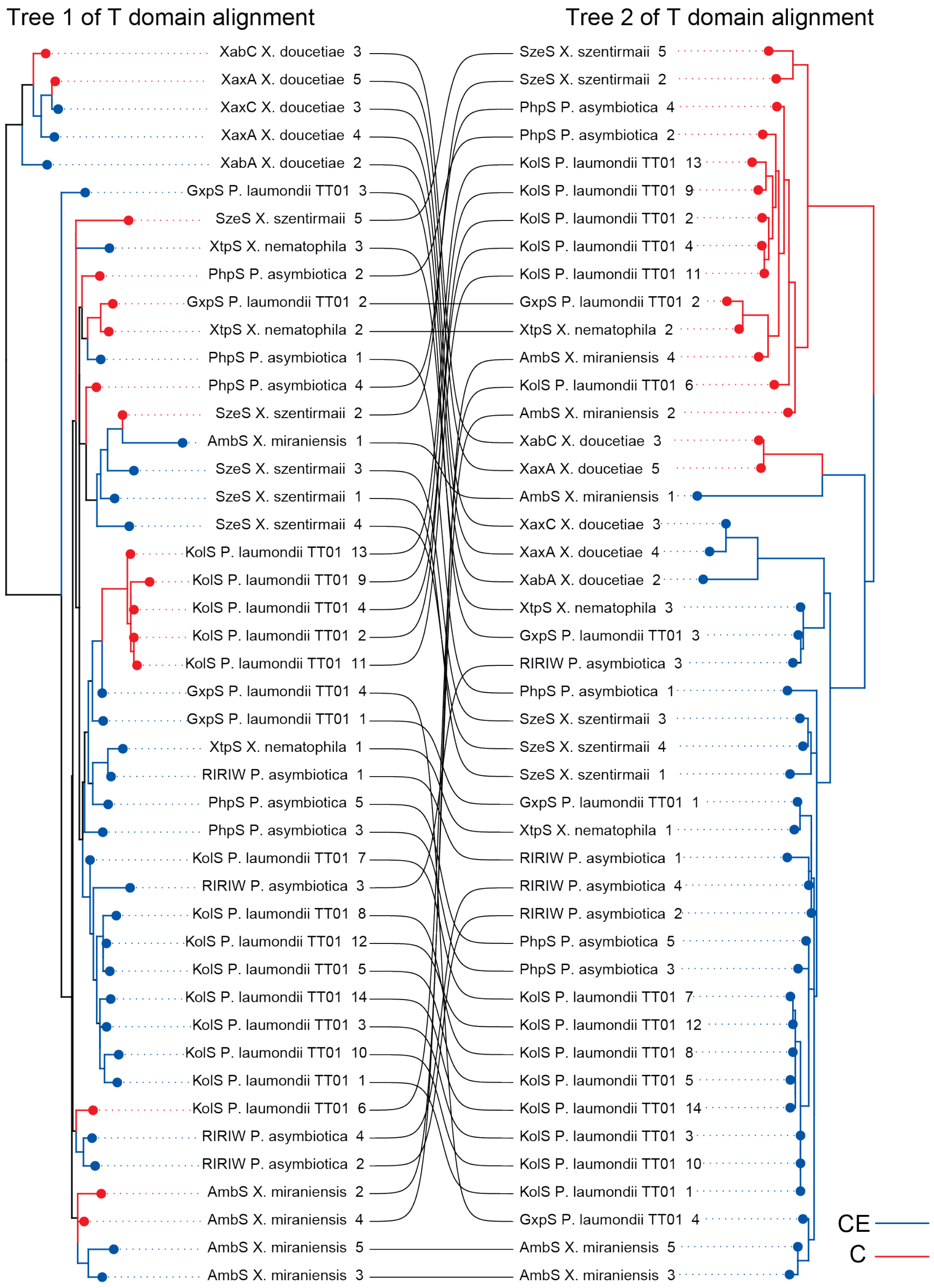


**Fig S4.** Comparison of Tree 1 from T domain alignment and Tree 2 from T domain alignment. Taxon names indicate abbreviation of NRPS, followed by bacterial species and then numbers of the XU within that NRPS. Lines connect the same NRPS and XU between the two trees. Red branches label $^{L}C_{L}$ domains and blue branches label CE (dual C) domains.

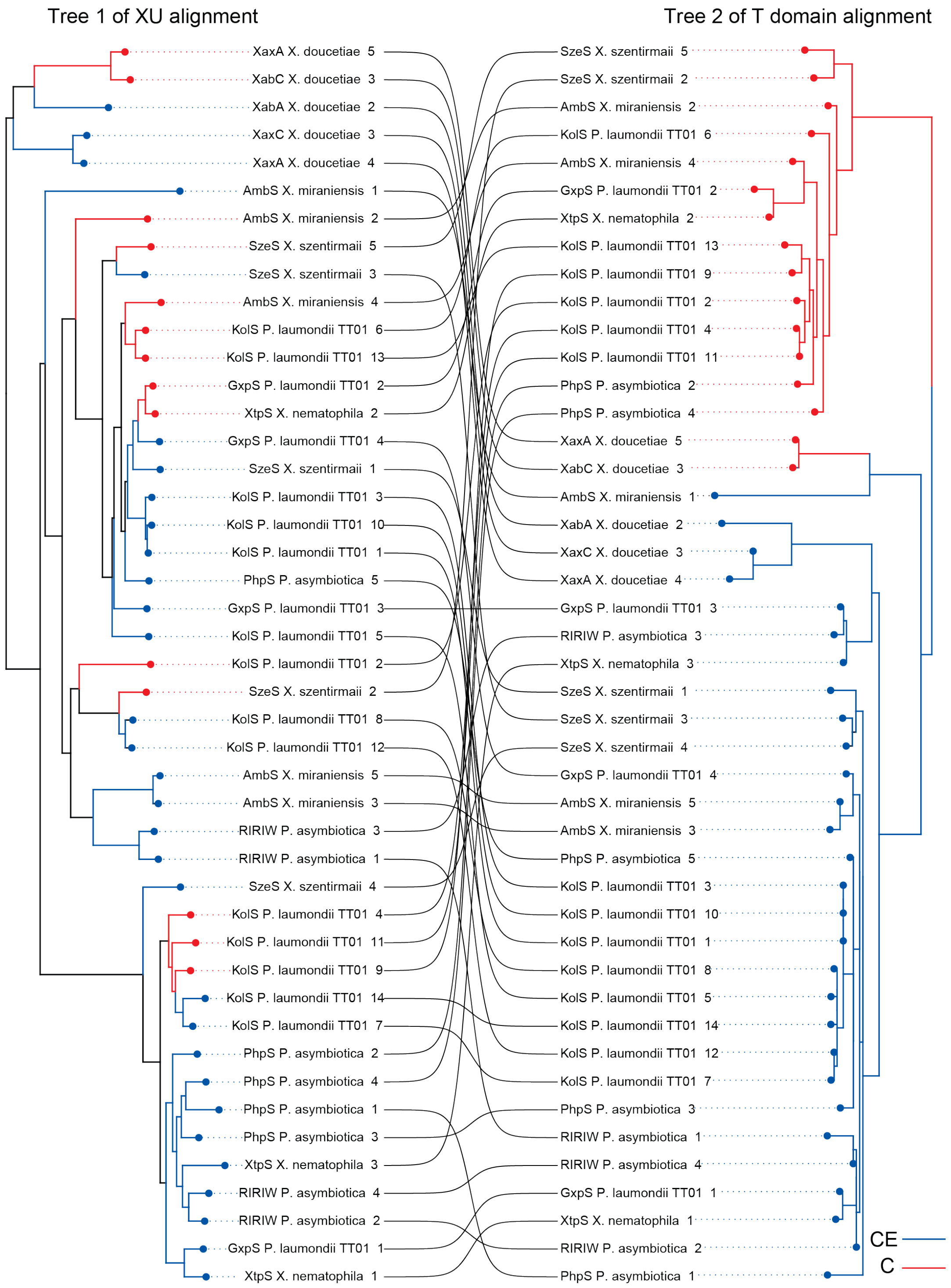


**Fig. S5.** Comparison of Tree 1 from XU domain alignment and Tree 2 from T domain alignment. Taxon names indicate abbreviation of NRPS, followed by bacterial species and then numbers of the XU within that NRPS. Lines connect the same NRPS and XU between the two trees. Red branches label $^{L}C_{L}$ domains and blue branches label CE (dual C) domains.

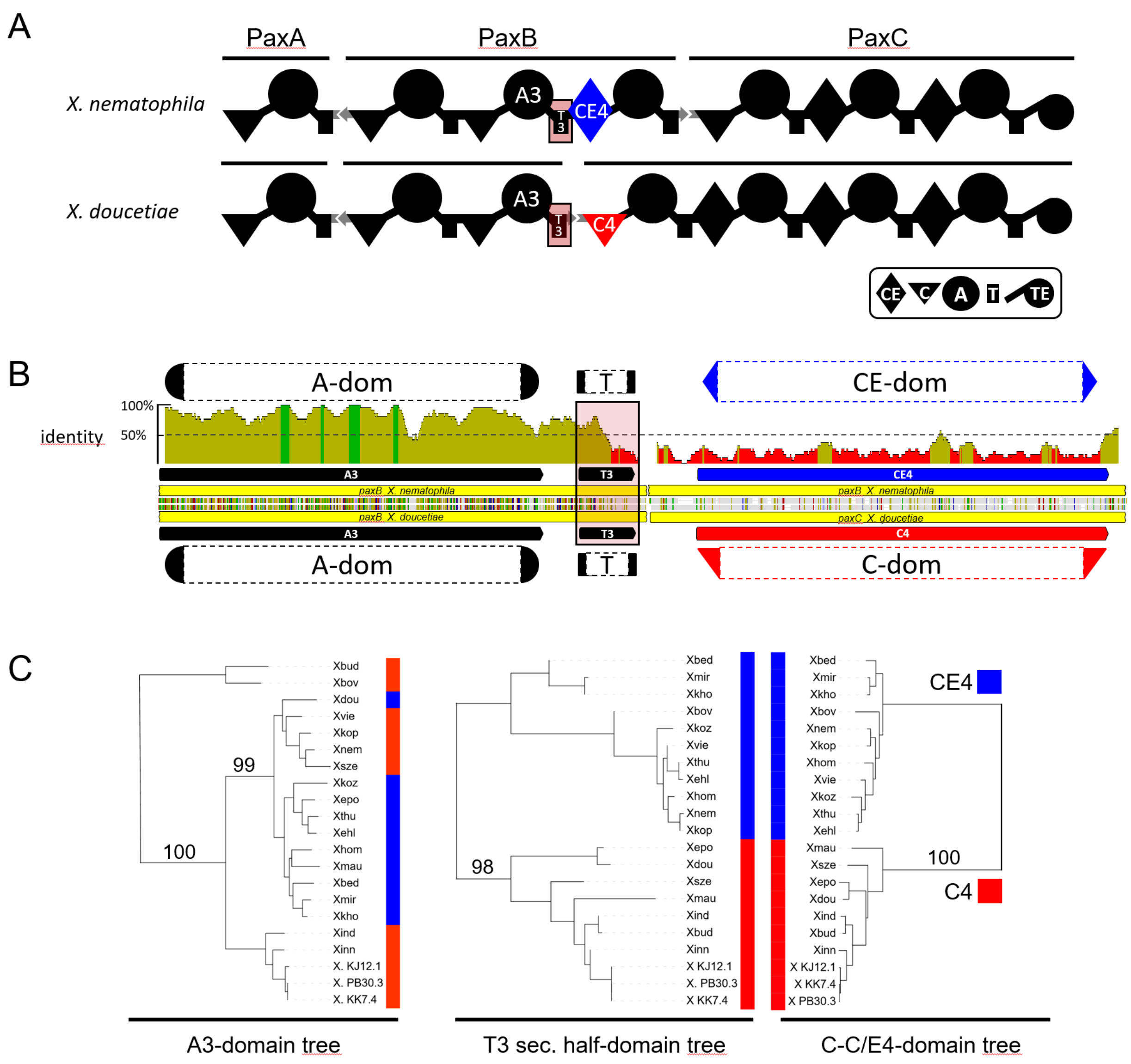


**Figure S6.** Phylogenetic analysis of T-domains in relation to preceding A-domain and following C/CE-domain. In (**A**) a schematic representation of the PAX producing NRPS[25,26] with the T3-domain under scrutiny highlighted (reddish square). The following CE4-domain is shown in blue and the C4-domain in red. In (**B**) a dual alignment of the A3-T3-CE4/C4 domains from *X. nematophila compared* to *X. doucetiae* can be seen. The amino acid alignment in the middle is shown with agreements in colour, genes as yellow bars and the domains indicated in the colour used in (**A**). The mean pairwise identity over all pairs in the column are calculated for a sliding window size 20 amino acids (green 100% identity, greenish-brown at least 30% under 100%, red below 30%). The drop of pairwise identity from high value between the A-domain region to the low identity between C- and CE-domains occurs in the middle of the T-domain. In (**C**) a phylogenetic tree of A3-, T3- second half (corresponds to T-fusion point IV in figure 2) and C4/CE4-domains is presented. The phylogenetic tree was calculated for the A3-domain, T3-domain second half and the following C/CE-domains separately. To this end multiple alignments of the protein sequences were generated using Clustal Omega 1.2.2.[27] with the refinement iterations number set at 10 while evaluating the full distance matrix for the initial guide tree as well as for the refinement iteration guide tree. Only bootstrap values at critical junctions are indicated. The colours blue (CE) and red (C) refer to the condensation domains of the A3-T3-C4/CE4 unit. Abbreviations of the indicated PAX NRPS organisms: Xbud, *X. budapestensis*; Xbed, *X. beddingii*; Xbov, *X. bovienii*; Xdou, *X. doucetiae*; Xehl, *X. ehlersii*; X eap, *X. eapokensis*; Xhom, *X. hominickii*; Xind, *X. indica*; Xkho, *X. khoisanae*; Xkop, *X. koppenhoeferi*; Xkoz, *X. kozodoii*; Xmau, *X. mauleonii*; Xmir, *X. miraniensis*; Xnem, *X. nematophila*; Xsze, *X. szentirmaii*; Xthu, *X. thuongxuanensis* str. 30TX1, Xvie, *X. vietnamensis, X. sp.* KJ12.1, X KK7.4, *X.* sp. KK7.4, *X.* sp. PB30.3, X PB30.3). PaxABC sequences were identified using the PaxABC peptide sequences of *X. nematophila* and *X. doucetiae* as query. Domain annotation was implemented by use of AntiSMASH 6.0[28].

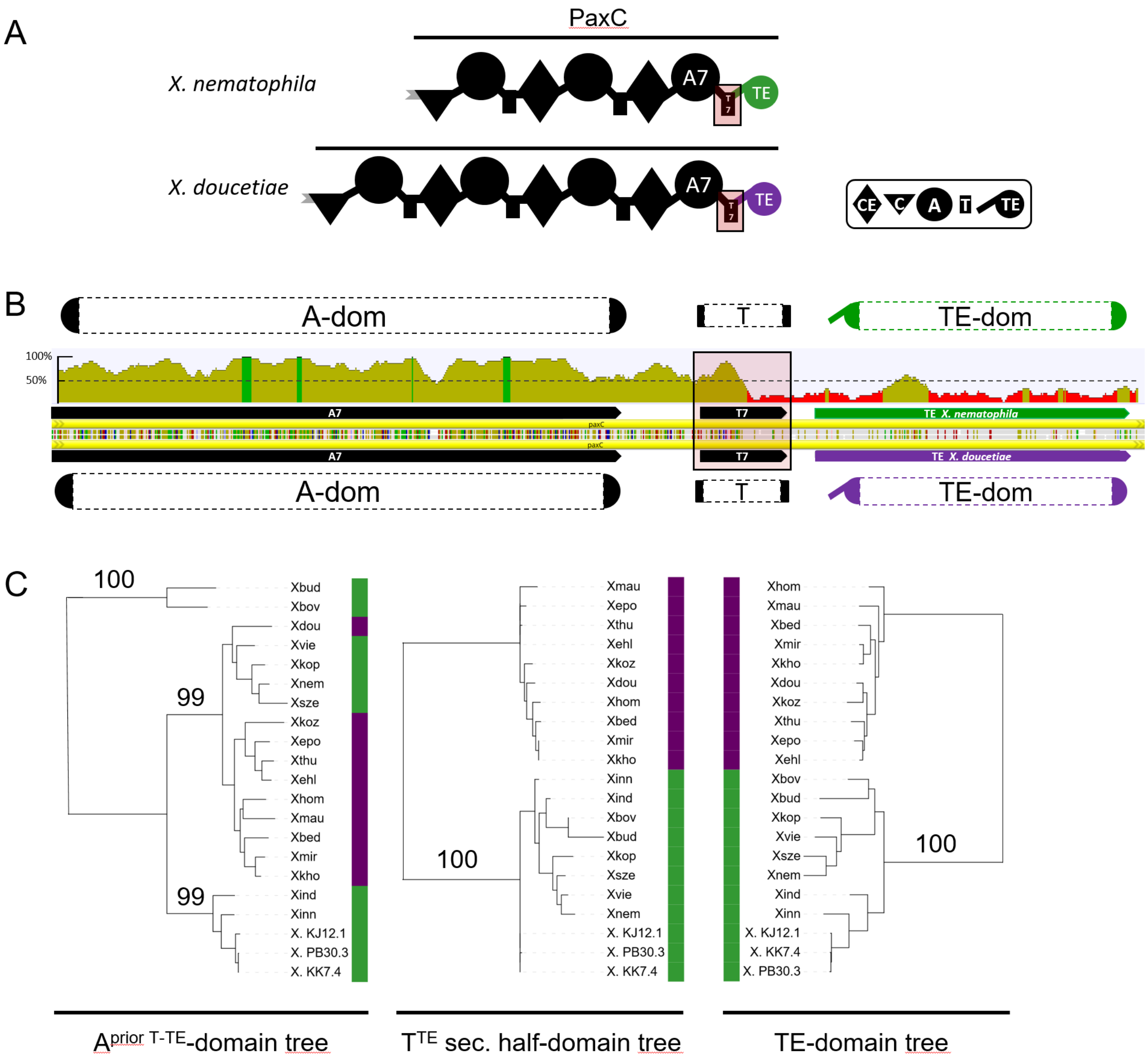


**Figure S7**. Phylogenetic analysis of T-domains in relation to preceding A-domain and the following TE-domain of the PAX-NRPS. The PAX biosynthesis in *Xenorhabdus* contains one of two types TE-domains being equally distributed in the *in silico* accessible biosynthesis. In (**A**) the final NRPS multienzymes are depicted with the *X. nematophila* TE-type in green and the *X. doucetiae* TE-type in purple. (**B**) A dual alignment of the A7-T7-TE unit from *X. nematophila* and *X. doucetiae* visualises the low identity between the two TE-types and that the drop of the sequence identity occurs in the middle of the T-domain. The phylogenetic tree in (**C**) was derived as described in Figure S6. The colour bars in all three phylogenetic trees refer to the TE in the A-T-TE unit. The *Xenorhabdus* species abbreviations are as in Figure S6.

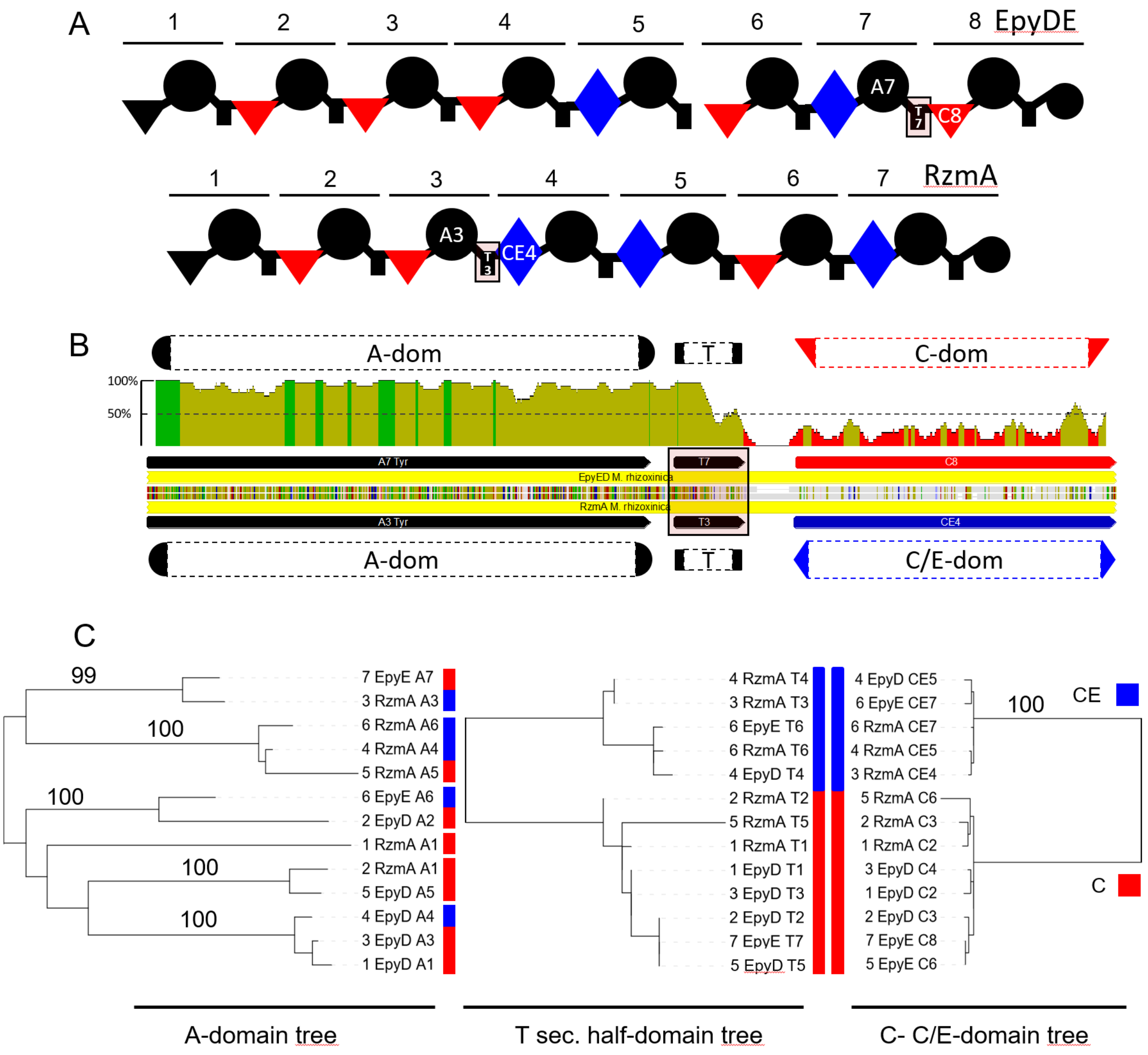


**Figure S8.** Phylogenetic analysis of T-domains in relation to preceding A-domain and following C/CE-domain of RzmA and EpyDE. (**A**) Schematic representation of the endopyrrole A producing NRPS EpyDE[29] and the rhizomide A producing NRPS RzmA[30] from *Mycetohabitans rhizoxinica* (DSM 19002). In (**B**) the EpyDE A7-T7-C8 unit and the RzmA A3-T3-CE4 unit are shown in a dual alignment. The phylogenetic trees of the A-domains, the T-domain second half and the C/CE-domains of RzmA and EpyDE were generated separately as described in Figure S6 using the same colour code (**C**).

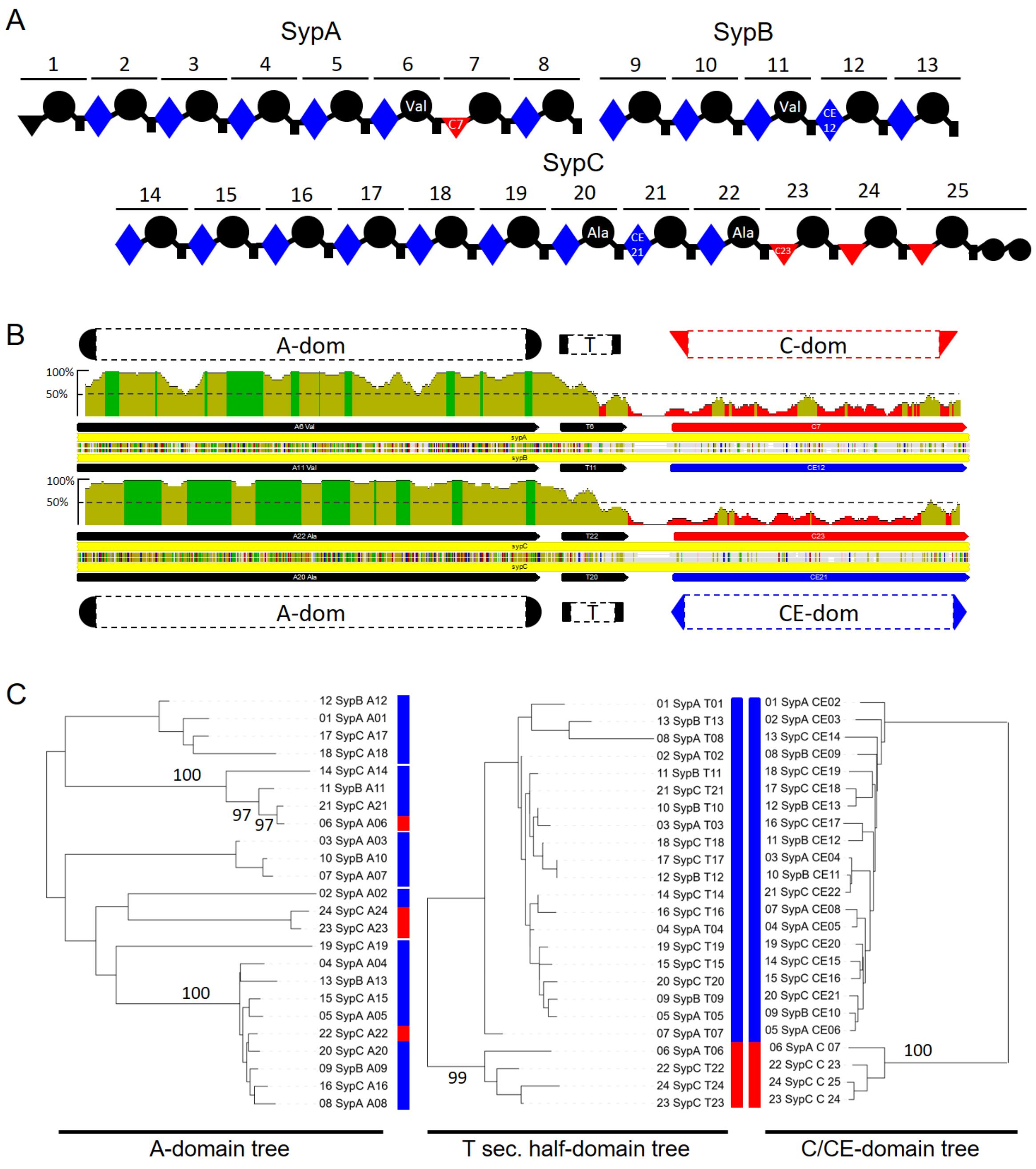


**Figure S9.** Phylogenetic analysis of T-domains in relation to preceding A-domain and following C/CE-domain of the syringopeptin SP-25a NRPS synthesis (SypABC; ALU60730.1, ALU60731.1, ALU60732.1) of *Pseudomonas syringae pv. lapsa* (DSM 50274) (**A**). The indicated A-domain substrate specificity was derived from published SP-25a[31] in conjunction with AntiSMASH 6.0 predictions[28]. In (**B**) two dual alignments of the SypA A7-T7-C8 to the SypB A11-T11-CE12 (top) and the SypC A20-T20-CE21 unit to the A22-T22-C23 (bottom) are shown. The phylogenetic trees of the A-domains, the T-domain second half and the C/CE-domains of SypABC were generated separately as described in Figure S6 using the same colour code (**C**).

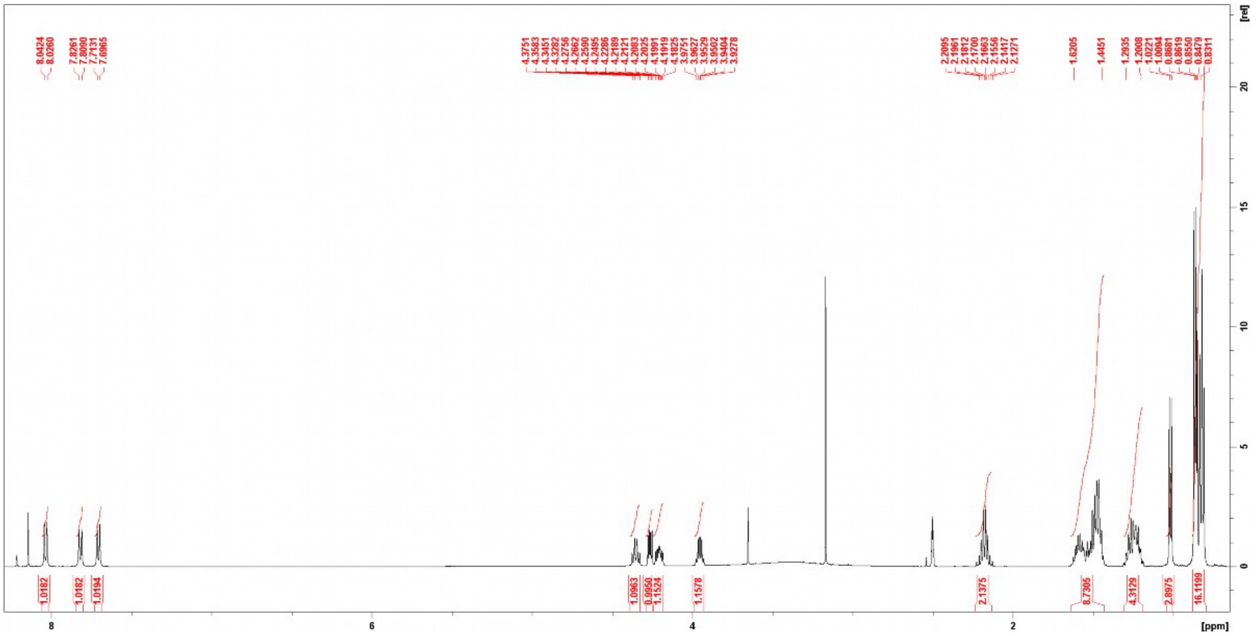

**Figure S10.** $^{1}H$ NMR (500 MHz, DMSO-$d_6$) spectrum compound **1**.

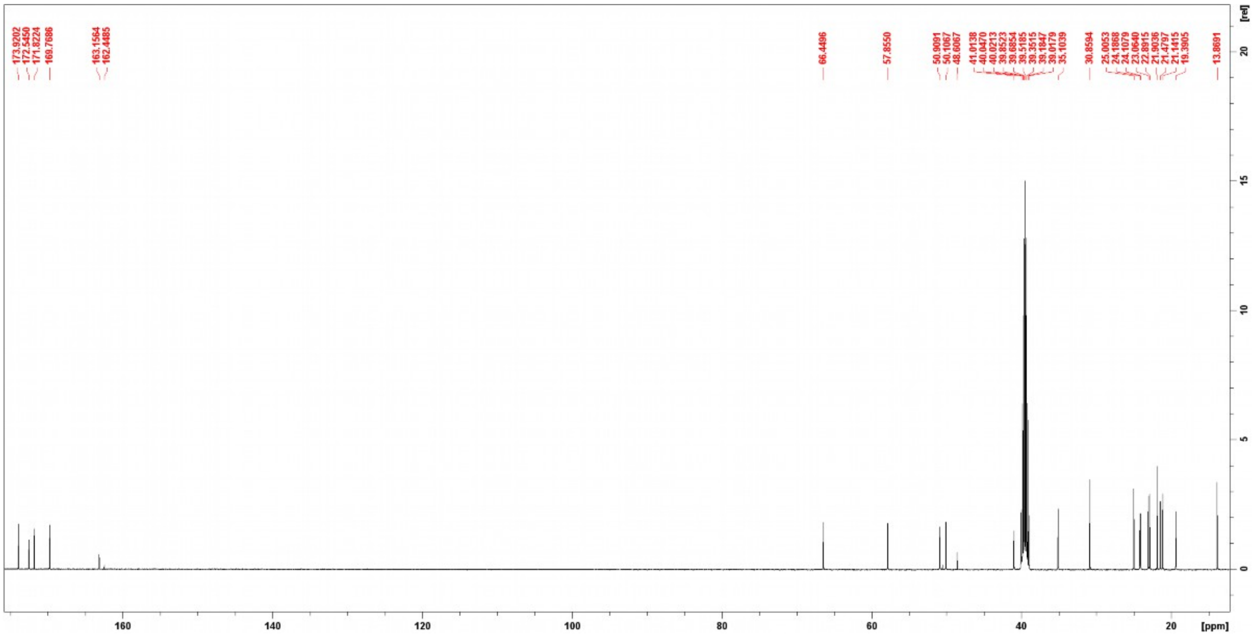

**Figure S11.** $^{13}C$ NMR (125 MHz, DMSO-$d_6$) spectrum of compound **1**.

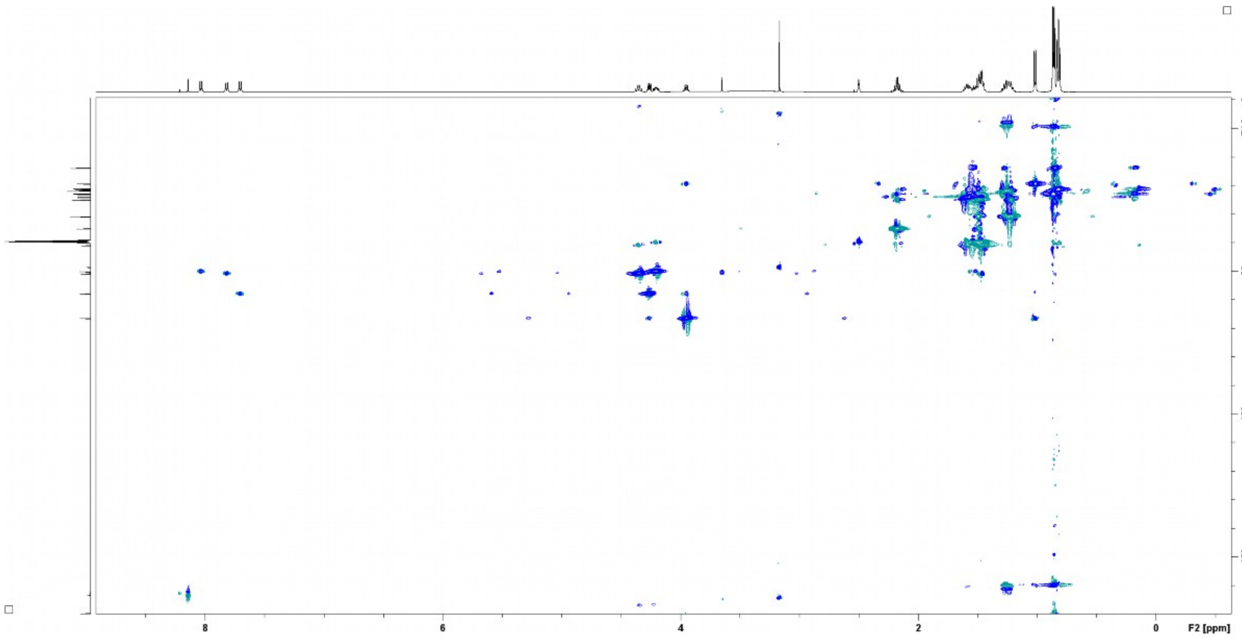

**Figure S12.** HSQC (DMSO-d$_6$) spectrum compound **1**.

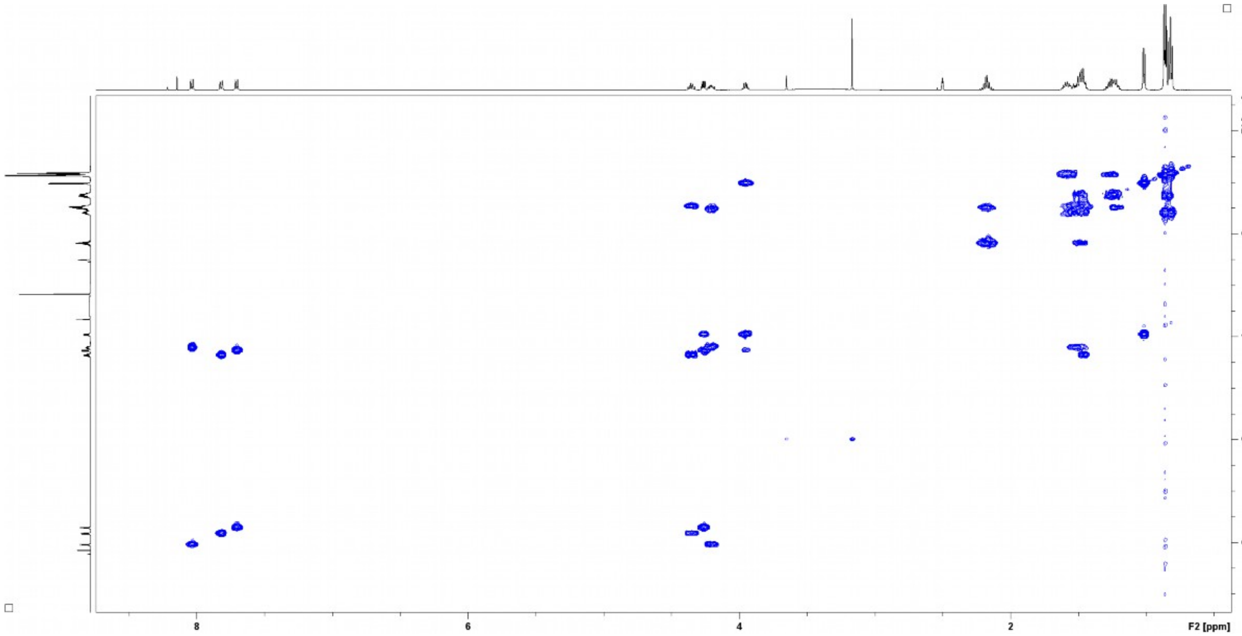

**Figure S13.** $^1$H-$^1$H COSY (DMSO-d$_6$) spectrum of compound **1**.

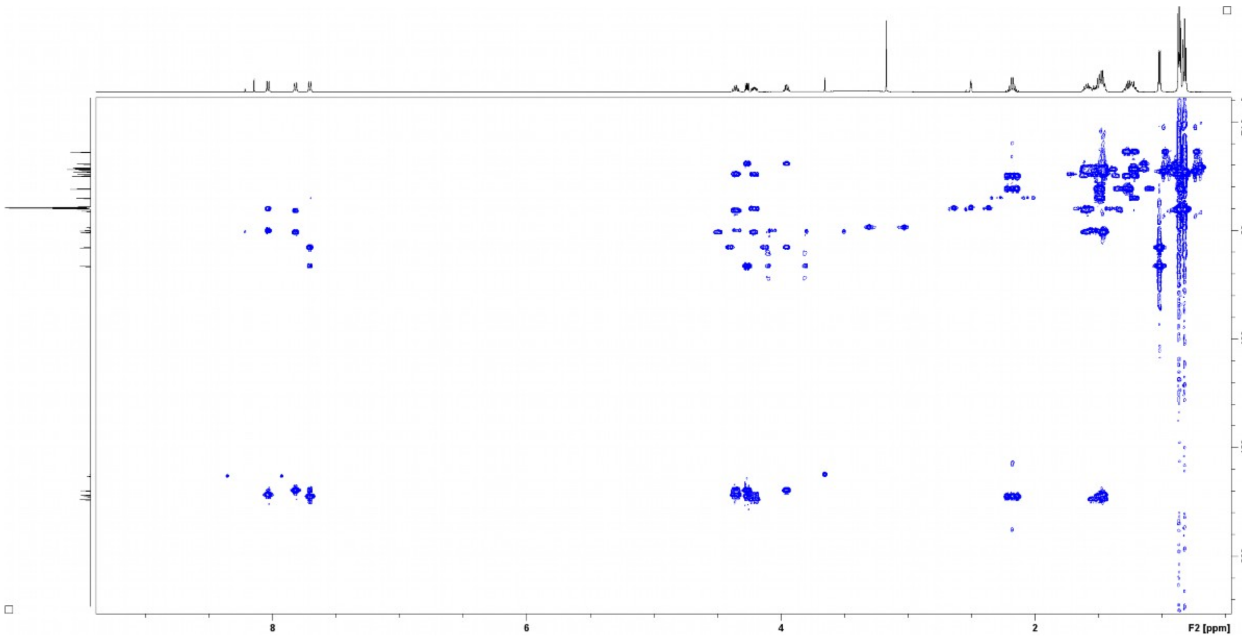


**Figure S14.** HMBC (DMSO-$d_6$) spectrum of compound **1**.

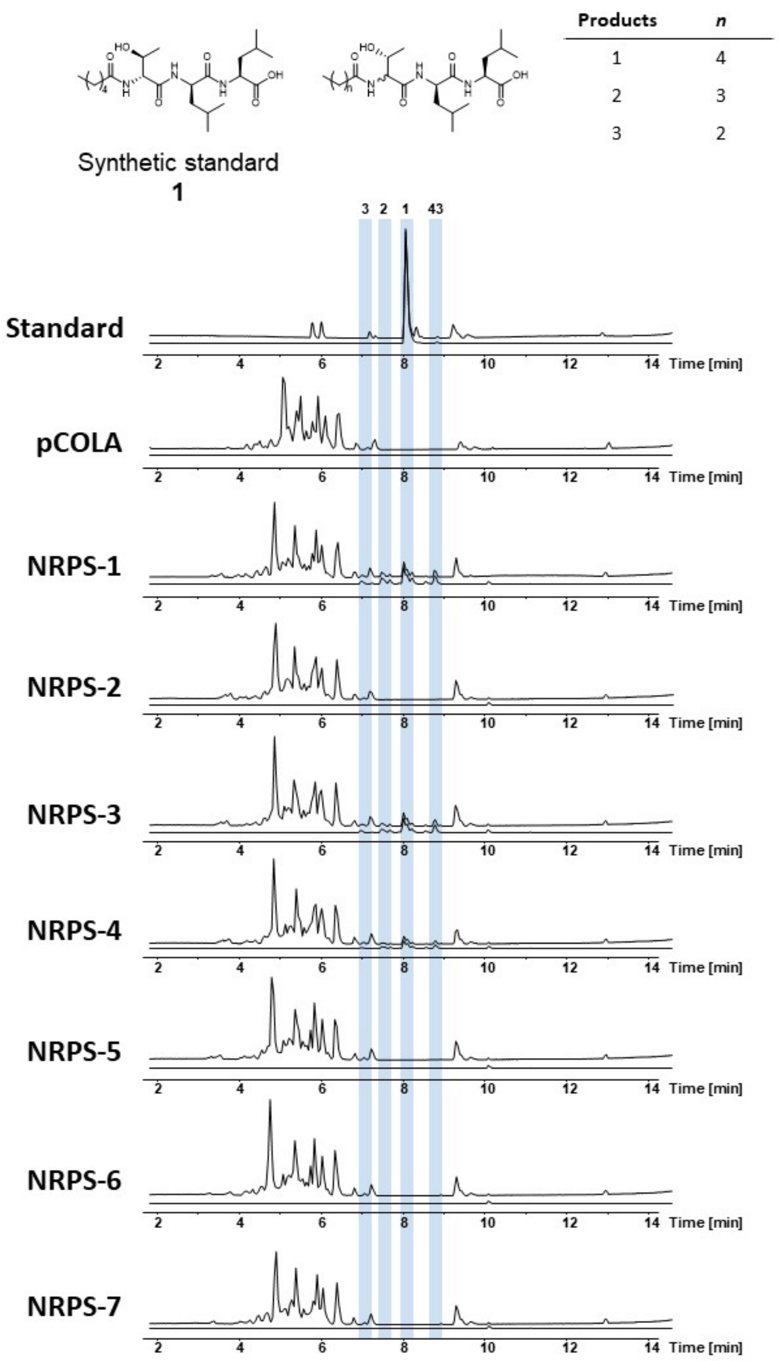


**Figure S15.** HPLC/MS data refers to Figure 2 (NRPS-1 to -7) of compound **1**, **2**, **3** and **43** produced in *E. coli* DH10B::*mtaA*. Base Peak Chromatogram (BPC, top) and Extracted Ion Chromatogram (EIC, below) of **1** (*m/z* $[M+H]^+$ = 444.30), **2** (*m/z* $[M+H]^+$ = 430.29), **3** (*m/z* $[M+H]^+$ = 416.27) and **43** (*m/z* $[M+H]^+$ = 458.32). Chromatograms were compared to an empty vector control and a synthetic standard of compound **1** (*m/z* $[M+H]^+$ = 444.30).

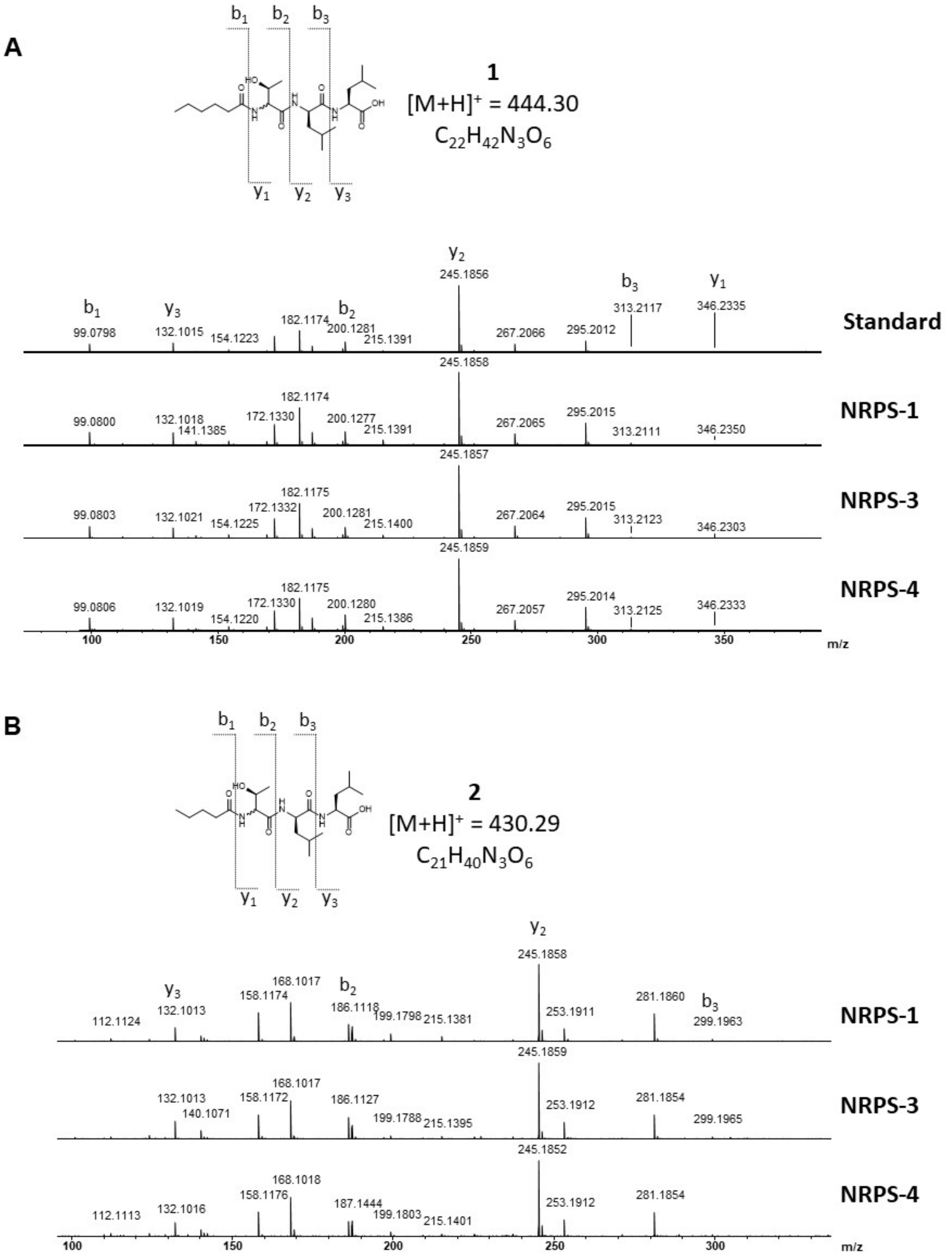


**Figure S16.** HPLC/MS data refers to Figure 2 (NRPS-1, -3 and -4) of compound **1** (**A**) and **2** (**B**) produced in *E. coli* DH10B::*mtaA*. Comparison of $MS^2$ spectra. Compound **1** fragmentation was compared to a synthetic **1**.

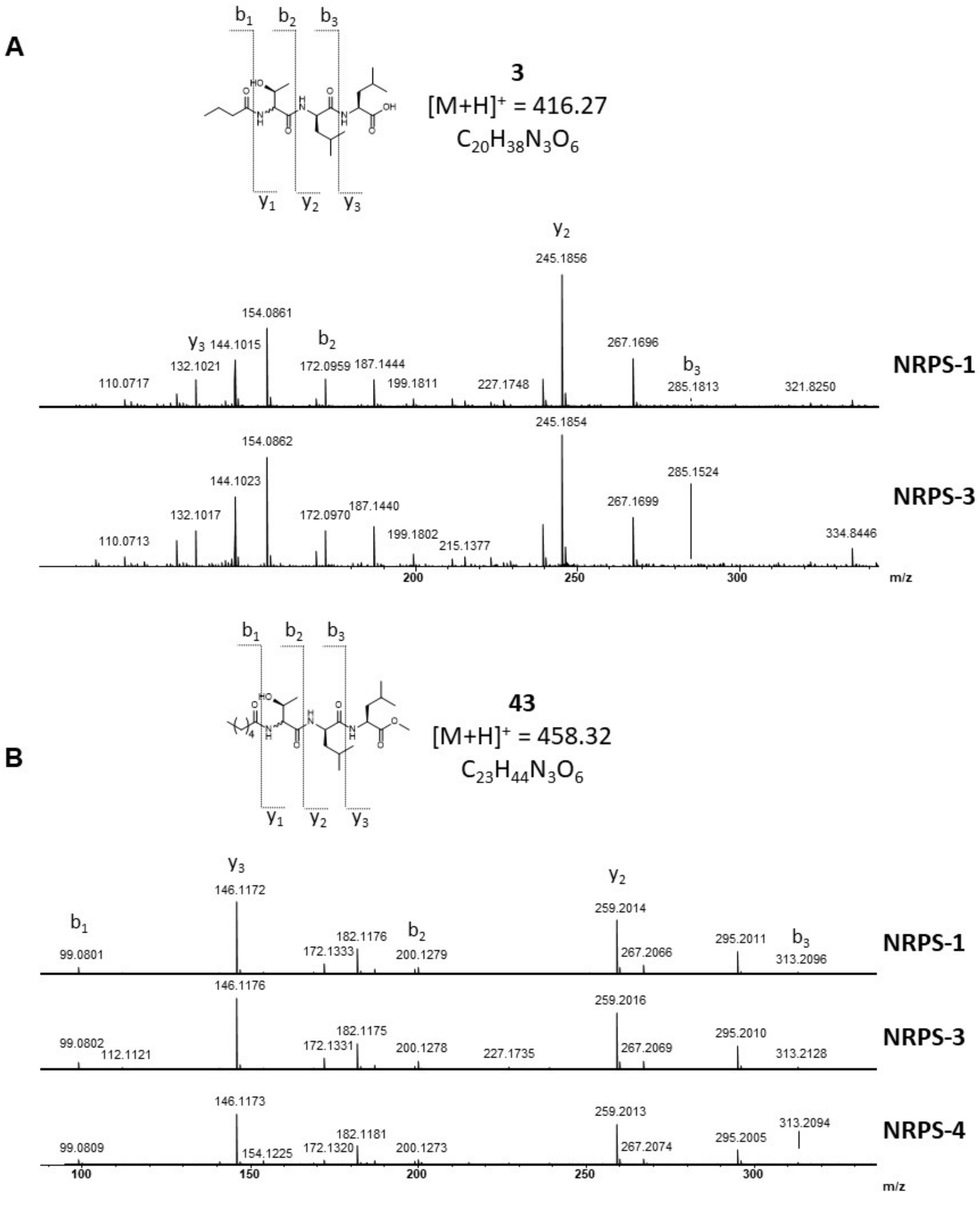


**Figure S17.** HPLC/MS data refers to Figure 2 (NRPS-1, -3 and -4) of compound **3** (**A**) and **43** (**B**) produced in *E. coli* DH10B::*mtaA*. Comparison of $MS^2$ spectra.

A

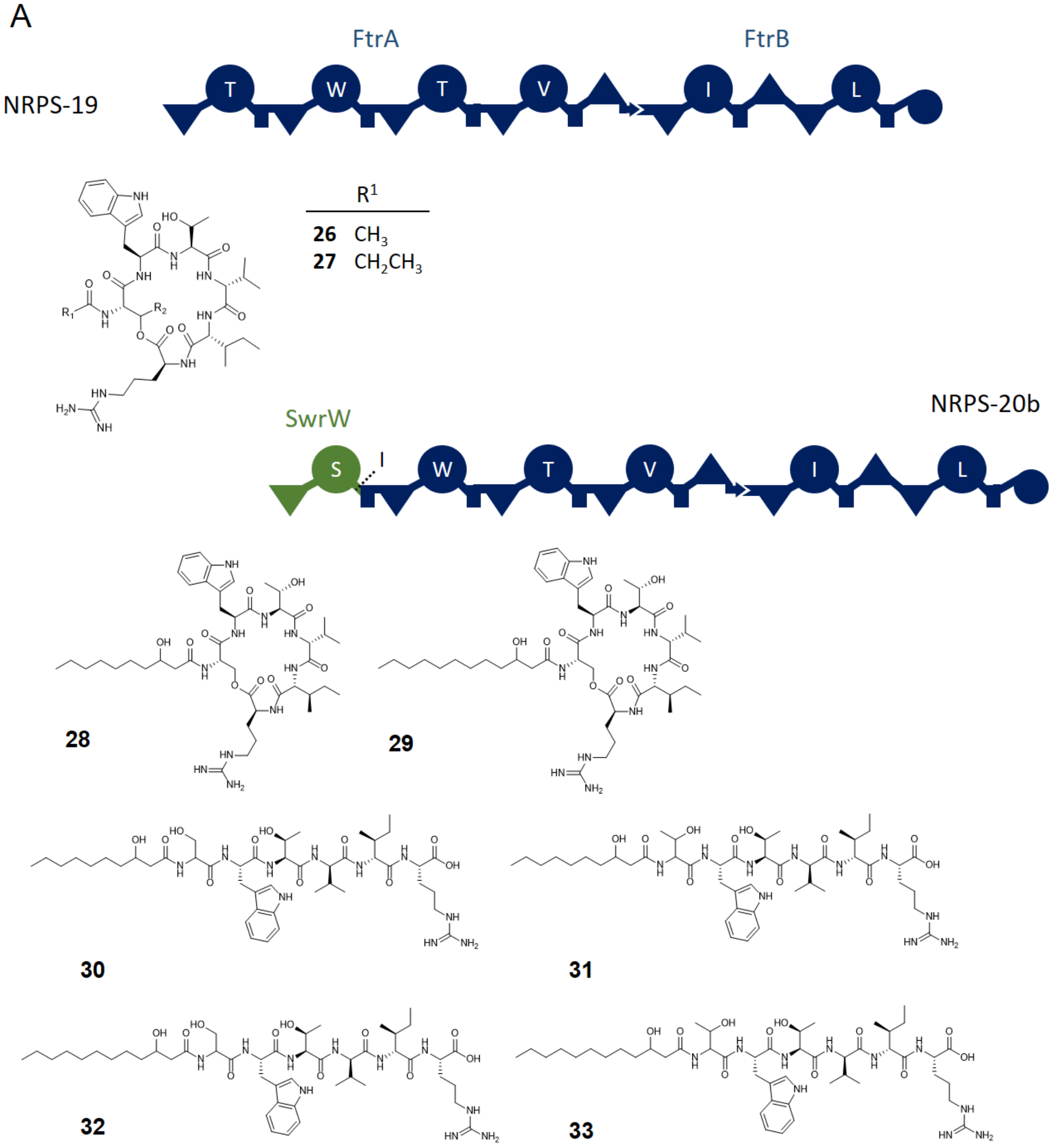


B

| NRPS | Peptide | Peptide | Organism | Donor BGC | Fusion site | Production $(mg\ l^{-1})$ | % of NRPS-8 |
|---|---|---|---|---|---|---|---|
| -19 | **26, 27** | C2-TWTviR | *X. mauleonii* | *ftrAB* | WT | 56.0 ±3.5 | 100 |
| -20b | **28 , 29** | C10-βOH-SWTviR | *S. marcescens* | *swrW* | IV | 2.5 ±0.2 | 4 |

**Figure S18.** (**A**) Domain architecture of Fatttvir (**FA** **T**hr **T**yr **T**hr **V**al **I**le a**R**g) producing FtrAB (NRPS-19) and NRPS-20b with their peptide product structures **26-33** shown below. Structure elucidation of **26** is shown at Figures S19 – S21 and S44 - S49. (**B**) For quantification the signal intensities for **28** and **29** were summarized and compared to the summarized amount of **26** and **27** in the WT.

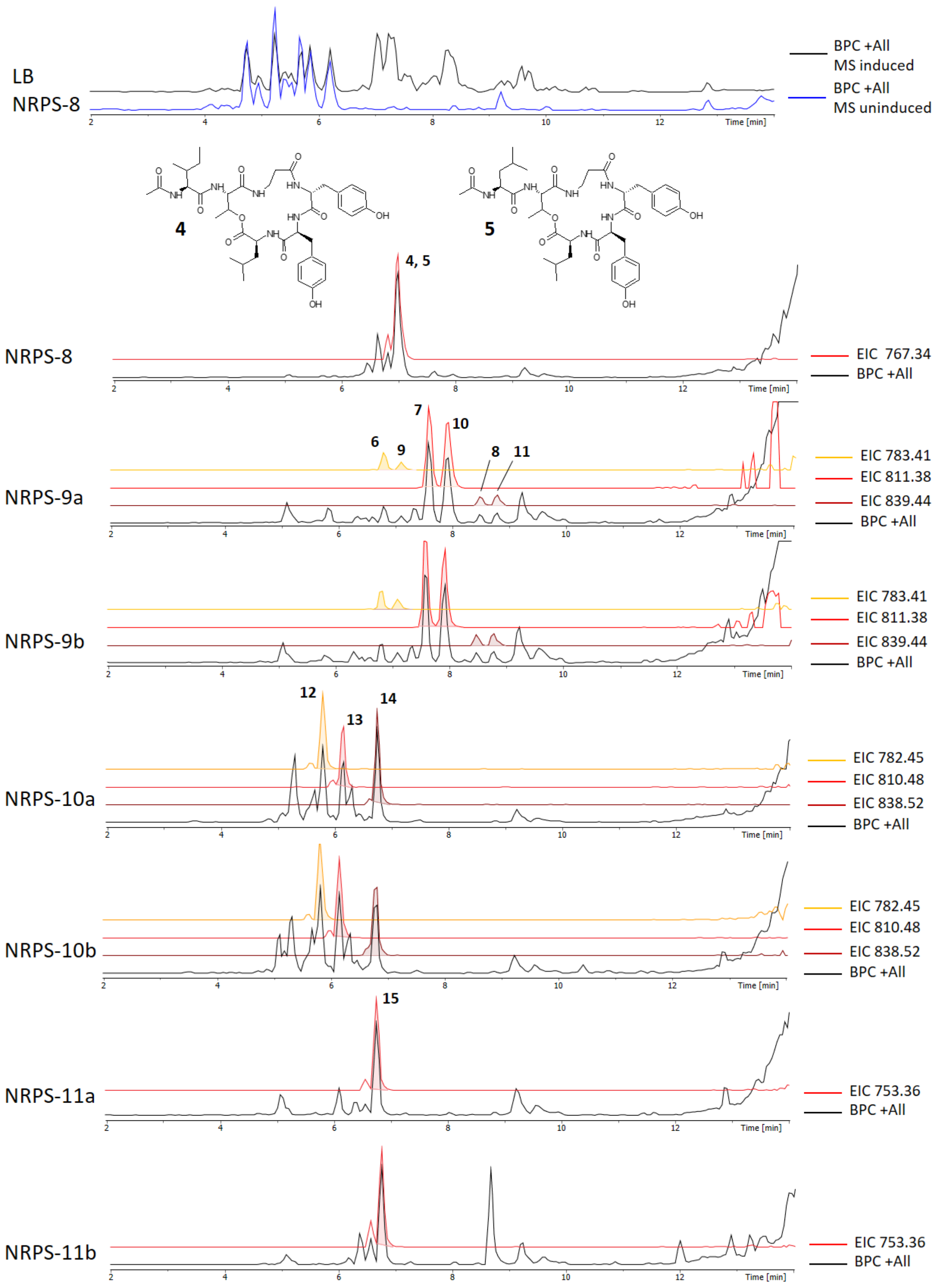
LB
NRPS-8
BPC +All
MS induced
BPC +All
MS uninduced
Time [min]
4
5
4, 5
NRPS-8
EIC 767.34
BPC +All
NRPS-9a
7
10
6
9
8
11
EIC 783.41
EIC 811.38
EIC 839.44
BPC +All
NRPS-9b
EIC 783.41
EIC 811.38
EIC 839.44
BPC +All
NRPS-10a
12
13
14
EIC 782.45
EIC 810.48
EIC 838.52
BPC +All
NRPS-10b
EIC 782.45
EIC 810.48
EIC 838.52
BPC +All
NRPS-11a
15
EIC 753.36
BPC +All
NRPS-11b
EIC 753.36
BPC +All

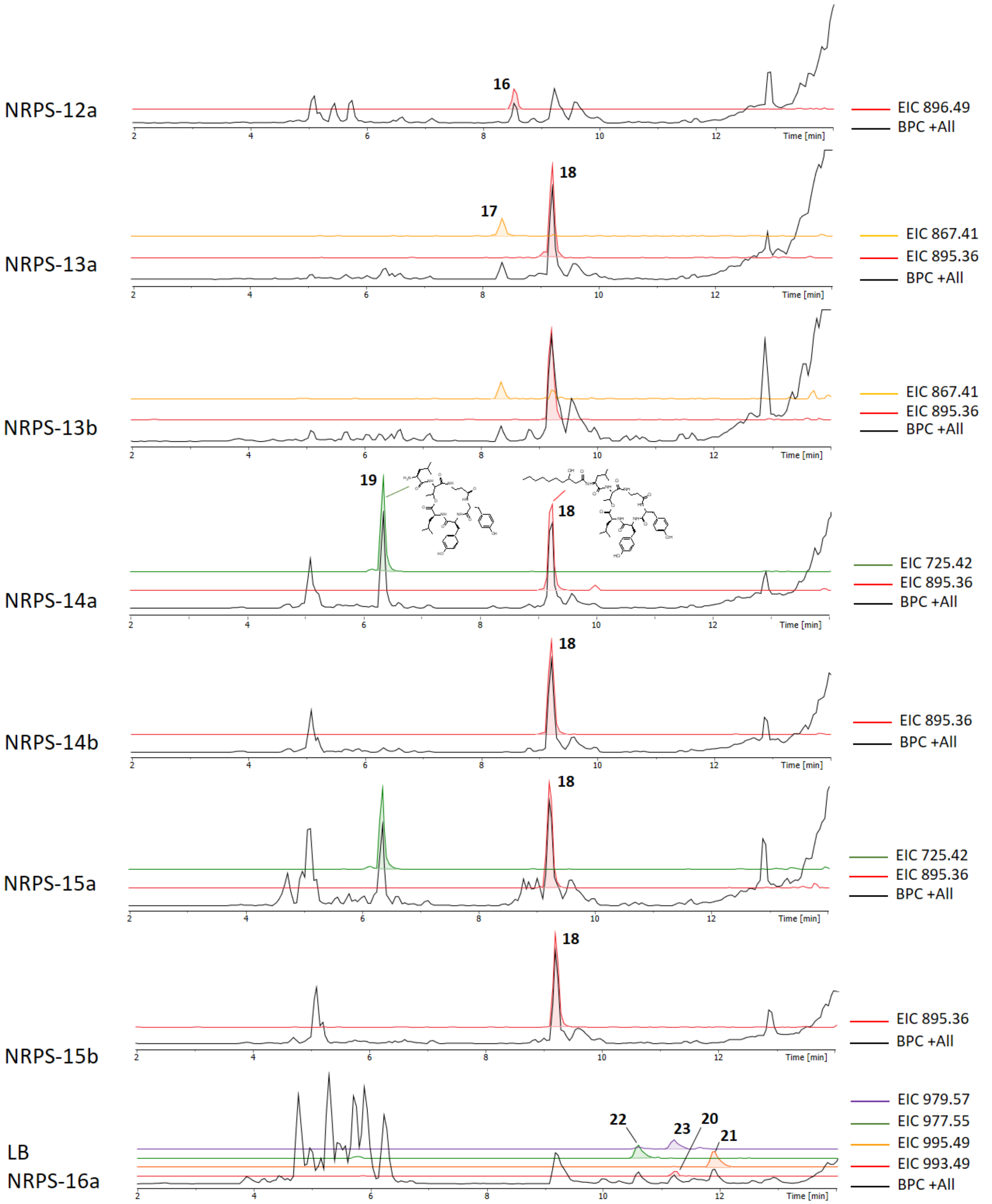
NRPS-12a
16
EIC 896.49
BPC +All
NRPS-13a
17
18
EIC 867.41
EIC 895.36
BPC +All
NRPS-13b
EIC 867.41
EIC 895.36
BPC +All
NRPS-14a
19
18
EIC 725.42
EIC 895.36
BPC +All
NRPS-14b
18
EIC 895.36
BPC +All
NRPS-15a
18
EIC 725.42
EIC 895.36
BPC +All
NRPS-15b
18
EIC 895.36
BPC +All
LB
NRPS-16a
22
23
20
21
EIC 979.57
EIC 977.55
EIC 995.49
EIC 993.49
BPC +All
Time [min]

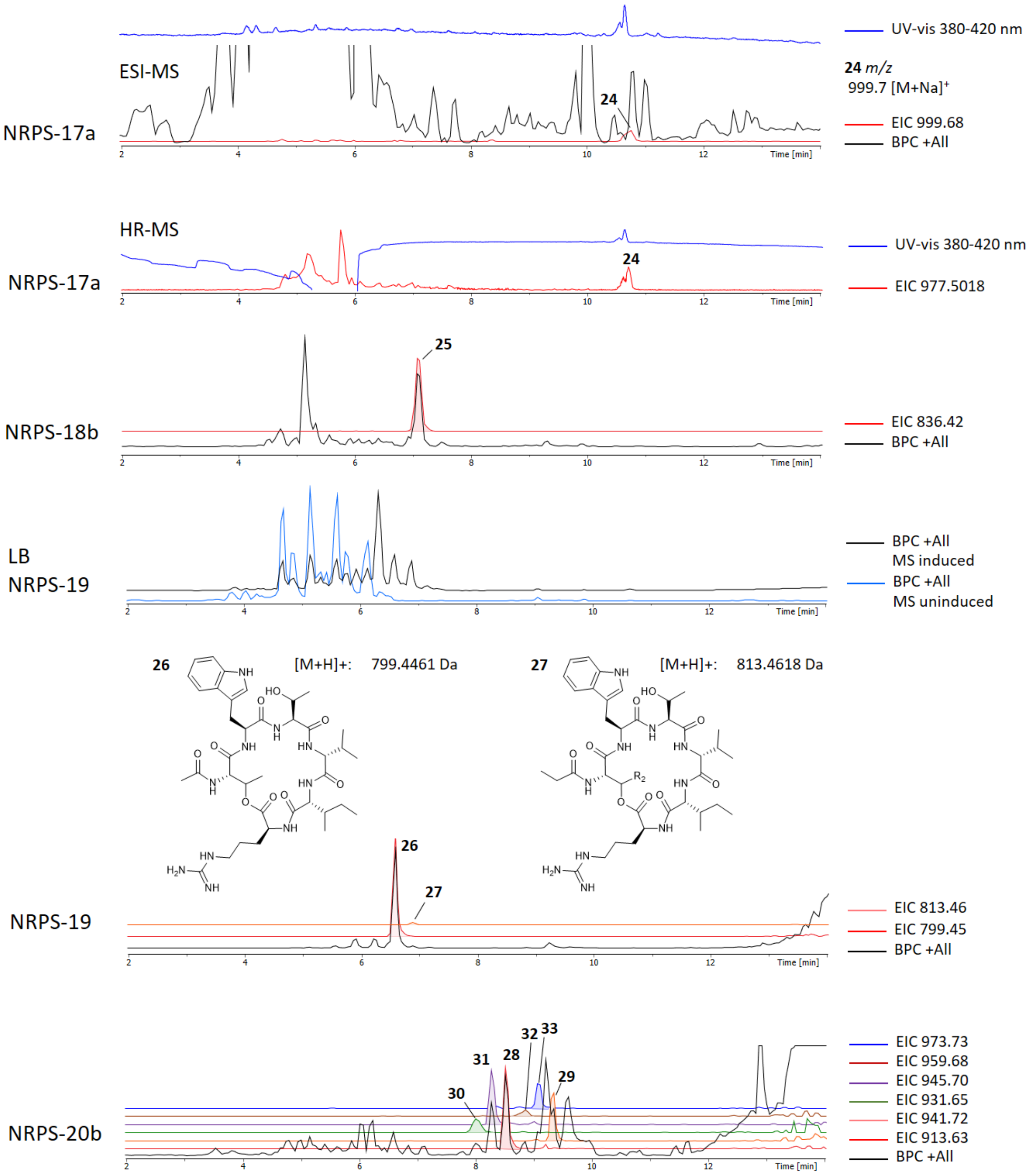


**Figure S19.** Chromatograms and structures of **4**, **5**, **26** and **27** and their NRPS-engineering derivatives.

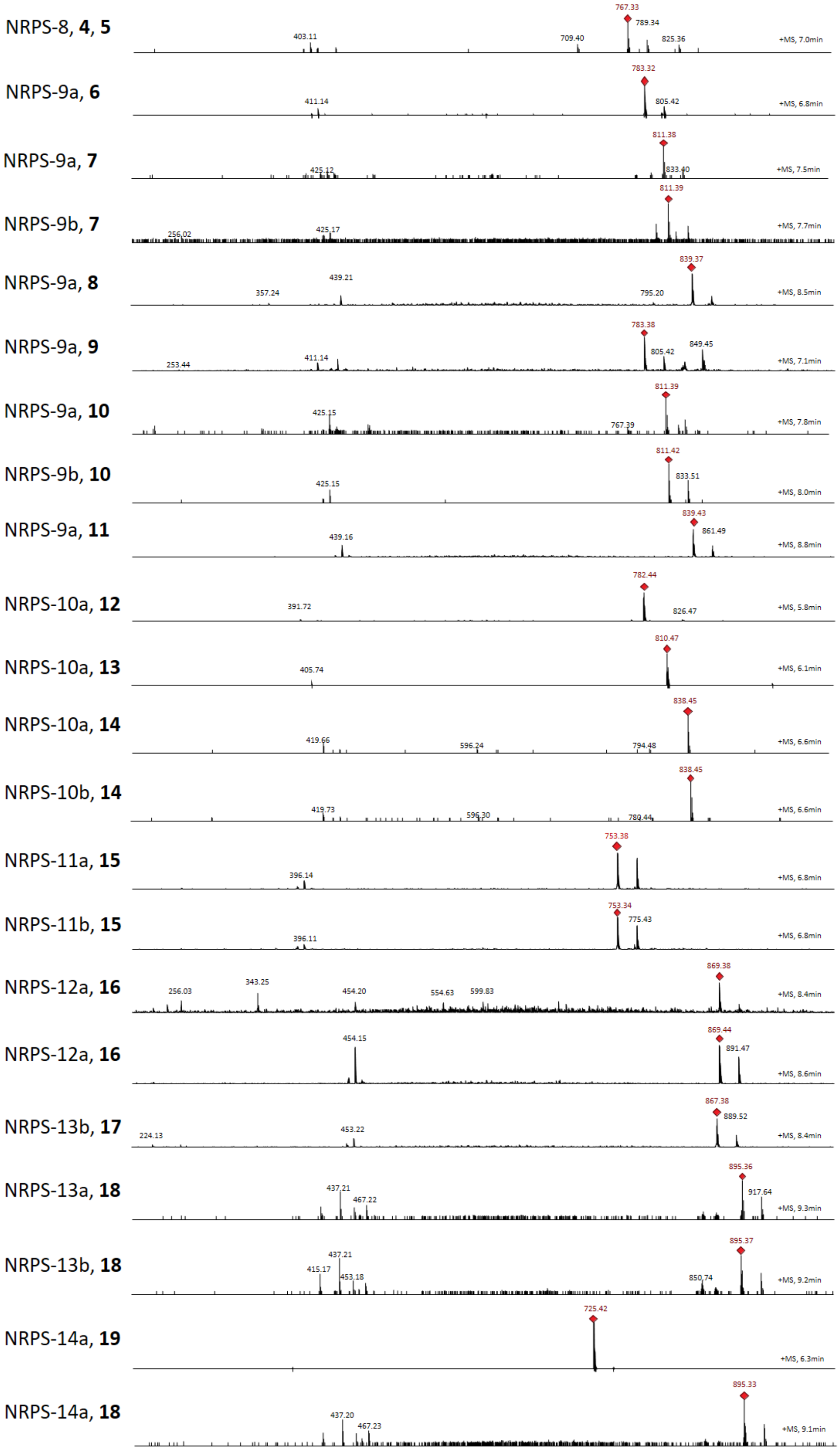

NRPS-8, **4**, **5**
767.33
789.34
403.11
709.40
825.36
+MS, 7.0min
NRPS-9a, **6**
783.32
411.14
805.42
+MS, 6.8min
NRPS-9a, **7**
811.38
425.12
833.40
+MS, 7.5min
NRPS-9b, **7**
811.39
256.02
425.17
+MS, 7.7min
NRPS-9a, **8**
839.37
439.21
357.24
795.20
+MS, 8.5min
NRPS-9a, **9**
783.38
849.45
805.42
253.44
411.14
+MS, 7.1min
NRPS-9a, **10**
811.39
425.15
767.39
+MS, 7.8min
NRPS-9b, **10**
811.42
833.51
425.15
+MS, 8.0min
NRPS-9a, **11**
839.43
861.49
439.16
+MS, 8.8min
NRPS-10a, **12**
782.44
391.72
826.47
+MS, 5.8min
NRPS-10a, **13**
810.47
405.74
+MS, 6.1min
NRPS-10a, **14**
838.45
419.66
596.24
794.48
+MS, 6.6min
NRPS-10b, **14**
838.45
419.73
596.30
780.44
+MS, 6.6min
NRPS-11a, **15**
753.38
396.14
+MS, 6.8min
NRPS-11b, **15**
753.34
775.43
396.11
+MS, 6.8min
NRPS-12a, **16**
869.38
343.25
256.03
454.20
554.63
599.83
+MS, 8.4min
NRPS-12a, **16**
869.44
454.15
891.47
+MS, 8.6min
NRPS-13b, **17**
867.38
889.52
453.22
224.13
+MS, 8.4min
NRPS-13a, **18**
895.36
437.21
917.64
467.22
+MS, 9.3min
NRPS-13b, **18**
895.37
437.21
415.17
453.18
850.74
+MS, 9.2min
NRPS-14a, **19**
725.42
+MS, 6.3min
NRPS-14a, **18**
895.33
437.20
467.23
+MS, 9.1min

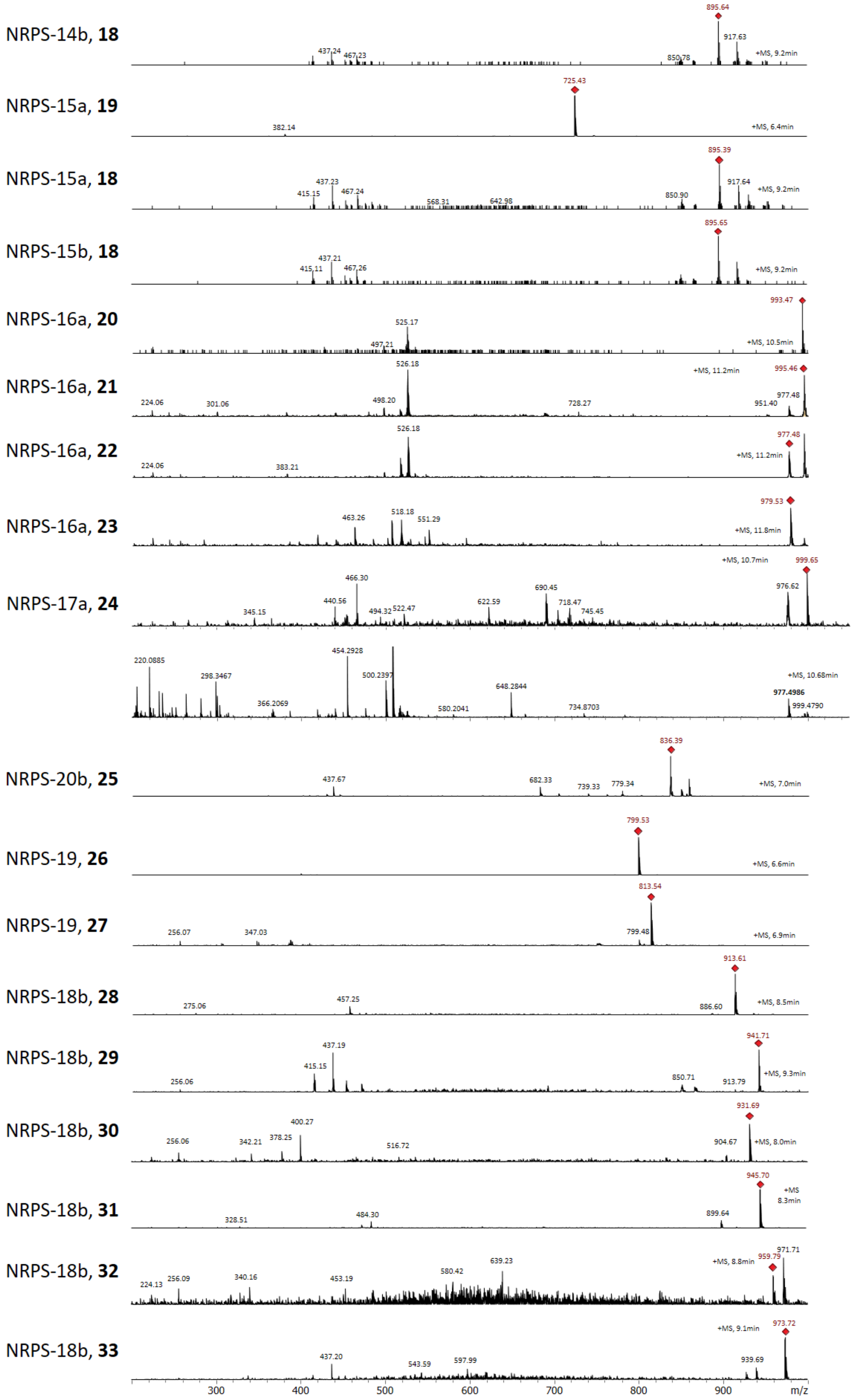


**Figure S20**. MS-spectra of peptides **4-33** of NRPS-8 to -20 corresponding to the extracted ion chromatograms in **Fig. S19**.

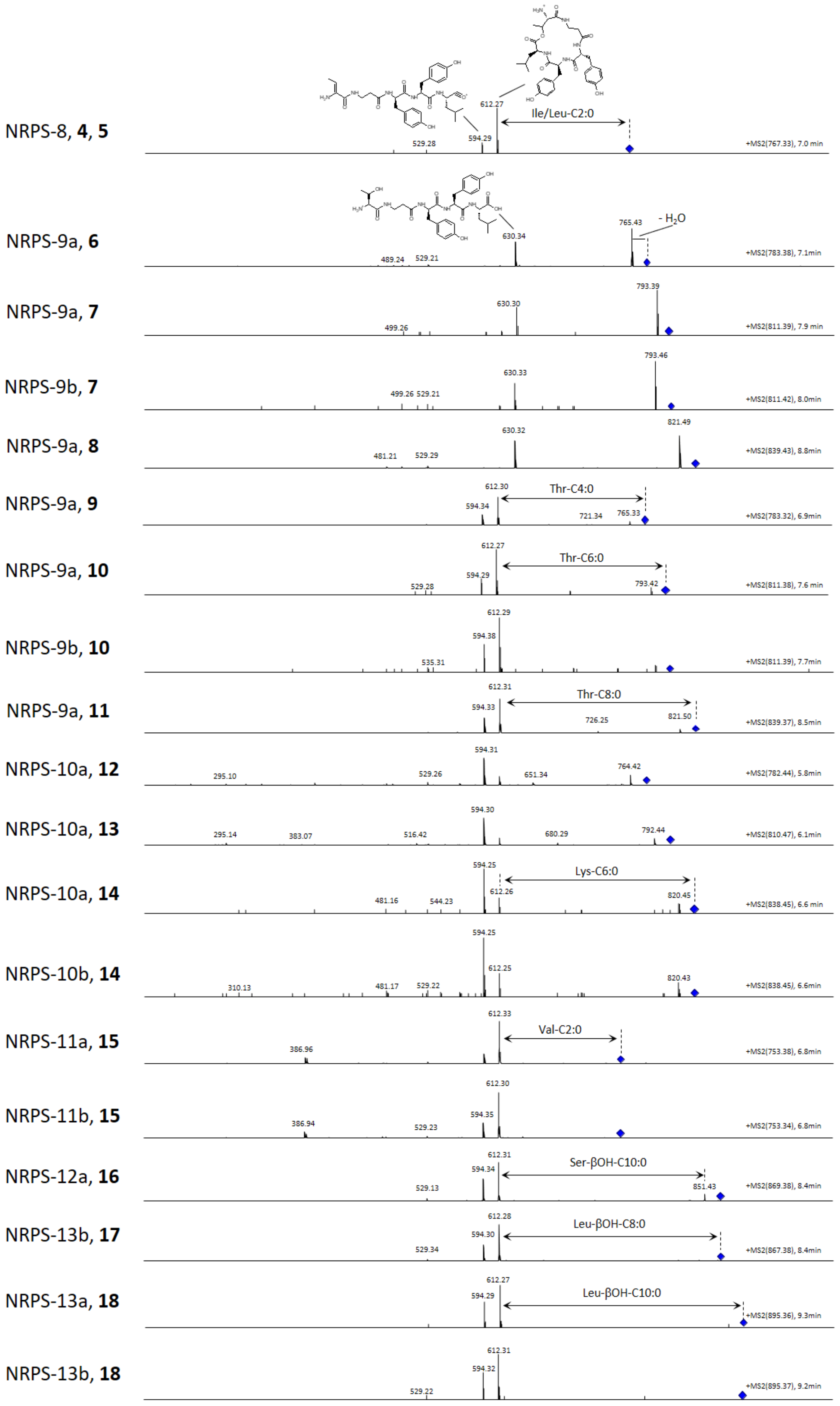

NRPS-8, 4, 5
612.27
Ile/Leu-C2:0
529.28
594.29
+MS2(767.33), 7.0 min
NRPS-9a, 6
765.43
- $H_2O$
630.34
489.24
529.21
+MS2(783.38), 7.1min
NRPS-9a, 7
793.39
630.30
499.26
+MS2(811.39), 7.9 min
NRPS-9b, 7
793.46
630.33
499.26 529.21
+MS2(811.42), 8.0min
NRPS-9a, 8
821.49
630.32
481.21
529.29
+MS2(839.43), 8.8min
NRPS-9a, 9
612.30
Thr-C4:0
594.34
721.34
765.33
+MS2(783.32), 6.9min
NRPS-9a, 10
612.27
Thr-C6:0
594.29
529.28
793.42
+MS2(811.38), 7.6 min
NRPS-9b, 10
612.29
594.38
535.31
+MS2(811.39), 7.7min
NRPS-9a, 11
612.31
Thr-C8:0
594.33
726.25
821.50
+MS2(839.37), 8.5min
NRPS-10a, 12
594.31
295.10
529.26
651.34
764.42
+MS2(782.44), 5.8min
NRPS-10a, 13
594.30
295.14
383.07
516.42
680.29
792.44
+MS2(810.47), 6.1min
NRPS-10a, 14
594.25
Lys-C6:0
612.26
481.16
544.23
820.45
+MS2(838.45), 6.6 min
NRPS-10b, 14
594.25
612.25
310.13
481.17
529.22
820.43
+MS2(838.45), 6.6min
NRPS-11a, 15
612.33
Val-C2:0
386.96
+MS2(753.38), 6.8min
NRPS-11b, 15
612.30
594.35
386.94
529.23
+MS2(753.34), 6.8min
NRPS-12a, 16
612.31
Ser-βOH-C10:0
594.34
529.13
851.43
+MS2(869.38), 8.4min
NRPS-13b, 17
612.28
Leu-βOH-C8:0
594.30
529.34
+MS2(867.38), 8.4min
NRPS-13a, 18
612.27
Leu-βOH-C10:0
594.29
+MS2(895.36), 9.3min
NRPS-13b, 18
612.31
594.32
529.22
+MS2(895.37), 9.2min

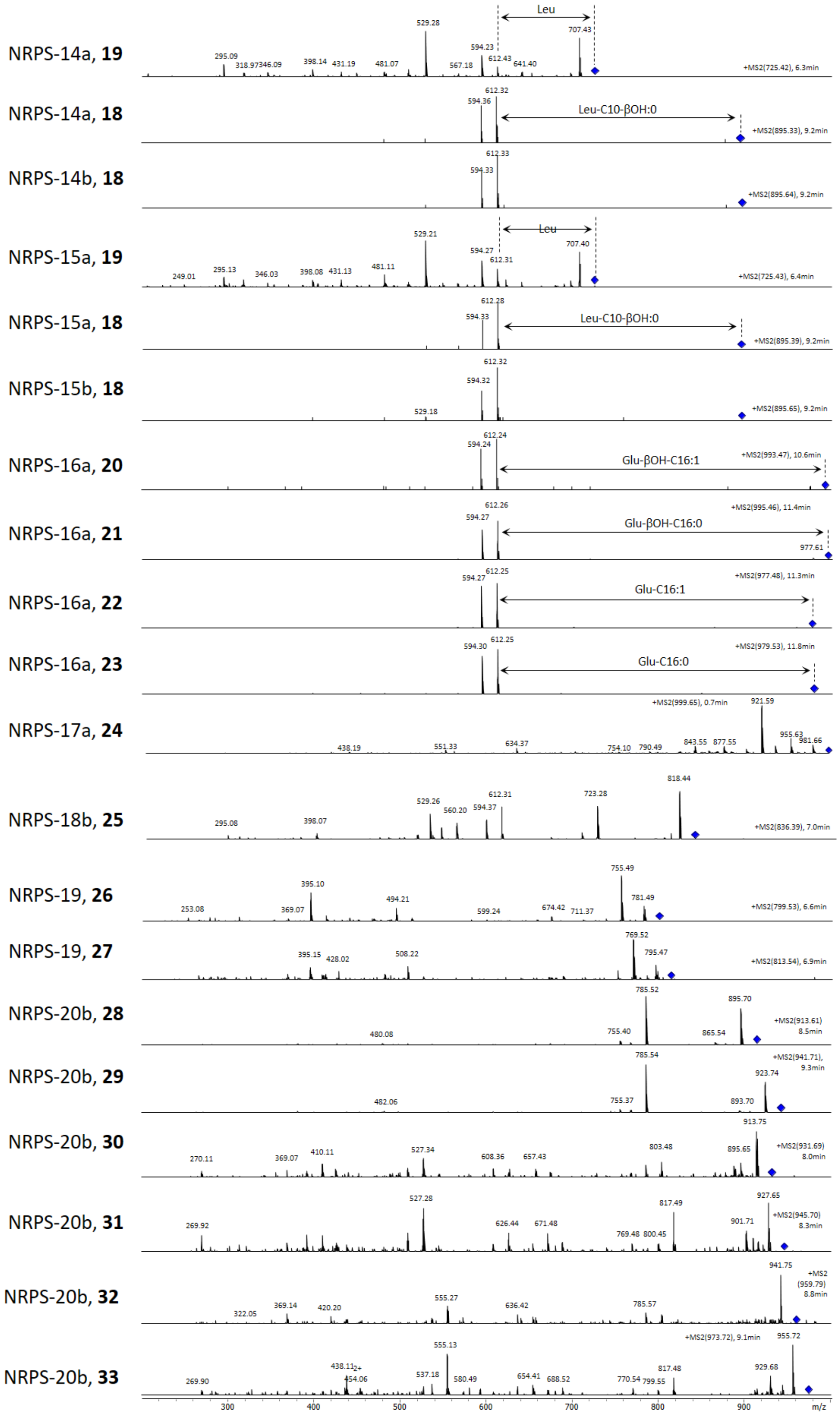


**Figure S21**. MS$^2$ spectra of peptides **4-33** of NRPS-8 to -20 corresponding to the signals in **Fig. S20**.

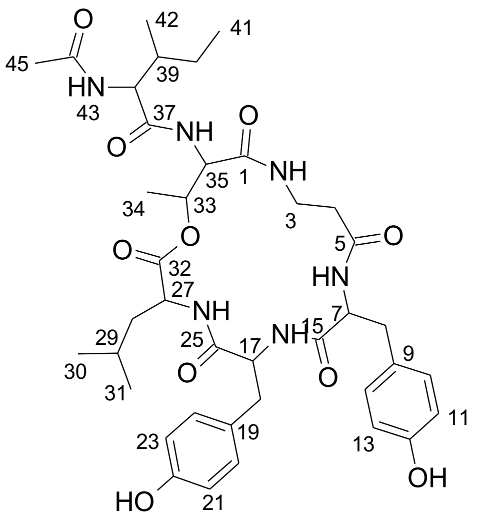


**4**

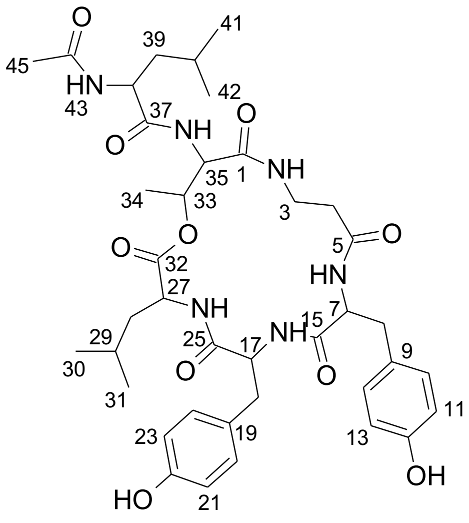


**5**

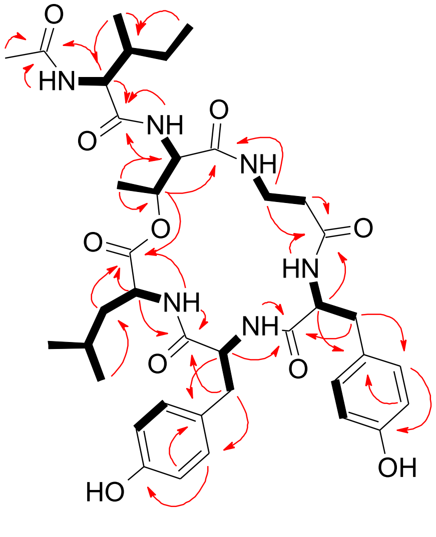


**4**

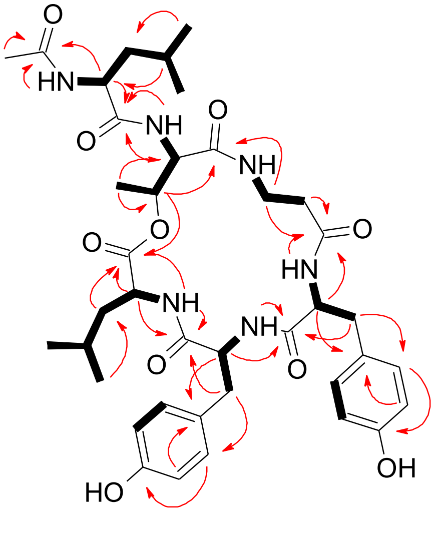


**5**

HMBC H→ C    1H-1H COSY H━ H

**Figure S22.** Key HMBC and $^{1}$H-$^{1}$H COSY correlations of compounds **4** and **5**.

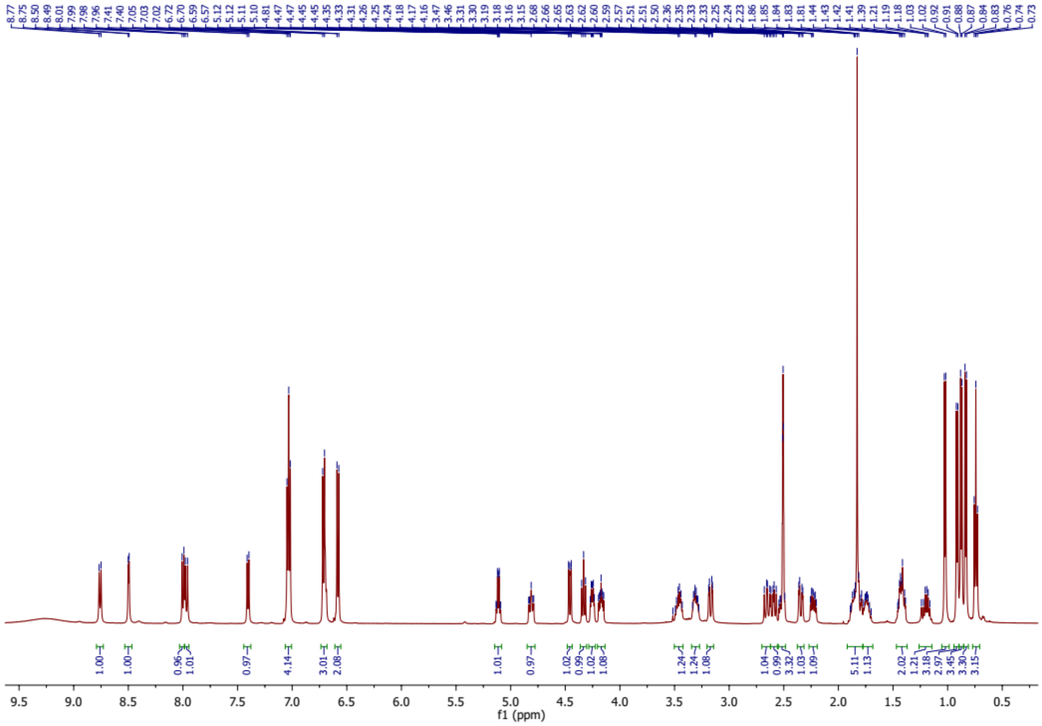

**Figure S23.** $^{1}H$ NMR spectrum of compound **4**.

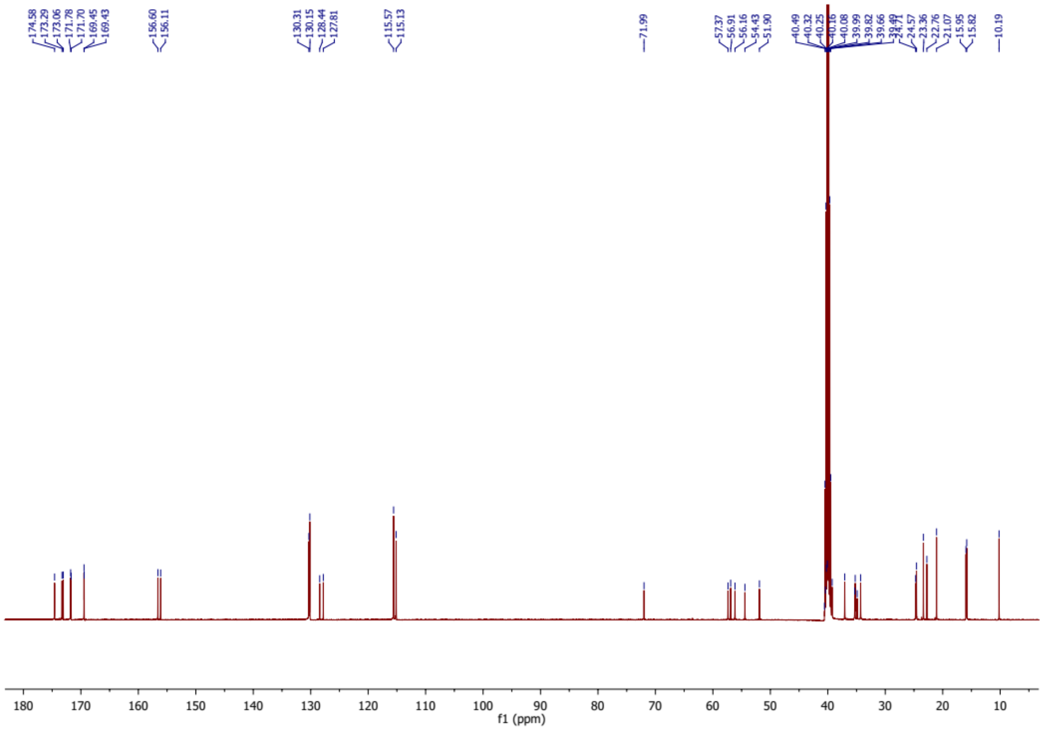

**Figure S24.** $^{13}C$ NMR spectrum of compound **4.**

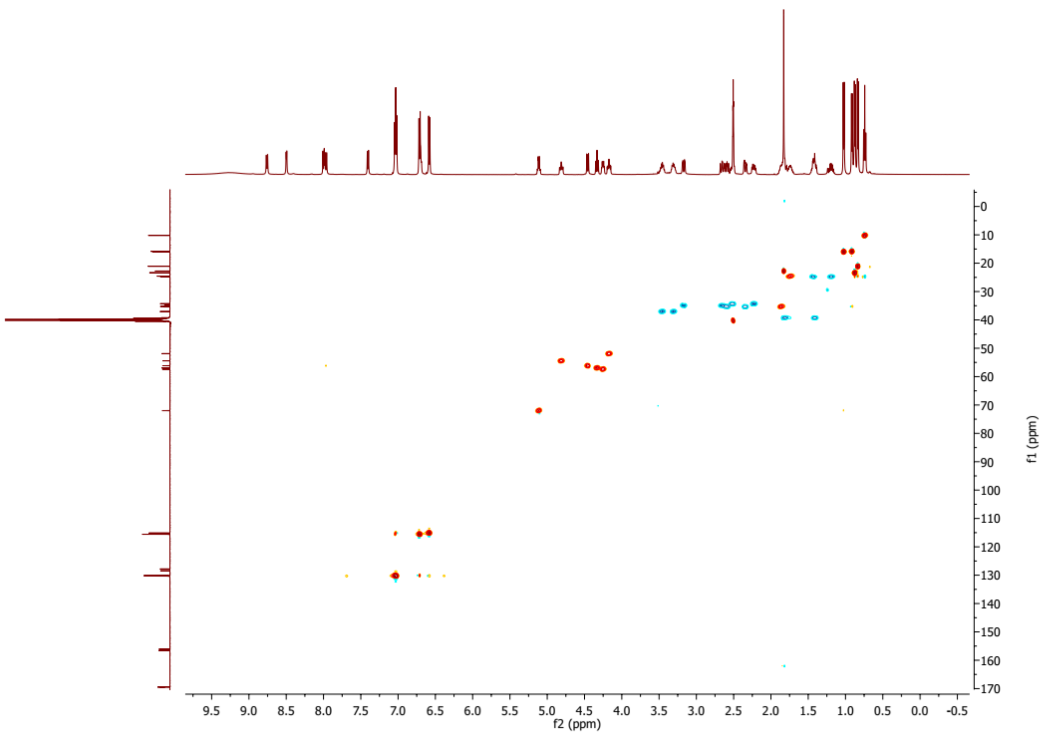


**Figure S25.** HSQC spectrum of compound **4**.

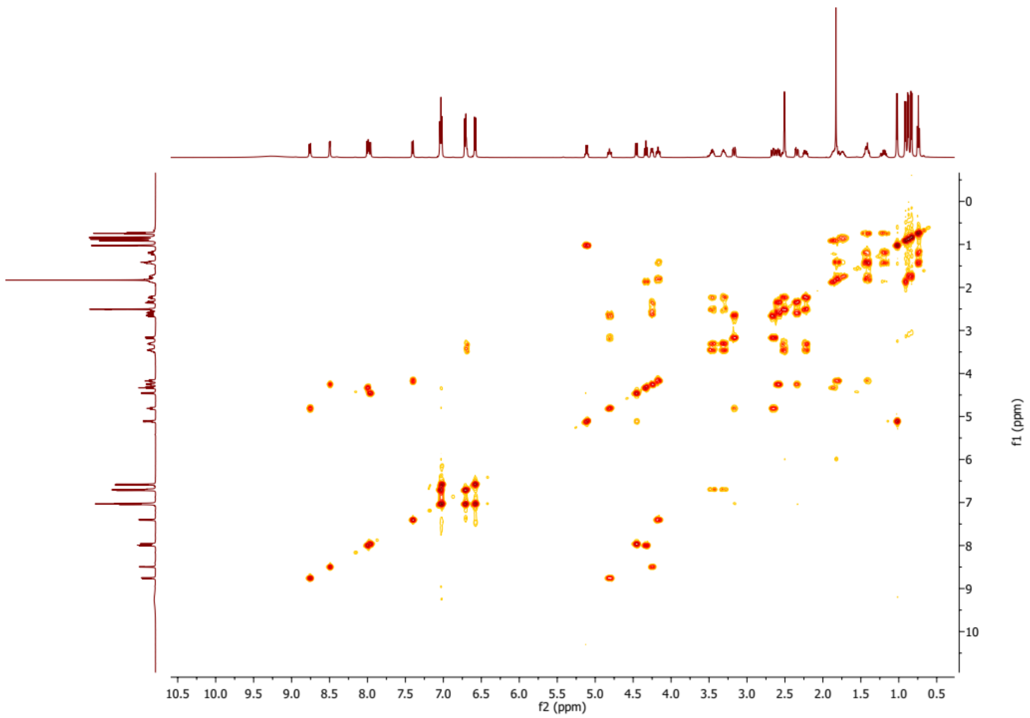


**Figure S26.** $^1H$-$^1H$ COSY spectrum of compound **4**.

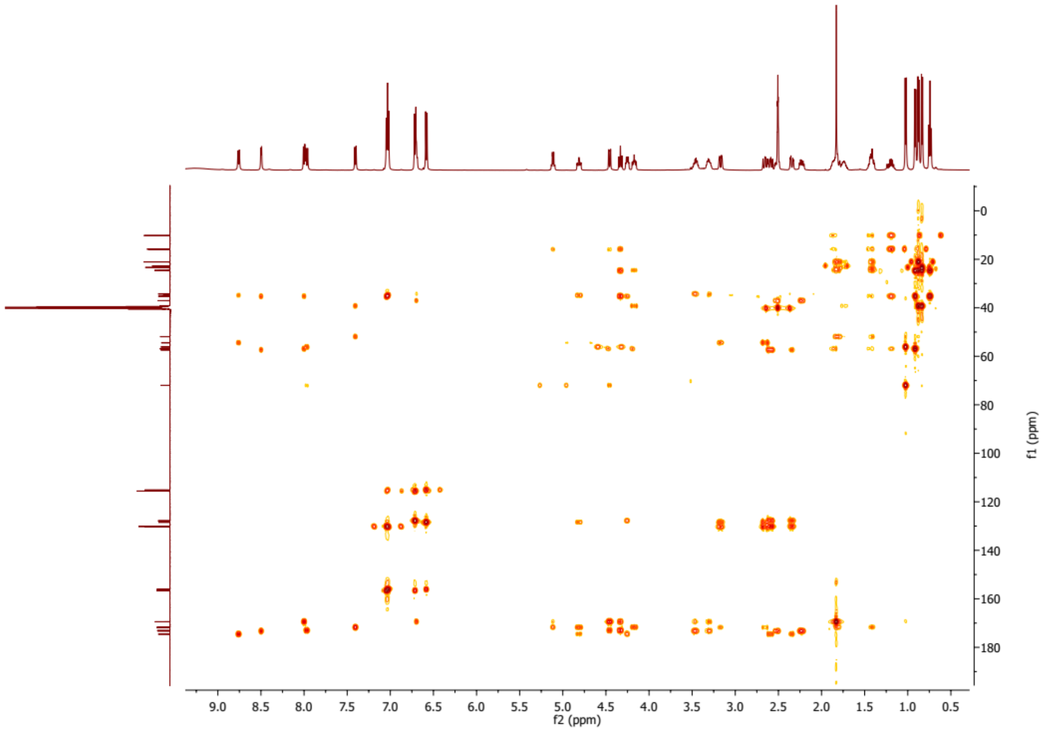

**Figure S27.** HMBC spectrum of compound **4**.

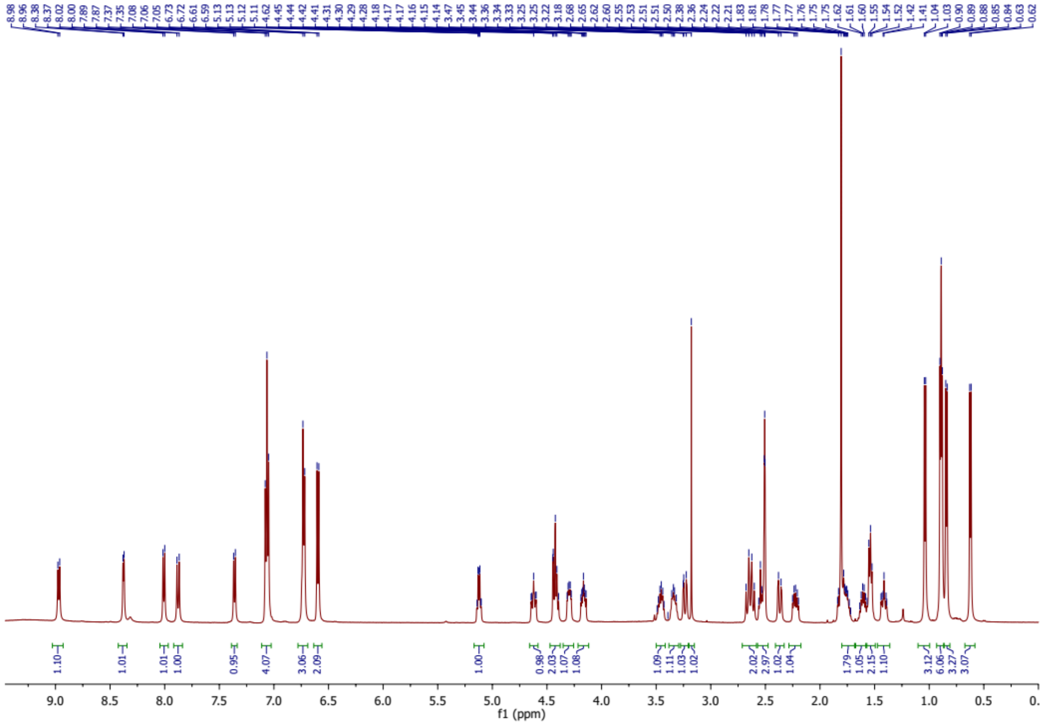

**Figure S28.** [1]H NMR spectrum of compound **5**.

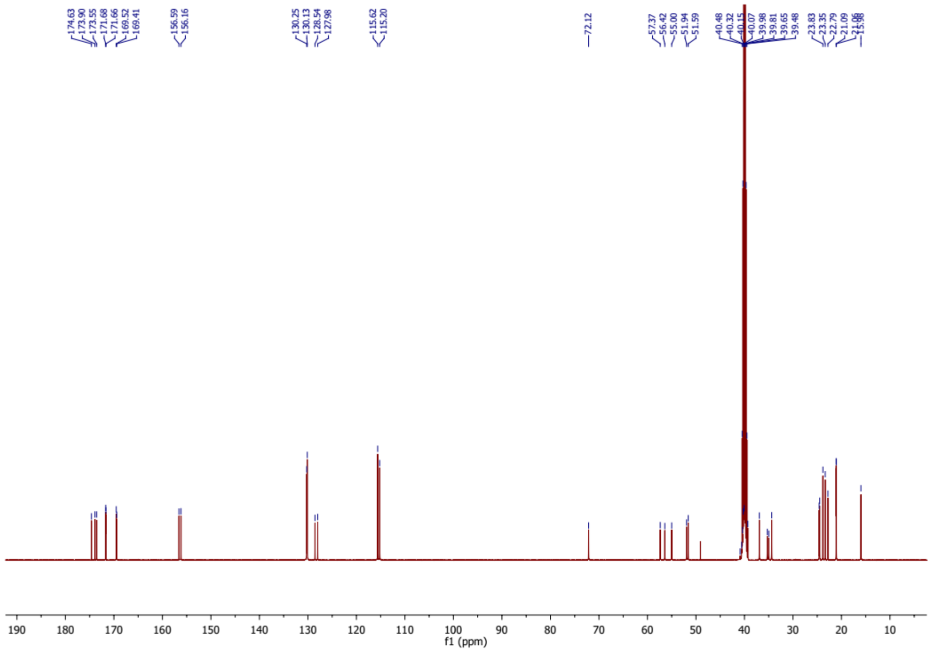

**Figure S29.** $^{13}$C NMR spectrum of compound **5**.

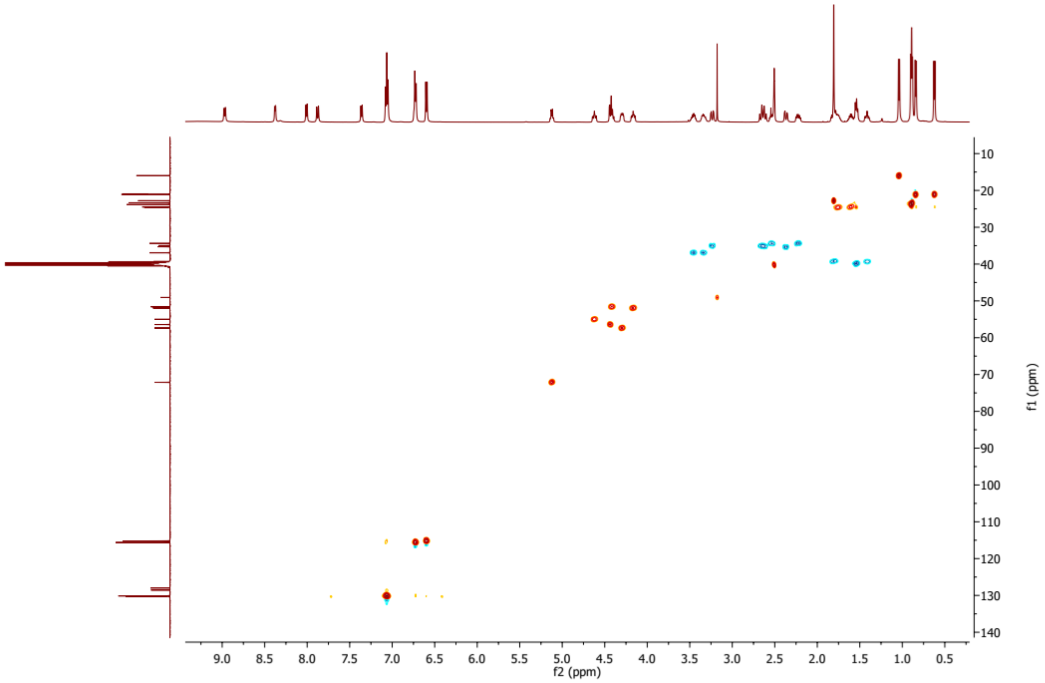

**Figure S30.** HSQC spectrum of compound **5**.

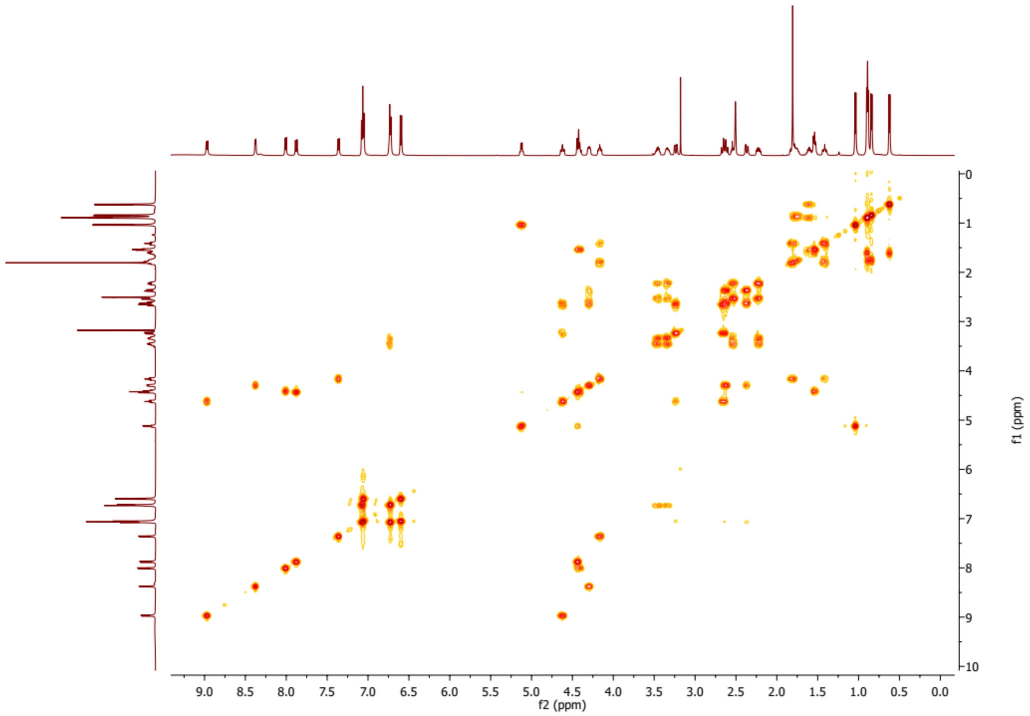


**Figure S31.** $^{1}H$-$^{1}H$ COSY spectrum of compound **5**.

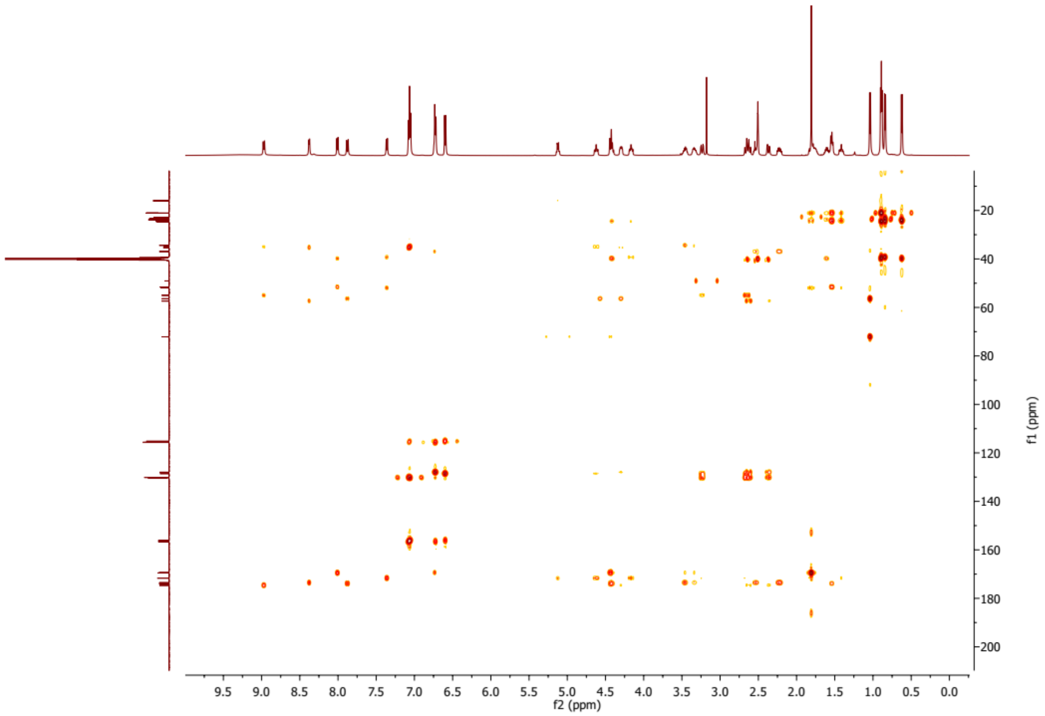


**Figure S32.** HMBC spectrum of compound **5**.

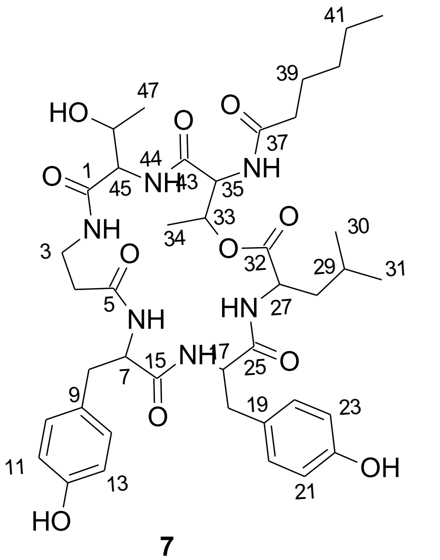

**7**

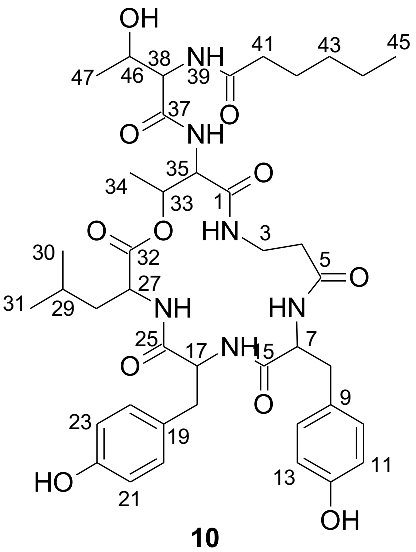

**10**

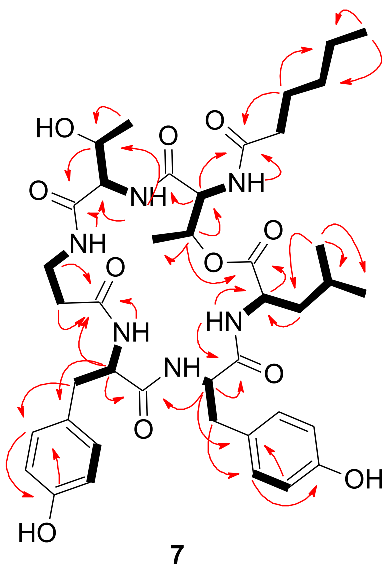

**7**

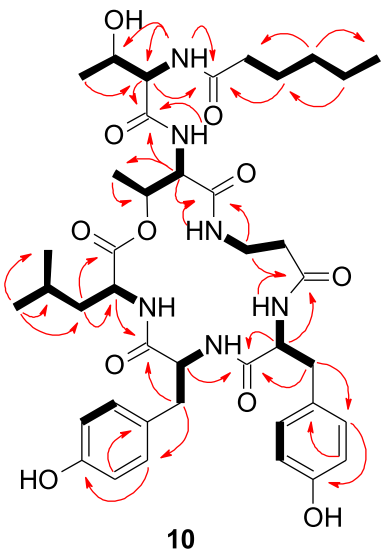

**10**

HMBC H→ C  1H-1H COSY H▬ H

**Figure S33.** Key HMBC and $^1$H-$^1$H COSY correlations of compounds **7** and **10**.

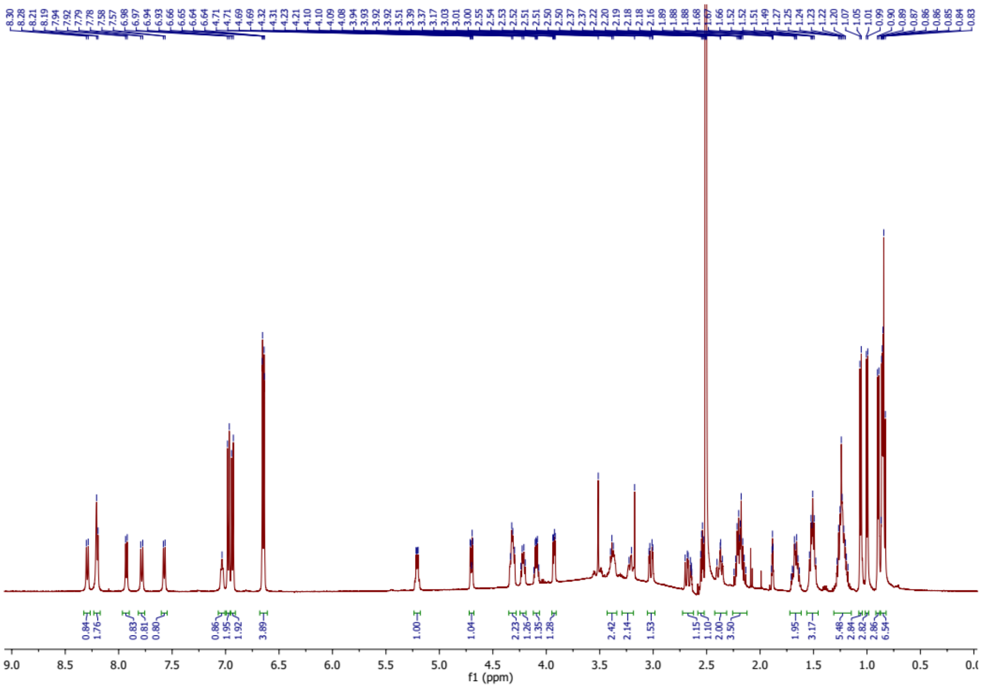

**Figure S34.** $^{1}$H NMR spectrum of compound **7.**

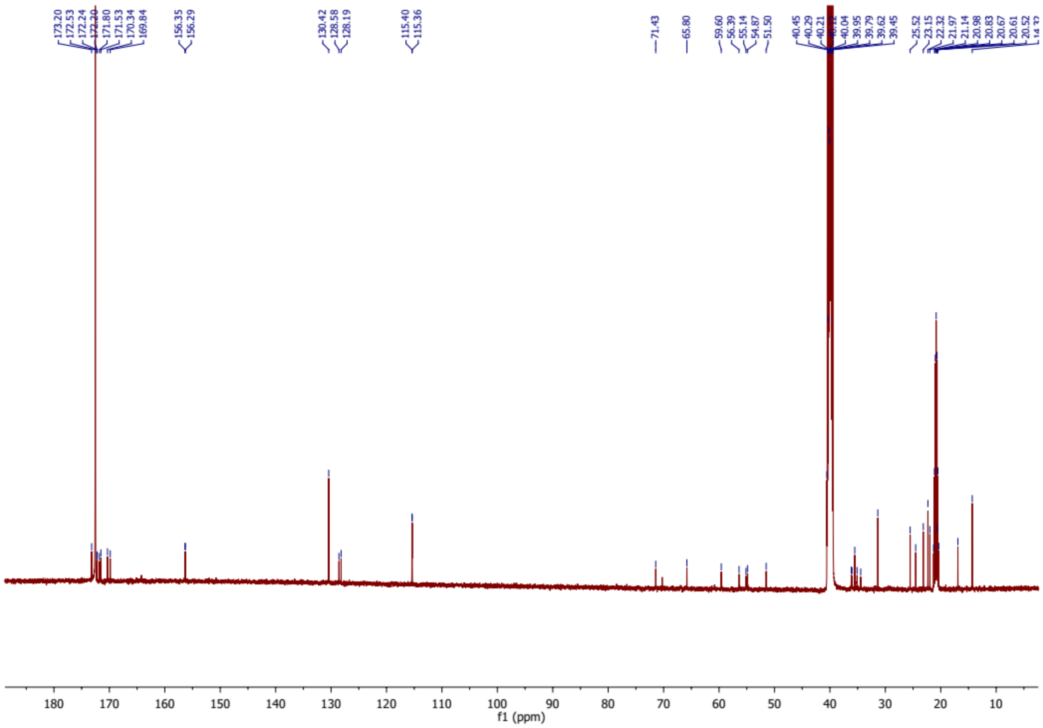

**Figure S35.** $^{13}$C NMR spectrum of compound **7**.

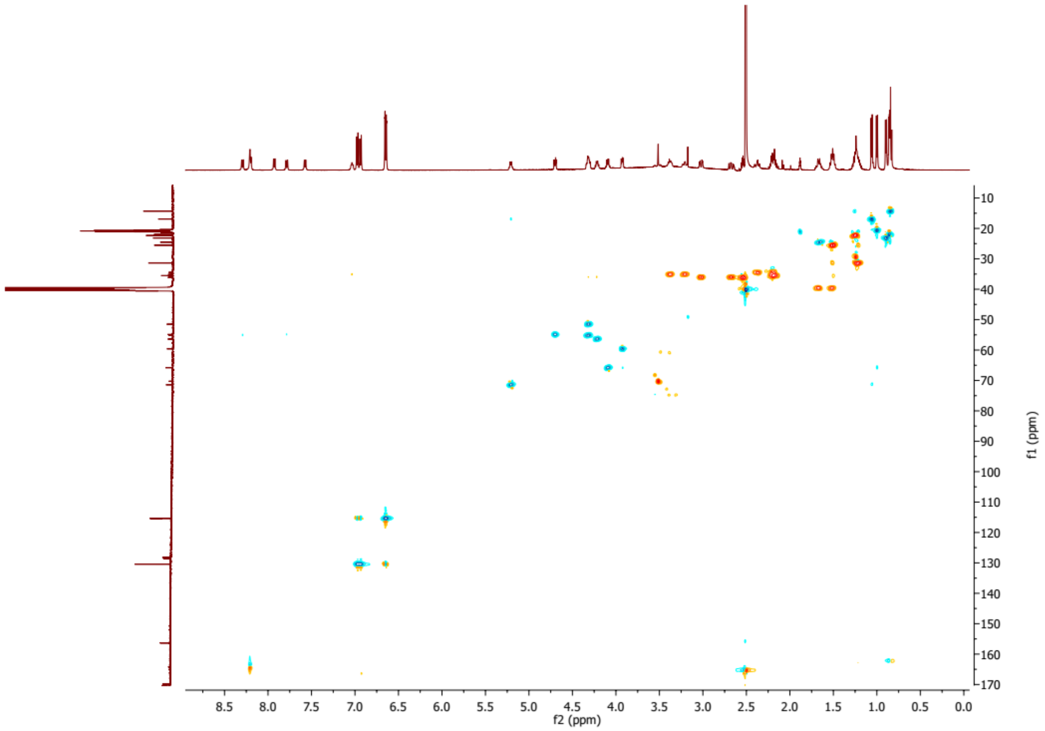


**Figure S36.** HSQC spectrum of compound **7**.

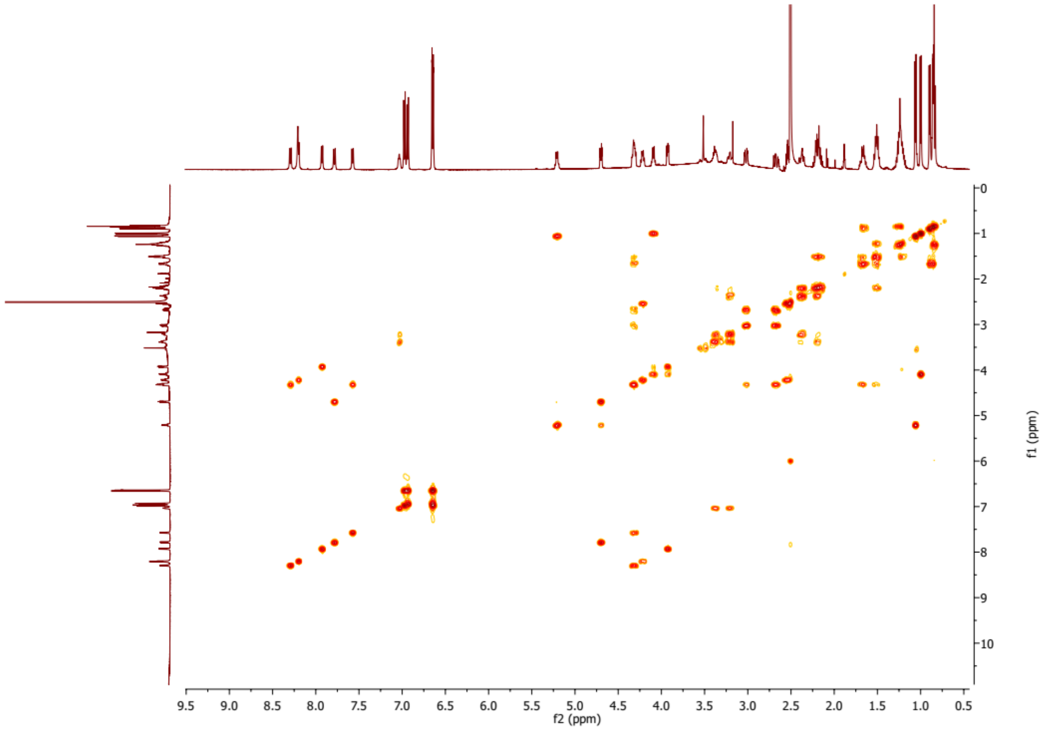


**Figure S37.** $^{1}H$-$^{1}H$ COSY spectrum of compound **7**.

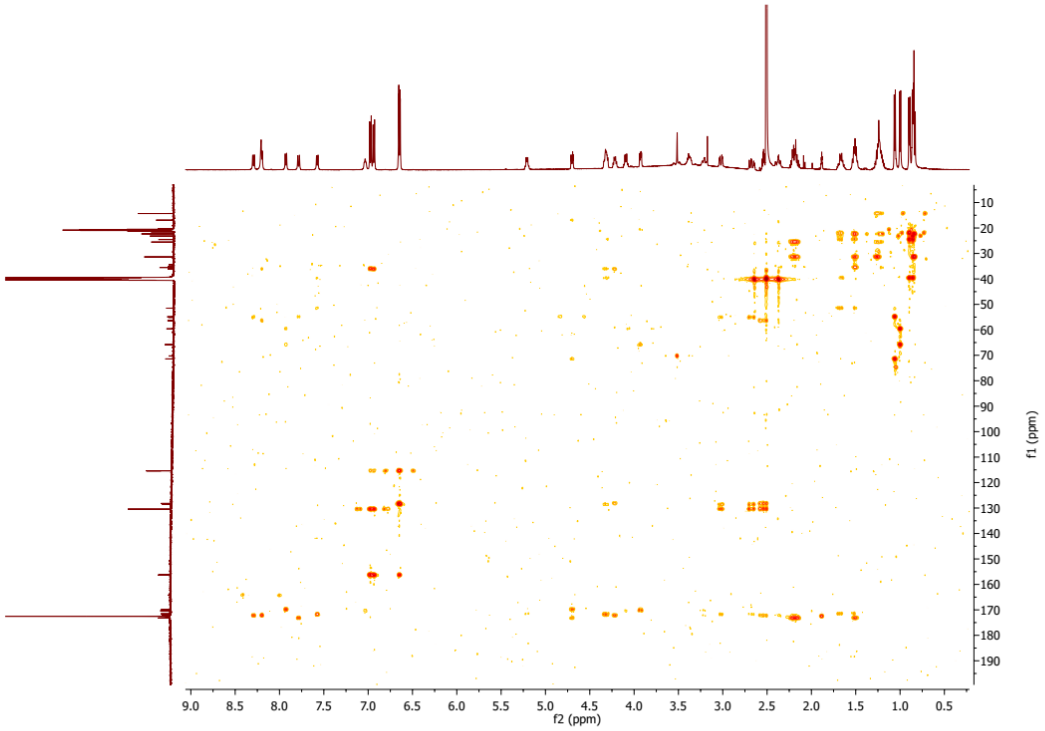

**Figure S38.** HMBC spectrum of compound **7**.

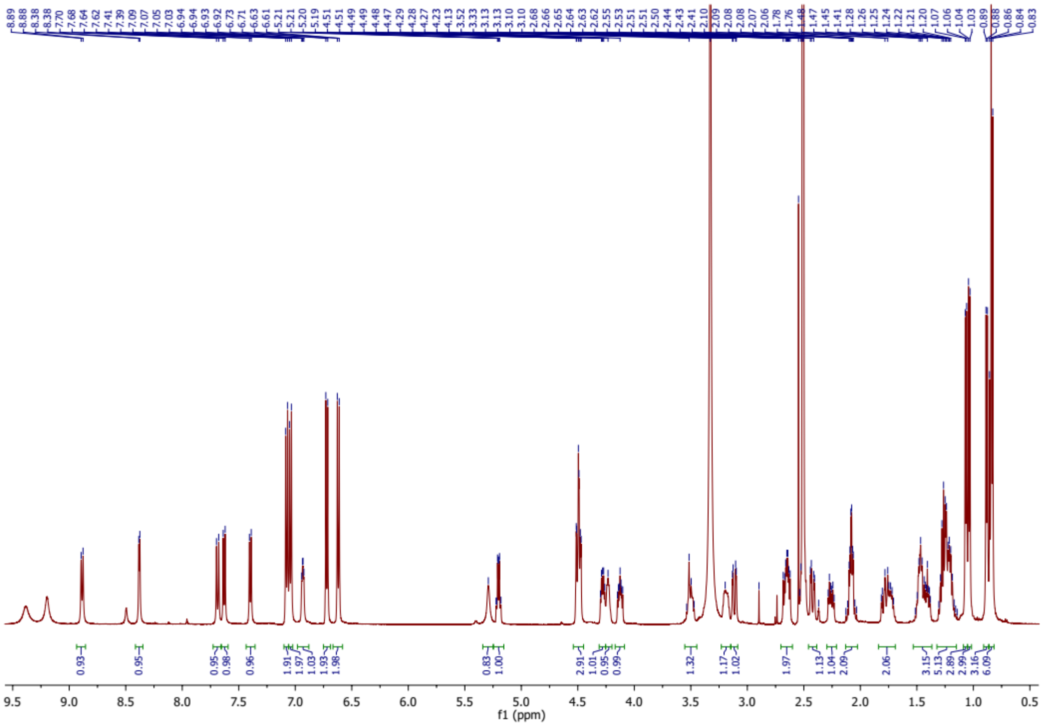

**Figure S39.** [1]H NMR spectrum of compound **10**.

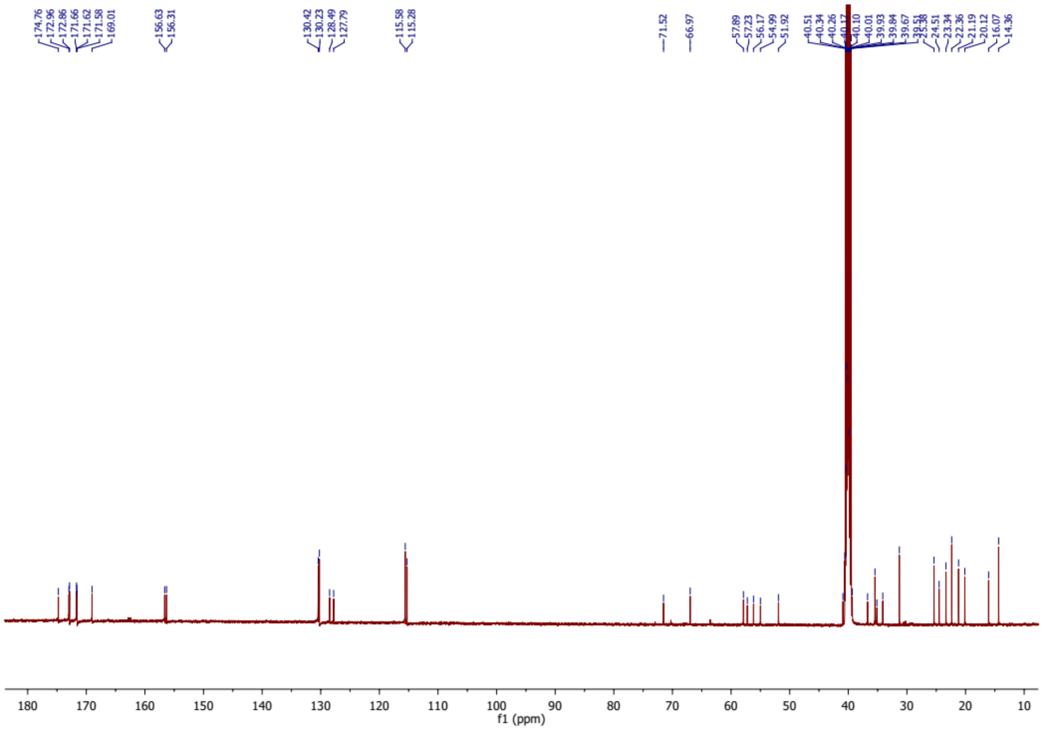

**Figure S40.** $^{13}C$ NMR spectrum of compound **10**.

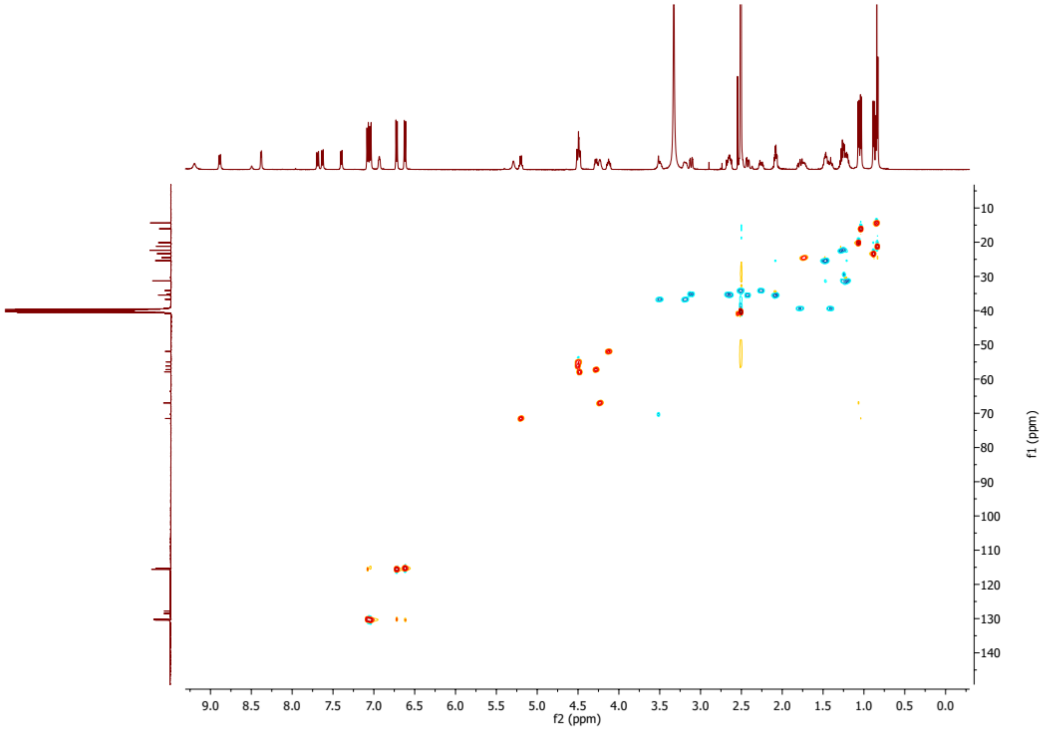

**Figure S41.** HSQC spectrum of compound **10**.

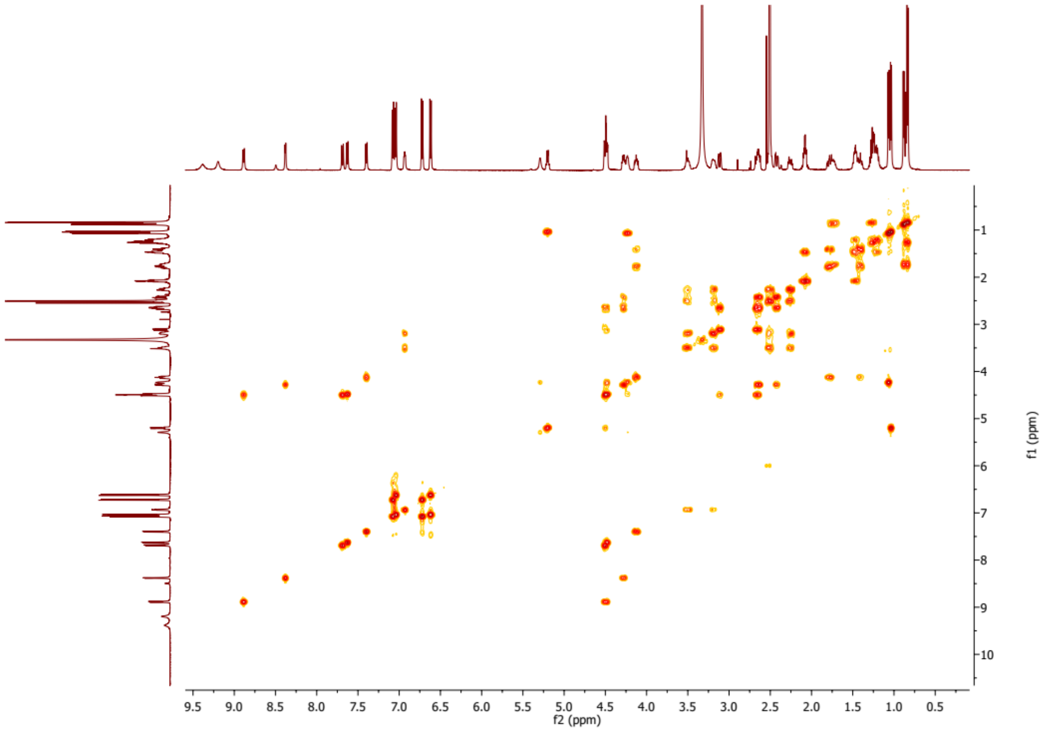


**Figure S42.** $^{1}H$-$^{1}H$ COSY spectrum of compound **10**.

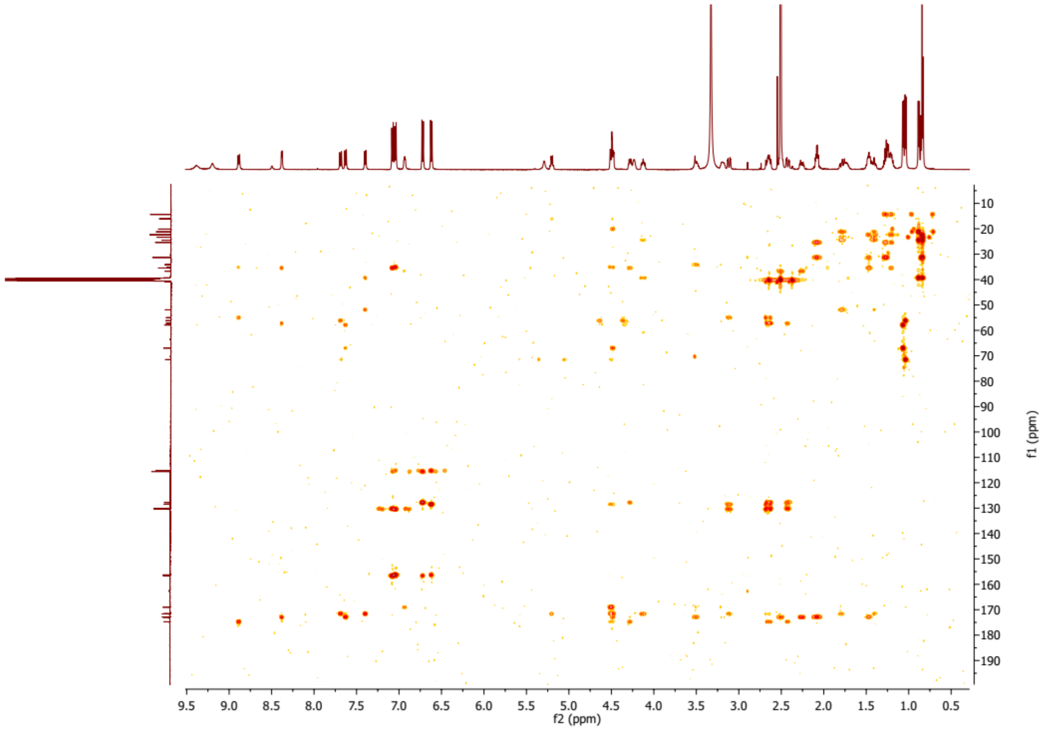


**Figure S43.** HMBC spectrum of compound **10**.

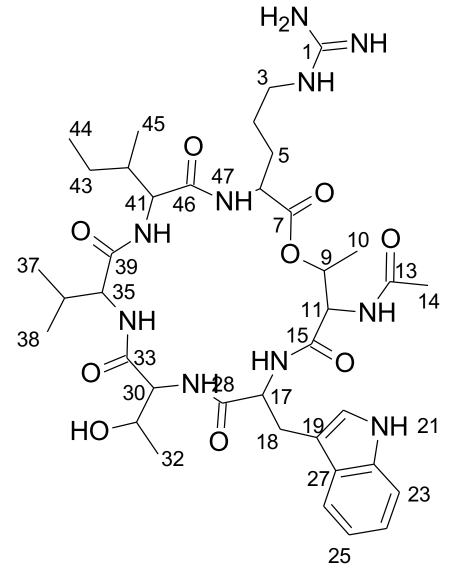


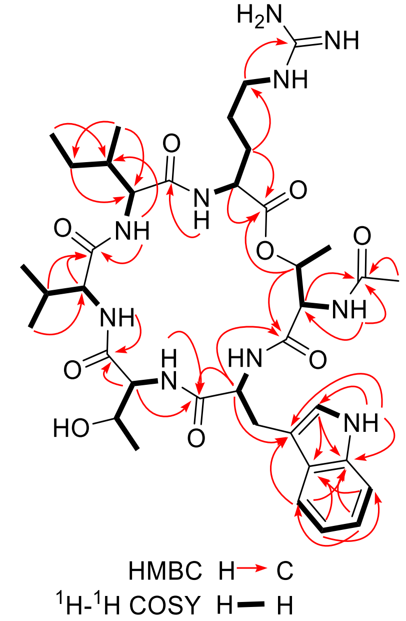


**Figure S44.** Key HMBC and $^1$H-$^1$H COSY correlations of compound **26**.

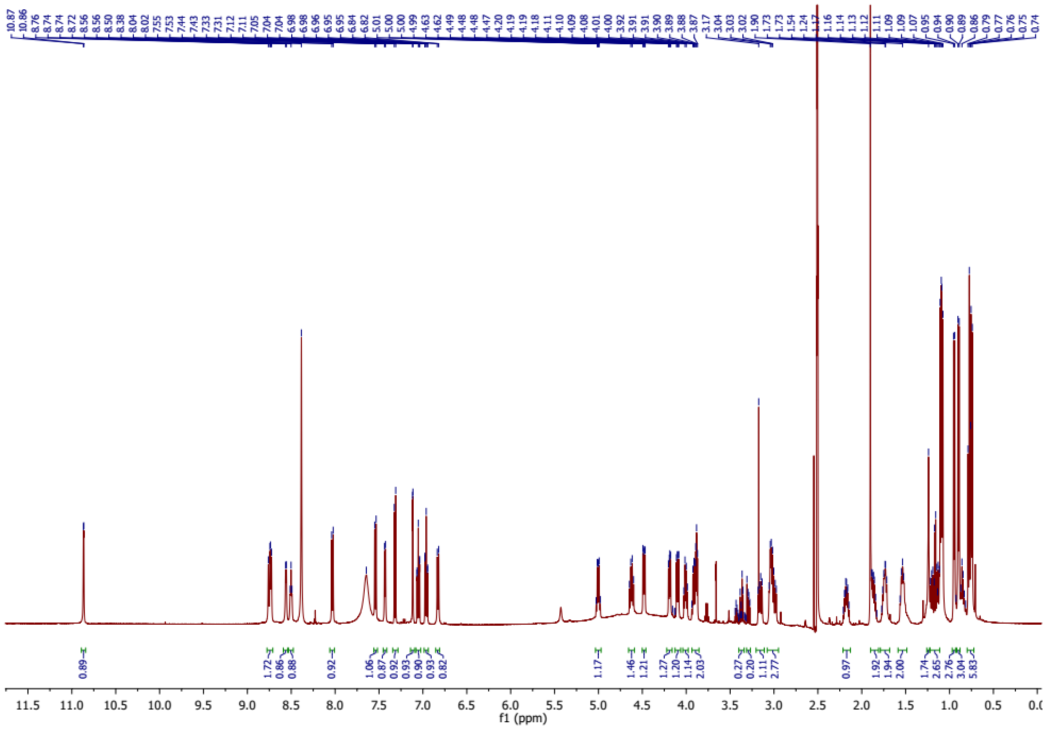


**Figure S45.** $^{1}$H NMR spectrum of compound **26**.

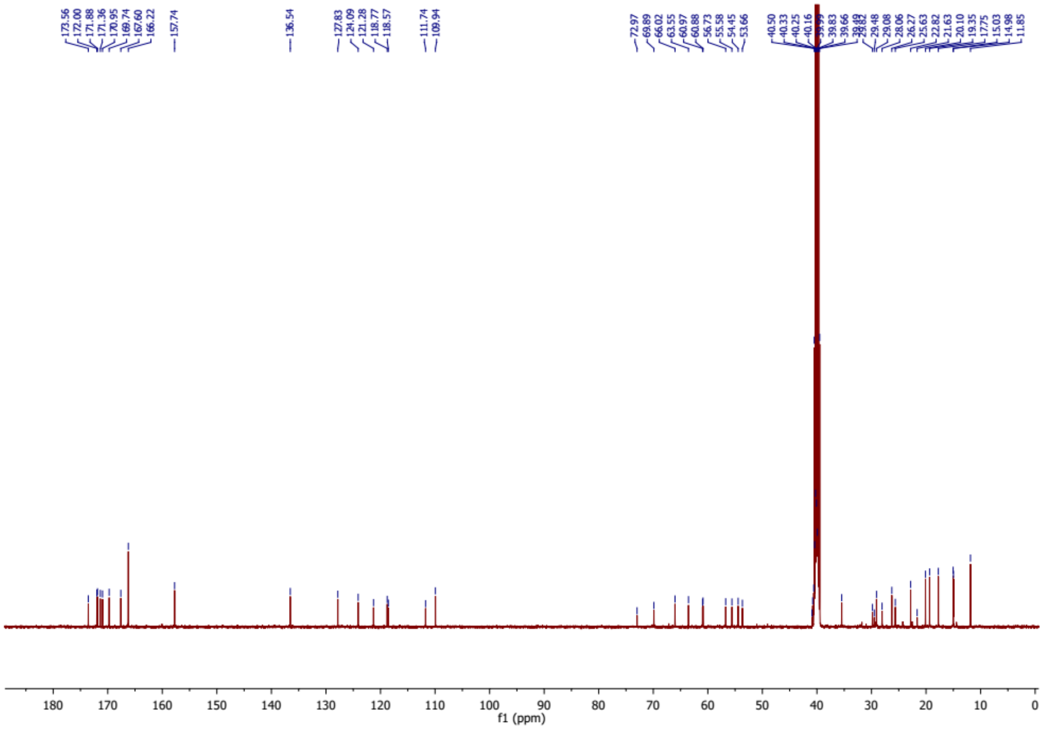


**Figure S46.** $^{13}$C NMR spectrum of compound **26**.

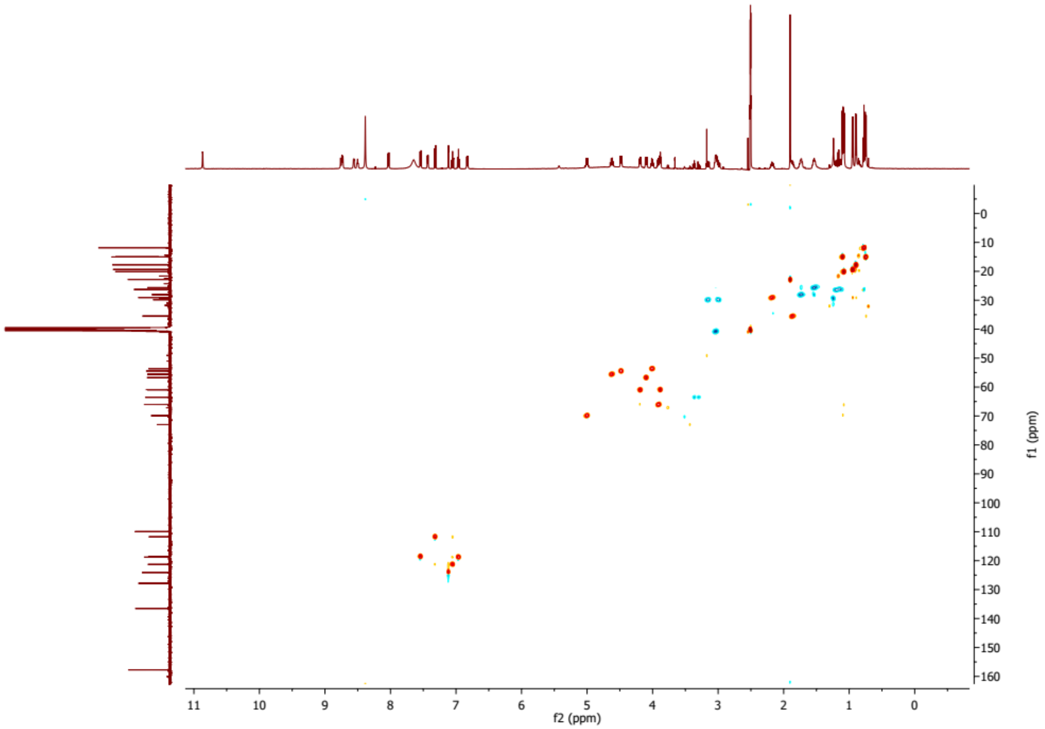


**Figure S47.** HSQC spectrum of compound **26**.

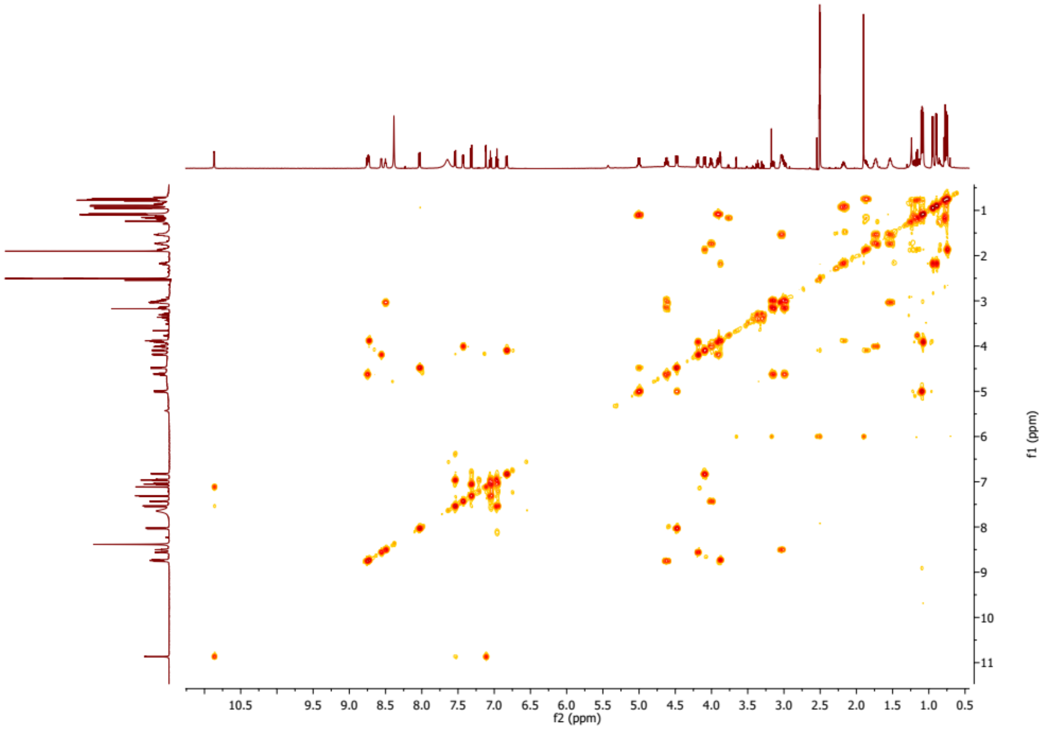


**Figure S48.** [1]H-[1]H COSY spectrum of compound **26**.

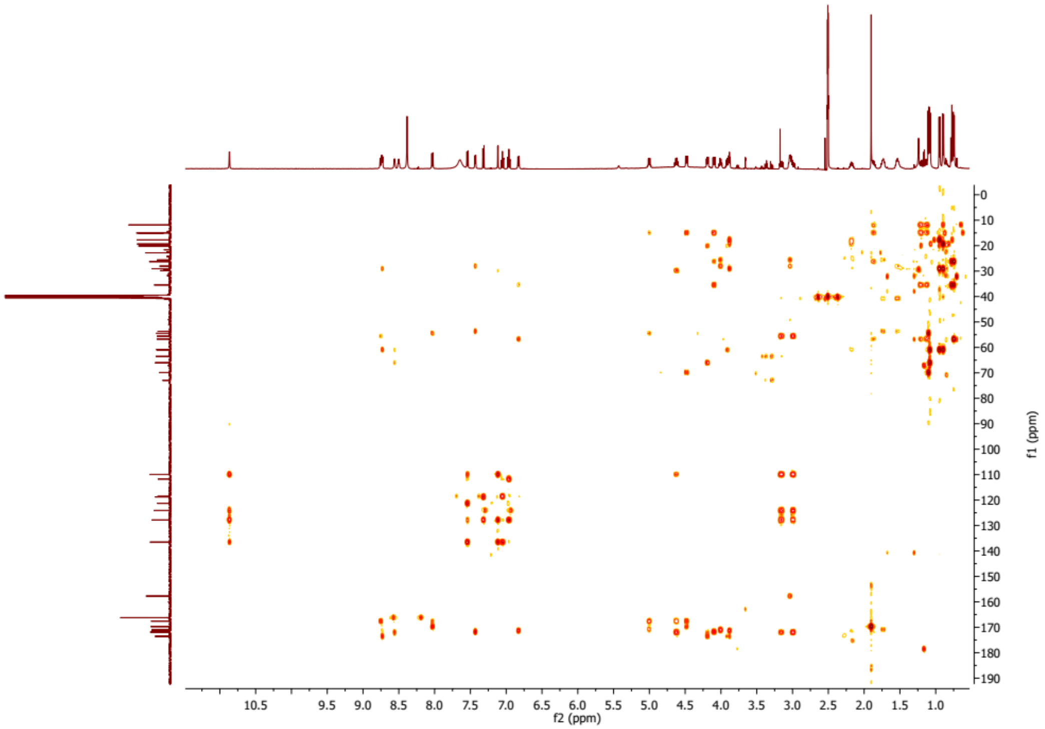


**Figure S49.** HMBC spectrum of compound **26**.

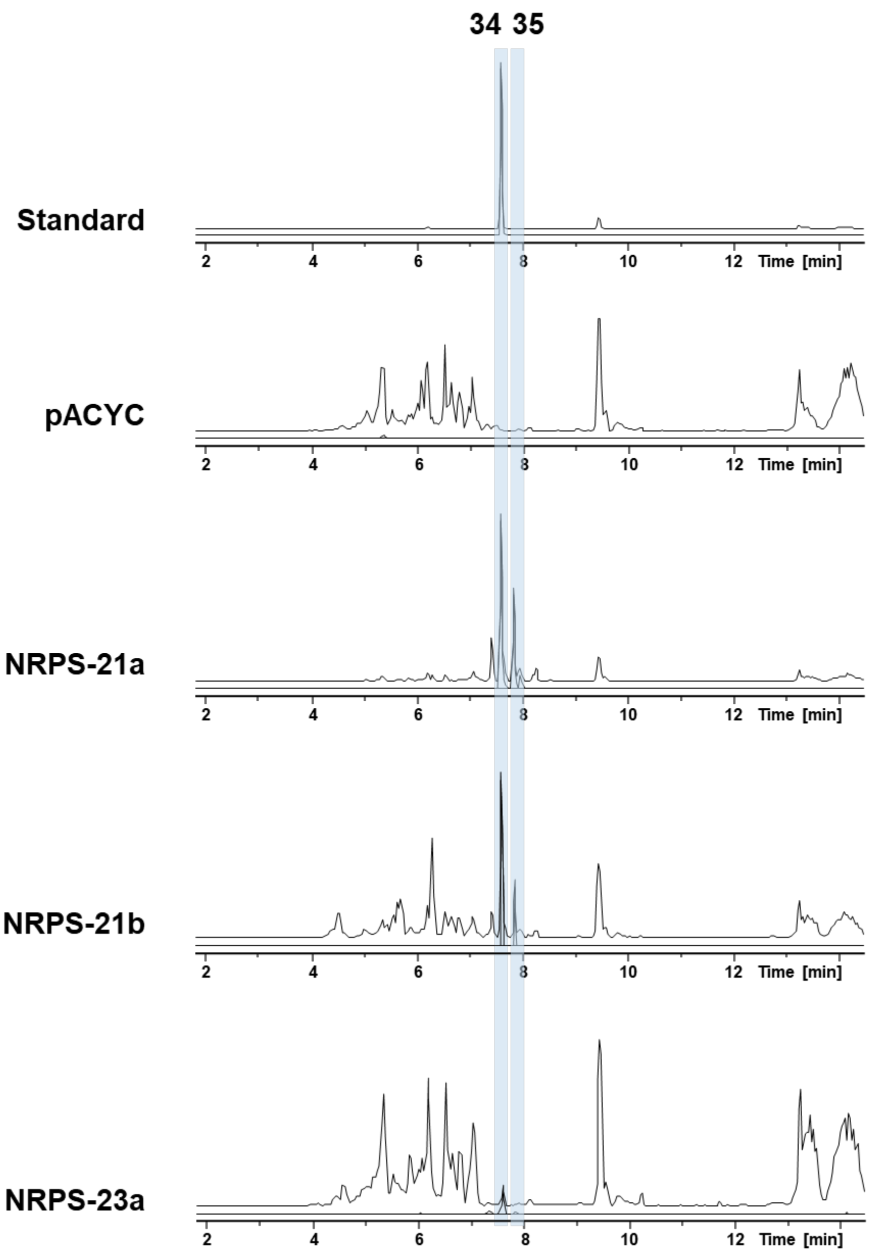


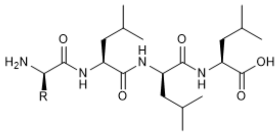


| Products | R |
|---|---|
| **34** | *i*Pr |
| **35** | *i*Bu |

**Figure S50**. HPLC/MS data refers to Figure 4 (NRPS-21 and -23) of compound **34** and **35** produced in *E. coli* DH10B::*mtaA*. Base Peak Chromatogram (BPC, top) and Extracted Ion Chromatogram (EIC, below) of **34** (*m/z* $[M+H]^+$ = 457.34) and **35** (*m/z* $[M+H]^+$ = 471.35). Chromatograms were compared to an empty vector control and a synthetic standard of compound **34** (*m/z* $[M+H]^+$ = 457.34).

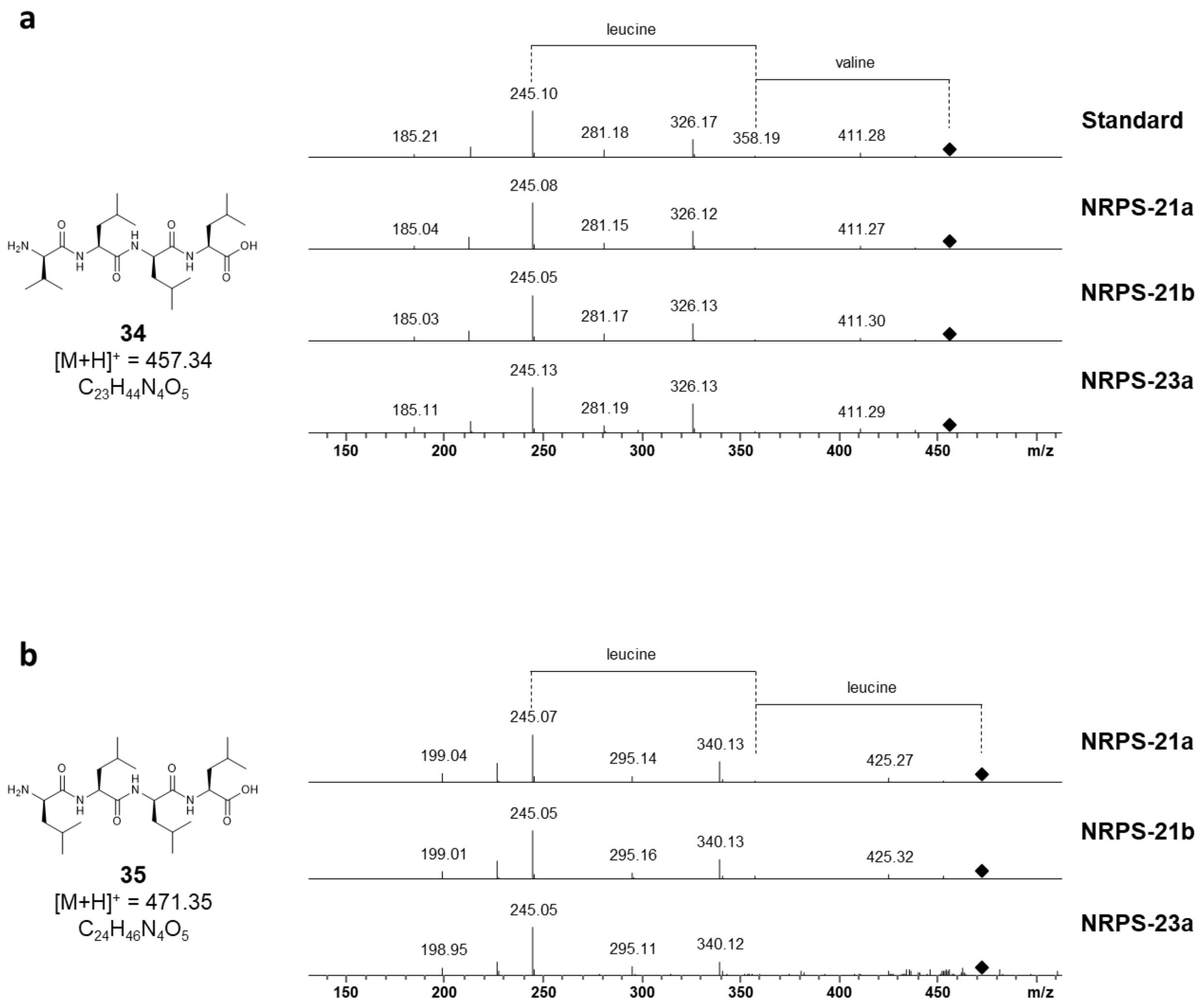


**Figure S51. a**) HPLC/MS data refers to Figure 4 (NRPS-21 and -23) of compound **34** produced in *E. coli* DH10B::*mtaA*. $MS^2$ and amino acid fragmentation of compound **34** produced by NRPS-21 and -23 compared to a synthetic standard of compound **34**. **b**) HPLC/MS data refers to Figure 4 (NRPS-21 and -23) of compound **35** produced in *E. coli* DH10B::*mtaA*. $MS^2$ and amino acid fragmentation of compound **35** produced by NRPS-21 and -23.

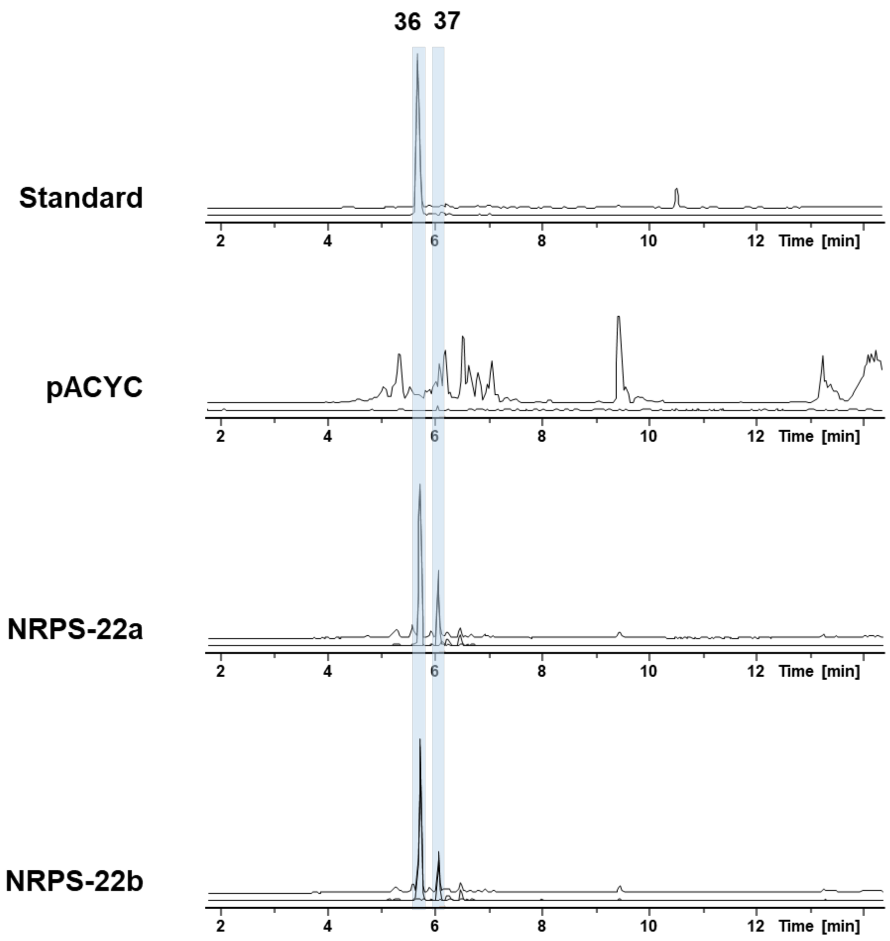


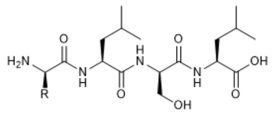


| Products | R |
|---|---|
| **36** | *i*Pr |
| **37** | *i*Bu |

**Figure S52.** HPLC/MS data refers to Figure 4 (NRPS-22) of compound **36** and **37** produced in *E. coli* DH10B::*mtaA*. Base Peak Chromatogram (BPC, top) and Extracted Ion Chromatogram (EIC, below) of **36** (*m/z* $[M+H]^+$ = 431.28) and **37** (*m/z* $[M+H]^+$ = 445.30). Chromatograms were compared to an empty vector control and a synthetic standard of compound **36** (*m/z* $[M+H]^+$ = 431.28).

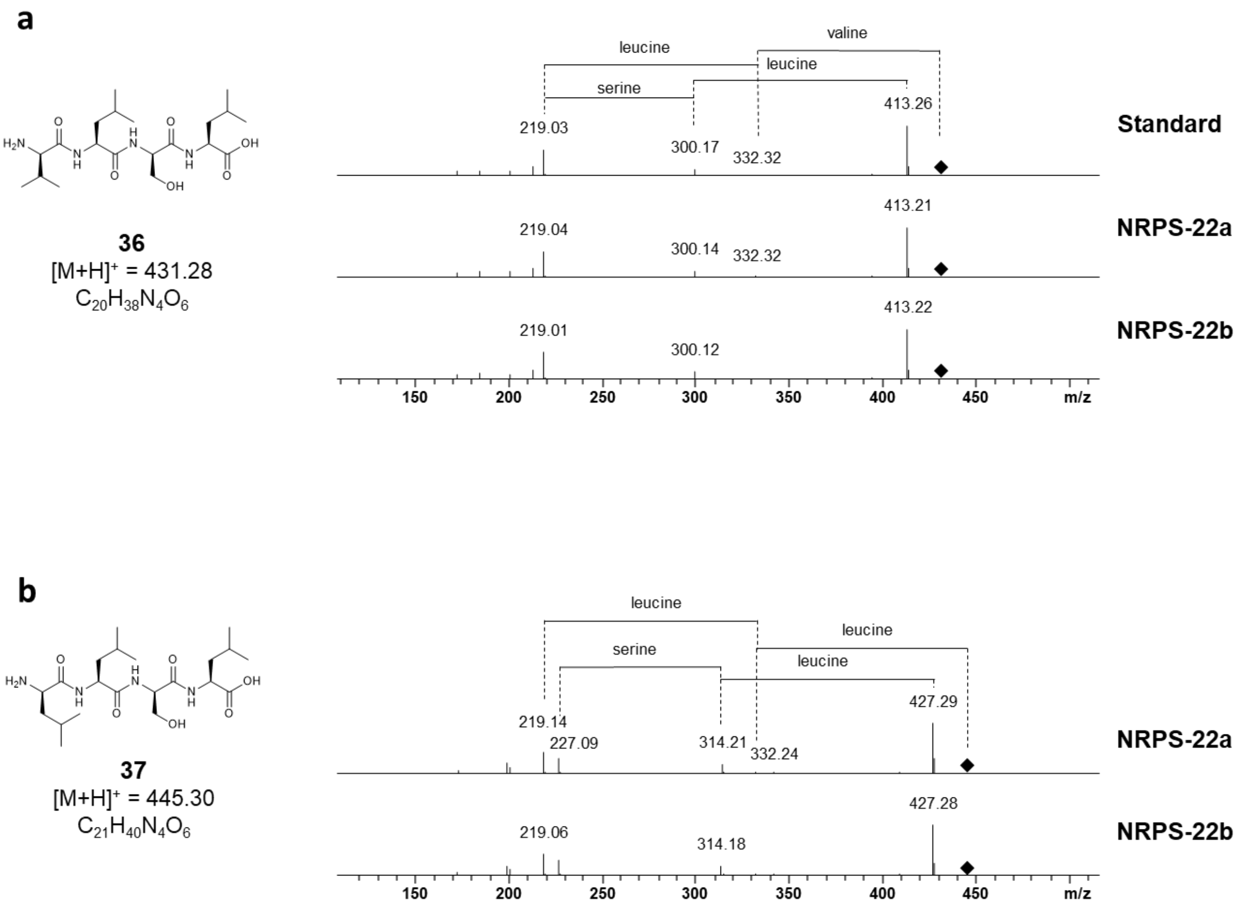


**Figure S53. a**) HPLC/MS data refers to Figure 4 (NRPS-22) of compound **36** produced in *E. coli* DH10B::*mtaA*. $MS^2$ and amino acid fragmentation of compound **36** produced by NRPS-22 compared to a synthetic standard of compound **36**. **b**) HPLC/MS data refers to Figure 4 (NRPS-22) of compound **37** produced in *E. coli* DH10B::*mtaA*. $MS^2$ and amino acid fragmentation of compound **37** produced by NRPS-22.

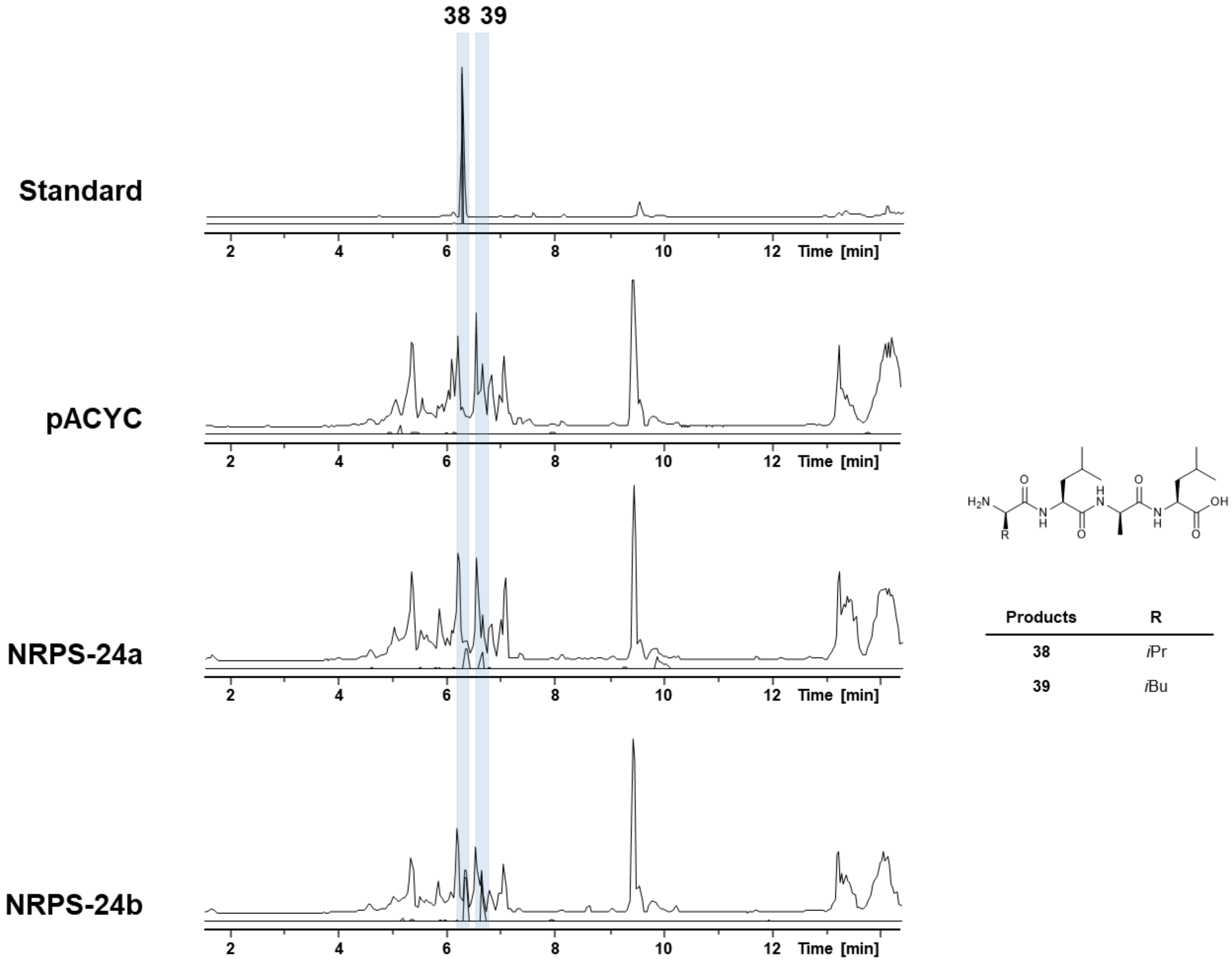


**Figure S54.** HPLC/MS data refers to Figure 4 (NRPS-24) of compound **38** and **39** produced in *E. coli* DH10B::*mtaA*. Base Peak Chromatogram (BPC, top) and Extracted Ion Chromatogram (EIC, below) of **38** (*m/z* $[M+H]^+$ = 415.29) and **39** (*m/z* $[M+H]^+$ = 429.31). Chromatograms were compared to an empty vector control and a synthetic standard of compound **38** (*m/z* $[M+H]^+$ = 415.29).

a

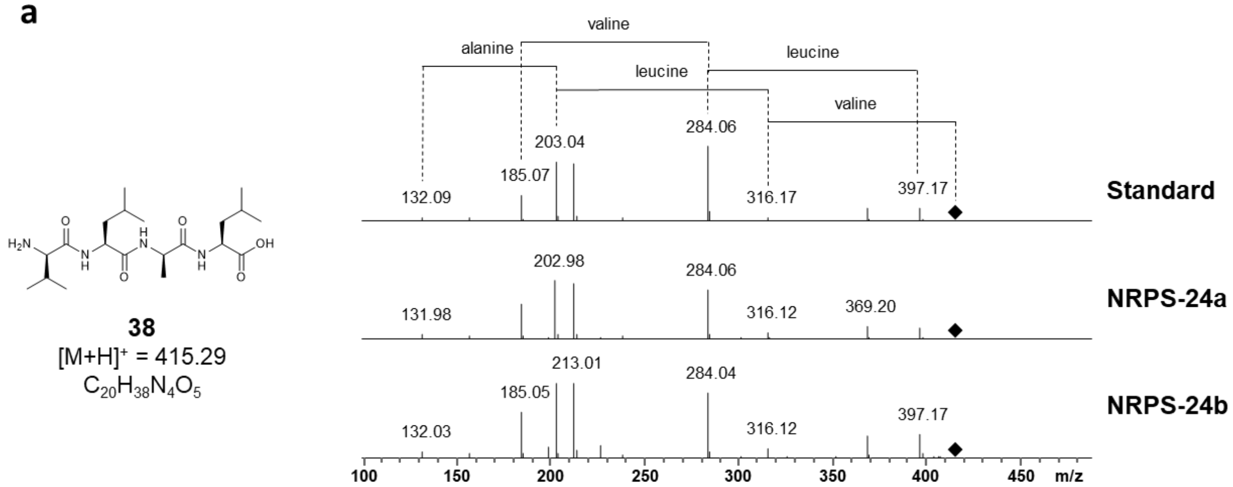


b

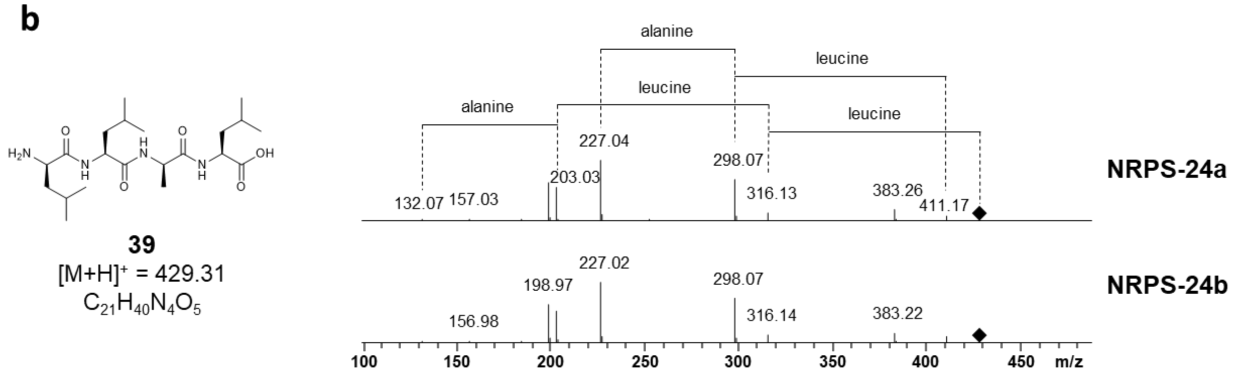


**Figure S55. a**) HPLC/MS data refers to Figure 4 (NRPS-24) of compound **38** produced in *E. coli* DH10B::*mtaA*. $MS^2$ and amino acid fragmentation of compound **38** produced by NRPS-24 compared to a synthetic standard of compound **38**. **b**) HPLC/MS data refers to Figure 4 (NRPS-24) of compound **39** produced in *E. coli* DH10B::*mtaA*. $MS^2$ and amino acid fragmentation of compound **39** produced by NRPS-24.

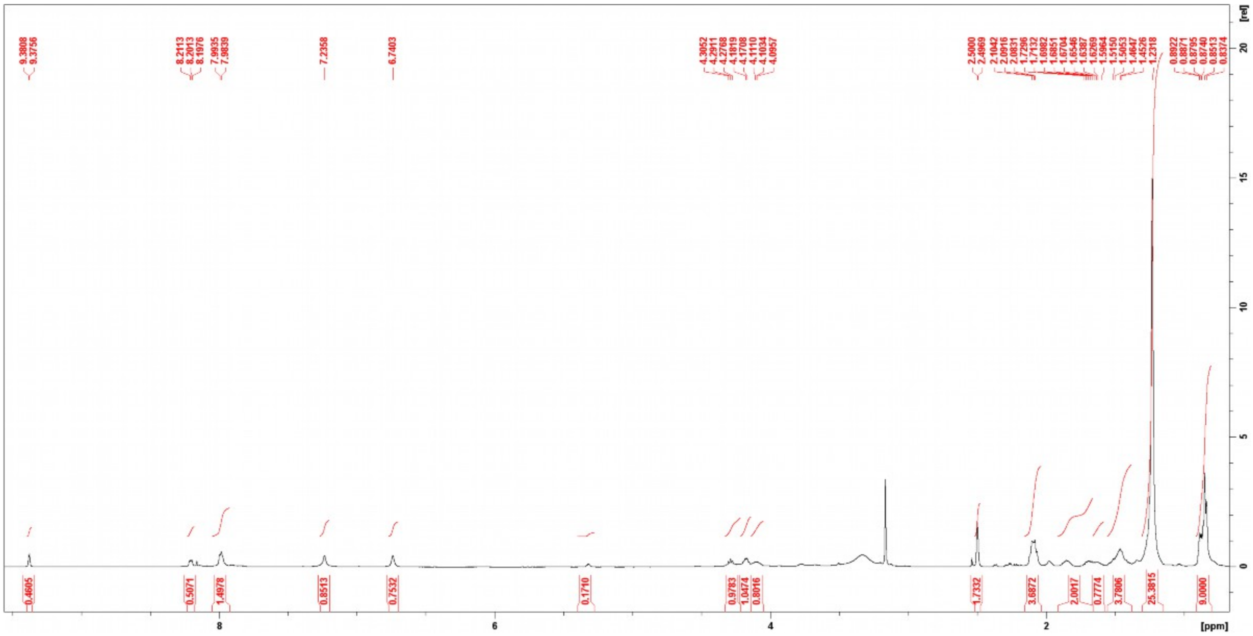

**Figure S56.** $^{1}H$ NMR (500 MHz, DMSO-$d_6$) spectrum compound **41**.

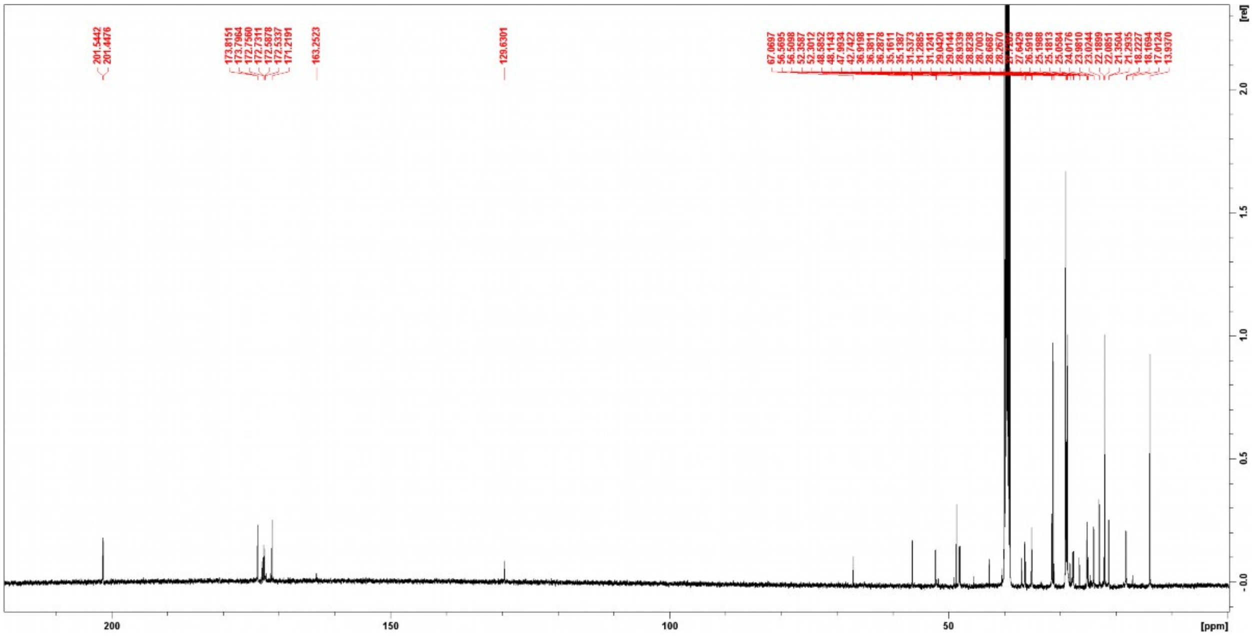

**Figure S57.** $^{13}C$ NMR (125 MHz, DMSO-$d_6$) spectrum compound **41**.

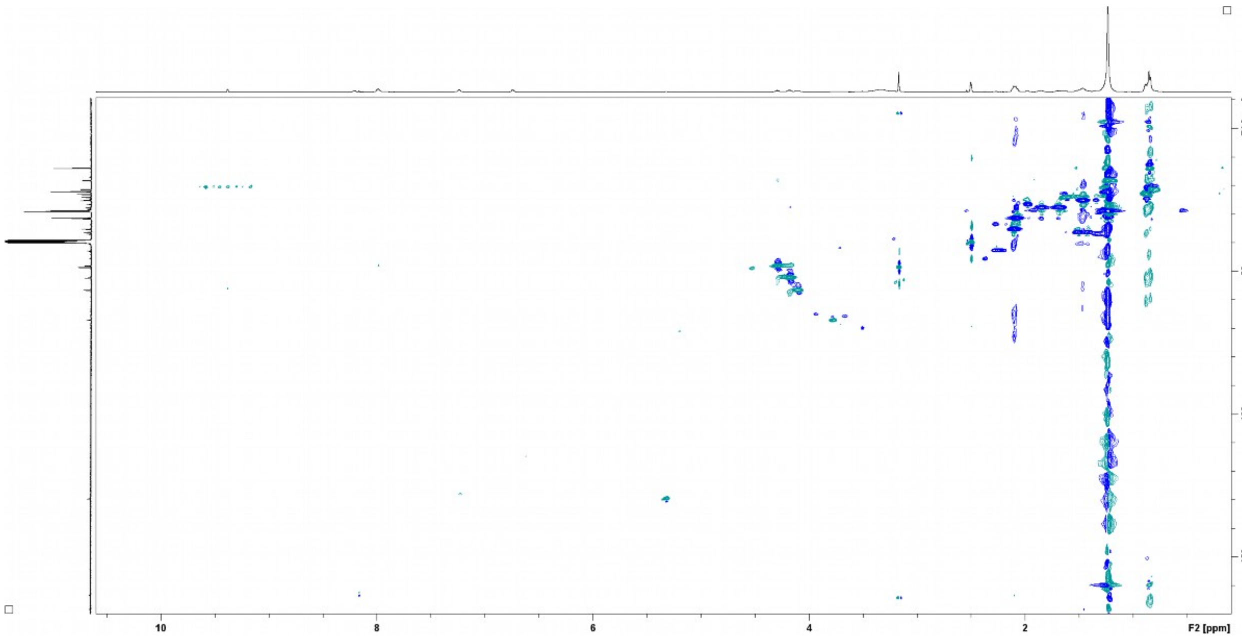


**Figure S58.** HSQC (DMSO-d6) spectrum of compound **41**.

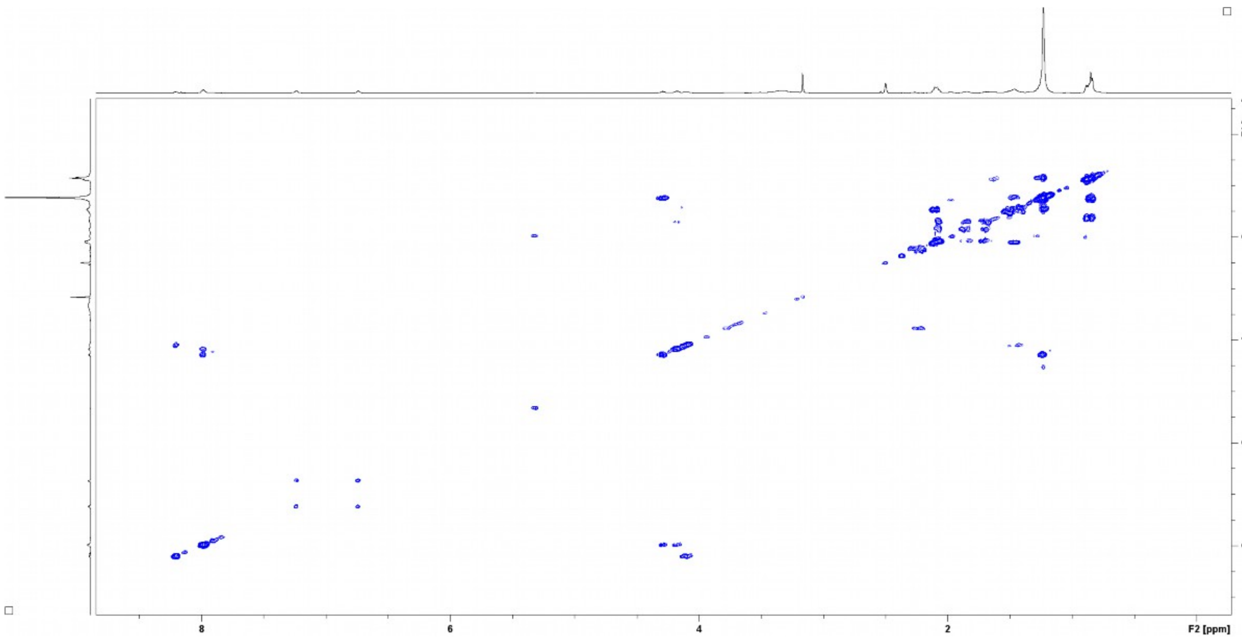


**Figure S59.** $^{1}H$-$^{1}H$ COSY (DMSO-d6) spectrum compound **41**.

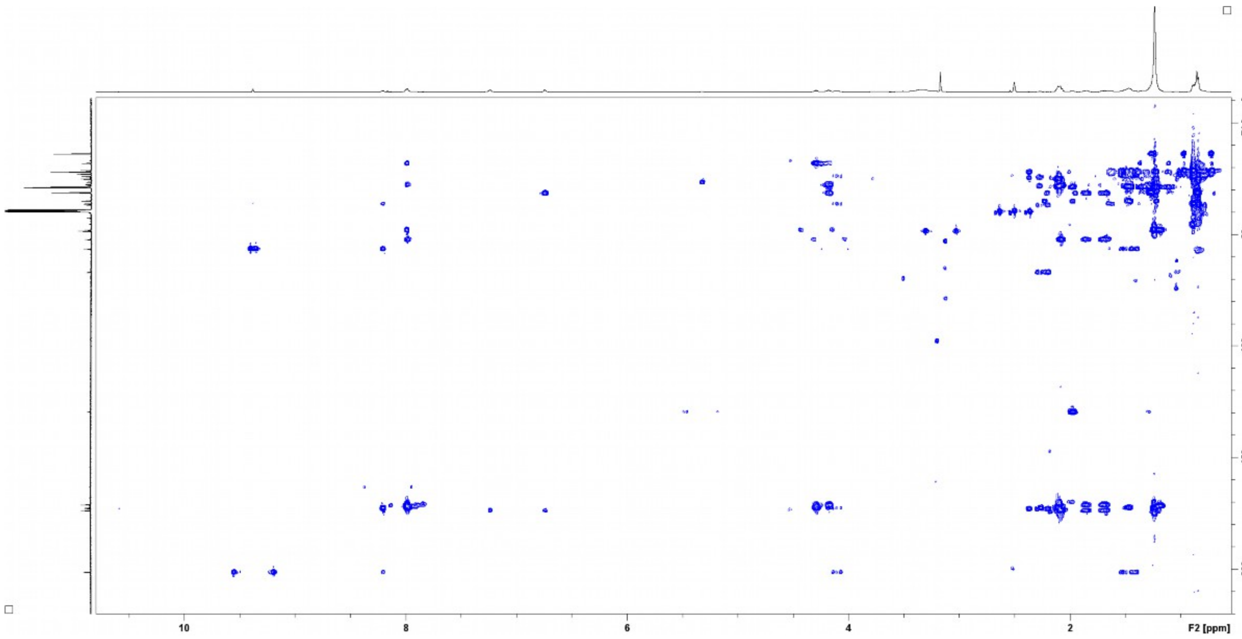


**Figure S60.** HMBC (DMSO-d6) spectrum compound **41**.

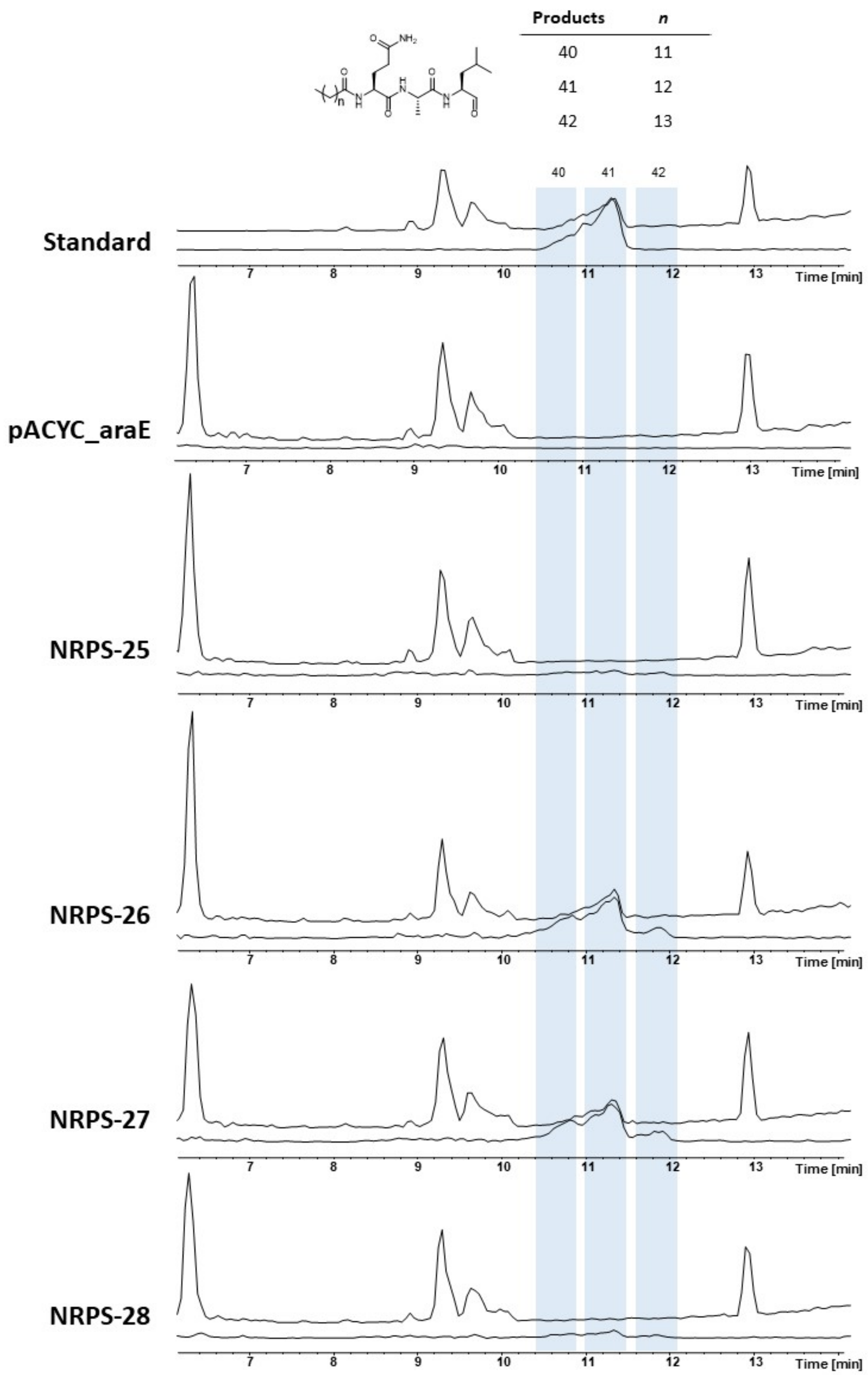


**Figure 61.** HPLC/MS data refers to Figure 5 (NRPS-25-28) of compound **40**, **41** and **42** produced in *E. coli* DH10B::*mtaA*. Base Peak Chromatogram (BPC, top) and Extracted Ion Chromatogram (EIC, below) of **40** (*m/z* $[M+H]^+$ = 511.38), **41** (*m/z* $[M+H]^+$ = 525.40), **42** (*m/z* $[M+H]^+$ = 539.41). Chromatograms were compared to an empty vector control and a purified compound **42** standard (*m/z* $[M+H]^+$ = 525.40).

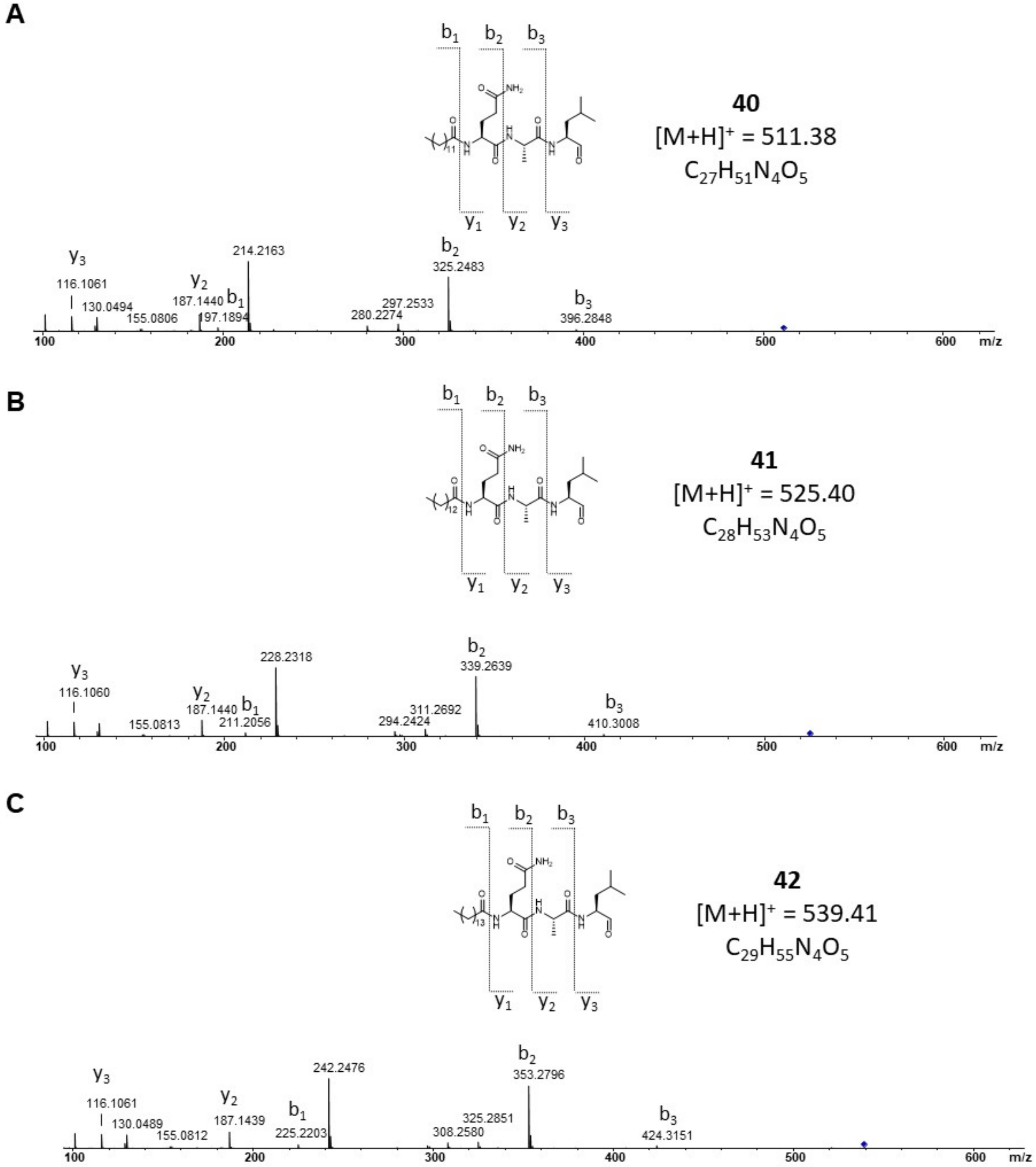


**Figure S62.** HPLC/MS data refers to Figure 5 (NRPS-26) of compound **40** (A), **41** (B) and **42** (C) produced in *E. coli* DH10B::*mtaA*. Comparison of $MS^2$ spectra.

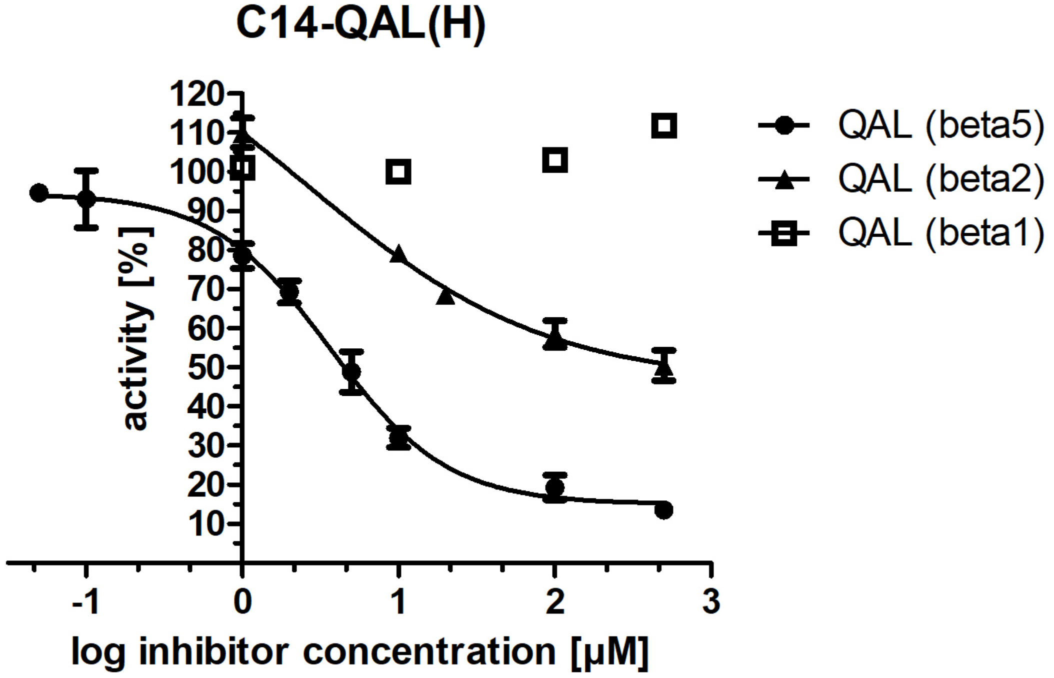


**Figure S63.** $IC_{50}$ determination of compound **41** (termed as C14-QAL(H)) for subunits beta1, -2 and -5 of yeast 20S proteasome (yCP).

**References**

1 Bode, E. *et al.* Promoter Activation in Δ hfq Mutants as an Efficient Tool for Specialized Metabolite Production Enabling Direct Bioactivity Testing. *Angewandte Chemie* **131**, 19133-19139, doi:10.1002/ange.201910563 (2019).

2 Bozhueyuek, K. A. J., Watzel, J., Abbood, N. & Bode, H. B. Synthetic Zippers as an Enabling Tool for Engineering of Non-Ribosomal Peptide Synthetases**. *Angewandte Chemie* **133**, 17672-17679, doi:10.1002/ange.202102859 (2021).

3 Gallastegui, N. & Groll, M. Analysing properties of proteasome inhibitors using kinetic and X-ray crystallographic studies. *Methods Mol Biol* **832**, 373-390, doi:10.1007/978-1-61779-474-2_26 (2012).

4 Groll, M. & Huber, R. Purification, crystallization, and X-ray analysis of the yeast 20S proteasome. *Methods Enzymol* **398**, 329-336, doi:10.1016/S0076-6879(05)98027-0 (2005).

5 Kabsch, W. *XDS*. *Acta Crystallographica Section D Biological Crystallography* **66**, 125-132, doi:10.1107/s0907444909047337 (2010).

6 Schneider, T. R. & Sheldrick, G. M. Substructure solution with SHELXD. *Acta Crystallogr D Biol Crystallogr* **58**, 1772-1779, doi:10.1107/s0907444902011678 (2002).

7 Huber, E. M. *et al.* A unified mechanism for proteolysis and autocatalytic activation in the 20S proteasome. *Nature Communications* **7**, 10900, doi:10.1038/ncomms10900 (2016).

8 Emsley, P., Lohkamp, B., Scott, W. G. & Cowtan, K. Features and development of *Coot*. *Acta Crystallographica Section D Biological Crystallography* **66**, 486-501, doi:10.1107/s0907444910007493 (2010).

9 Edgar, R. C. MUSCLE: multiple sequence alignment with high accuracy and high throughput. *Nucleic Acids Res* **32**, 1792-1797, doi:10.1093/nar/gkh340 (2004).

10 Capella-Gutierrez, S., Silla-Martinez, J. M. & Gabaldon, T. trimAl: a tool for automated alignment trimming in large-scale phylogenetic analyses. *Bioinformatics* **25**, 1972-1973, doi:10.1093/bioinformatics/btp348 (2009).

11 Jones, D. T., Taylor, W. R. & Thornton, J. M. The rapid generation of mutation data matrices from protein sequences. *Comput Appl Biosci* **8**, 275-282, doi:10.1093/bioinformatics/8.3.275 (1992).
12 Maddison, W. & Maddison, D. R. Mesquite: A modular system for evolutionary analysis. *Evolution* **62**, 1103-1108 (2008).
13 Revell, L. J. phytools: an R package for phylogenetic comparative biology (and other things). *Methods in Ecology and Evolution* **3**, 217-223, doi:10.1111/j.2041-210X.2011.00169.x (2012).
14 Durfee, T. *et al.* The Complete Genome Sequence of *Escherichia coli* DH10B: Insights into the Biology of a Laboratory Workhorse. *Journal of Bacteriology* **190**, 2597-2606, doi:10.1128/jb.01695-07 (2008).
15 Schimming, O., Fleischhacker, F., Nollmann, F. I. & Bode, H. B. Yeast Homologous Recombination Cloning Leading to the Novel Peptides Ambactin and Xenolindicin. *ChemBioChem* **15**, 1290-1294, doi:10.1002/cbic.201402065 (2014).
16 Goldman, B. S. *et al.* Evolution of sensory complexity recorded in a myxobacterial genome. *Proceedings of the National Academy of Sciences* **103**, 15200-15205, doi:10.1073/pnas.0607335103 (2006).
17 Krug, D. *et al.* Discovering the hidden secondary metabolome of Myxococcus xanthus: a study of intraspecific diversity. *Appl Environ Microbiol* **74**, 3058-3068, doi:10.1128/AEM.02863-07 (2008).
18 Kissoyan, K. A. B. *et al.* Natural C. elegans Microbiota Protects against Infection via Production of a Cyclic Lipopeptide of the Viscosin Group. *Current Biology* **29**, 1030-1037.e1035, doi:10.1016/j.cub.2019.01.050 (2019).
19 Petersen, L. M. & Tisa, L. S. Influence of temperature on the physiology and virulence of the insect pathogen Serratia sp. Strain SCBI. *Appl Environ Microbiol* **78**, 8840-8844, doi:10.1128/AEM.02580-12 (2012).
20 Joyce, S. A. & Clarke, D. J. A hexA homologue from Photorhabdus regulates pathogenicity, symbiosis and phenotypic variation. *Molecular Microbiology* **47**, 1445-1457, doi:10.1046/j.1365-2958.2003.03389.x (2003).
21 Tobias, N. J. *et al.* Natural product diversity associated with the nematode symbionts Photorhabdus and Xenorhabdus. *Nature Microbiology* **2**, 1676-1685, doi:10.1038/s41564-017-0039-9 (2017).
22 Behsaz, B. *et al.* Integrating genomics and metabolomics for scalable non-ribosomal peptide discovery. *Nature Communications* **12**, doi:10.1038/s41467-021-23502-4 (2021).
23 Lorenzen, W., Ahrendt, T., Bozhüyük, K. A. J. & Bode, H. B. A multifunctional enzyme is involved in bacterial ether lipid biosynthesis. *Nature Chemical Biology* **10**, 425-427, doi:10.1038/nchembio.1526 (2014).
24 Kegler, C. & Bode, H. B. Artificial Splitting of a Non-Ribosomal Peptide Synthetase by Inserting Natural Docking Domains. *Angewandte Chemie* **132**, 13565-13569, doi:10.1002/ange.201915989 (2020).
25 Fuchs, S. W., Proschak, A., Jaskolla, T. W., Karas, M. & Bode, H. B. Structure elucidation and biosynthesis of lysine-rich cyclic peptides in Xenorhabdus nematophila. *Organic & Biomolecular Chemistry* **9**, 3130, doi:10.1039/c1ob05097d (2011).
26 Vo, T. D., Spahn, C., Heilemann, M. & Bode, H. B. Microbial Cationic Peptides as a Natural Defense Mechanism against Insect Antimicrobial Peptides. *ACS Chemical Biology* **16**, 447-451, doi:10.1021/acschembio.0c00794 (2021).
27 Sievers, F. & Higgins, D. G. Clustal Omega for making accurate alignments of many protein sequences. *Protein Sci* **27**, 135-145, doi:10.1002/pro.3290 (2018).
28 Blin, K. *et al.* antiSMASH 6.0: improving cluster detection and comparison capabilities. *Nucleic Acids Res* **49**, W29-W35, doi:10.1093/nar/gkab335 (2021).
29 Niehs, S. P. *et al.* Genome Mining Reveals Endopyrroles from a Nonribosomal Peptide Assembly Line Triggered in Fungal–Bacterial Symbiosis. *ACS Chemical Biology* **14**, 1811-1818, doi:10.1021/acschembio.9b00406 (2019).
30 Wang, X. *et al.* Discovery of recombinases enables genome mining of cryptic biosynthetic gene clusters in Burkholderiales species. *Proc Natl Acad Sci U S A* **115**, E4255-E4263, doi:10.1073/pnas.1720941115 (2018).
31 Isogai, A. *et al.* Structural Analysis of New Syringopeptins by Tandem Mass Spectrometry. *Bioscience, Biotechnology, and Biochemistry* **59**, 1374-1376, doi:10.1271/bbb.59.1374 (1995).
